# Pan-genome analyses of 226 finger millet-infecting *Magnaporthe oryzae* strains from eastern Africa

**DOI:** 10.1101/2025.10.14.682320

**Authors:** Peng Qi, Bochra A. Bahri, Jie Zhu, Hallie C. Wright, Kathryn A. Prado, Yunus Sahin, Hao Wang, Dong Won Kim, Brandon Mangum, Jane Grimwood, Jerry Jenkins, Joseph M. Atemia, Taiwo A Shittu, Sreenivasaprasad Muthumeenakshi, Thomas H. Pendergast, Tesfaye Alemu Tenkegna, Mathews M. Dida, Justin H. Ringo, John Takan, Kassahun Tesfaye, Surapareddy Sreenivasaprasad, Ki-Tae Kim, Yong-Hwan Lee, Santie de Villiers, Chang Hyun Khang, Katrien M. Devos

## Abstract

Blast disease, caused by the filamentous ascomycete *Magnaporthe oryzae*, is the main biotic constraint to finger millet production in eastern Africa. *M. oryzae* is a pathogen on many grass species, but the high host-specificity of blast isolates underscores the need to study pathogen diversity and virulence at the host level. Here, we fill a void on knowledge on finger millet-infecting strains of *M. oryzae* by sequencing 226 isolates pure-cultured from infected tissues, mainly peduncles and panicles, sampled from field-grown finger millet accessions across Ethiopia, Kenya, Tanzania and Uganda. Phylogenetic analysis showed that eastern African isolates are genetically distinct from Asian isolates, and differentiated into two groups potentially driven by climate variables. One group was dominated by isolates from Kenya and Uganda (KU group), and the other by isolates from Ethiopia and Tanzania (ET group). Analysis of the portfolio of 887 predicted effector genes showed that the KU group had significantly fewer effectors, concomitant with higher virulence levels on the two finger millet accessions tested.

We demonstrate that homologous recombination between transposable elements, leading to genomic deletions, plays a key role in gene removal. Transcript profiling of fungal genes in compatible and incompatible interactions revealed upregulation in the incompatible interaction as a characteristic of an estimated 30% of effector genes, including seven of the eight homologs of known avirulence genes from rice-infecting *M. oryzae* identified in our study. Prioritization of these effectors for functional validation will pave the way for identifying cognate resistance genes and improving finger millet for resistance to blast disease.

## Introduction

Finger millet (*Eleusine coracana* L. Gaertn), a critical food security crop in eastern Africa and India, faces a significant threat from blast disease caused by the fungus *Magnaporthe* (synonym *Pyricularia*) *oryzae* (JEEVAN *et al*. 2021). The fungus can infect any tissue, but yield losses are most severe upon infection of the peduncle (neck blast) and panicle (head blast), and can vary from less than 10% to up to 90% depending on the environmental conditions (RAO 1990; LULE *et al*. 2014; ODEPH *et al*. 2020). Blast disease can be managed with fungicides (PRAJAPATI *et al*. 2013; PATRO *et al*. 2020), but chemical pest management strategies are not within the realm of smallholder farmers growing finger millet as a subsistence crop under minimal inputs. Further, fungicide use has adverse effects on the environment (ZUBROD *et al*. 2019; PIMENTÃO *et al*. 2024) and carries a risk of pathogens acquiring resistance (SUZUKI *et al*. 2010; CASTROAGUDÍN *et al*. 2015; POLONI *et al*. 2021). Sustainable disease control provided by durable host resistance, however, requires an advanced understanding of the plant-pathogen dynamics.

Effectors are pivotal components of the interplay between a pathogen and its host, and can trigger both susceptibility through suppression of pathogen-associated molecular patterns (PAMP)-triggered immunity and resistance through interaction with plant host resistance (*R*) genes (JONES AND DANGL 2006; NGOU *et al*. 2022; JONES *et al*. 2024). Changes in either the fungal effector or host resistance gene repertoire can cause a switch from a compatible (susceptible) to an incompatible (resistance) interaction and *vice versa* (BIAŁAS *et al*. 2018; SÁNCHEZ-VALLET *et al*. 2018; LE NAOUR-VERNET *et al*. 2023; GLADIEUX *et al*. 2024). The result is a continuous cycle of effector gene mutation or loss by the fungus to evade host resistance mechanisms, and adaptative deployment of new resistance mechanisms by the host.

*M. oryzae* is a filamentous ascomycete capable of infecting a broad range of grasses. Pathotypes, however, have a high degree of host-specificity. Despite the importance of finger millet as a staple food in the developing world, and the large toll blast disease takes on finger millet production, information on the *E. coracana* pathotype (*MoE*) and its interactions at the genomic level are largely lacking. Here, we address this knowledge gap by generating a reference genome for an Ethiopian *MoE* blast strain, and short-read assemblies for an additional 225 isolates originating from infected tissues collected across eastern Africa, the center of origin and diversity of finger millet (DEVOS *et al*. 2023). Previously, two genetic groups, largely separating Asian from African isolates, were identified within the *MoE* pathotype based on various criteria including transposable elements (DOBINSON *et al*. 1993; TANAKA *et al*. 2009), whole genome single nucleotide polymorphisms (SNPs) (GLADIEUX *et al*. 2018a), and differential presence of the effector genes *PWL1* and *PWL2* (ASUKE *et al*. 2020; MASAKI *et al*. 2023). We show that *MoE* isolates from eastern Africa further diverged into two genetic groups, potentially associated with climate zones. The two African *MoE* groups differ from each other and the Asian *MoE* group in their genetic make-up, effector gene repertoire and virulence level. We further demonstrate that loss of chromosomal regions, and hence effector genes, occurs predominantly through homologous recombination between transposable elements. The presented insights derived from the *M. oryzae* pan-genome combined with transcriptome analyses at early infection time points will be invaluable in the quest to harness sustainable, effector-targeted resistance strategies for safeguarding finger millet and other vital crops against the threat of blast disease.

## Results

### The pan-genome of finger millet-infecting Magnaporthe oryzae isolates from eastern Africa

A total of 207 *M. oryzae* strains were isolated from infected finger millet inflorescences collected in Kenya (K; n=41), Uganda (U; n=56), Tanzania (T; n=54) and Ethiopia (E; n=56) during the period 2015-2017 (**Figure 1; Table S1; Data S1**). The 207 strains plus 10 Kenyan and nine Ugandan strains isolated during the period 2000-2002 (TAKAN *et al*. 2012) were Illumina sequenced (2×150 bp) to an average depth of 94.4x (range 15.9x – 223.1x) (**Data S1**; PRJNA1279063). One isolate, E2, originating from Ethiopia, also underwent PacBio sequencing (235.47x; average read length of 7855 bp; **Table S2**) to generate a reference genome assembly of 44.35 Mb organized into seven pseudomolecules (contig N/L50 of 4/5.8 Mb; see **Table S3** for summary statistics; Genbank accession number JBPPCE000000000). The E2 assembly comprised 97.7% of Ascomycota Benchmarking Universal Single-Copy Orthologs (BUSCO) (SIMÃO *et al*. 2015) (**Figure S1**). Illumina-based contigs were generated *de novo* for the other 225 strains, and contigs were ordered based on the E2 assembly. These short-read assemblies have an average contig N50 of 336.1 kb (range 51.9 kb – 832.9 kb) and a total average assembly size of 41.0 Mb (range 37.2 – 49.1) (**Data S1**).

**Figure 1.**
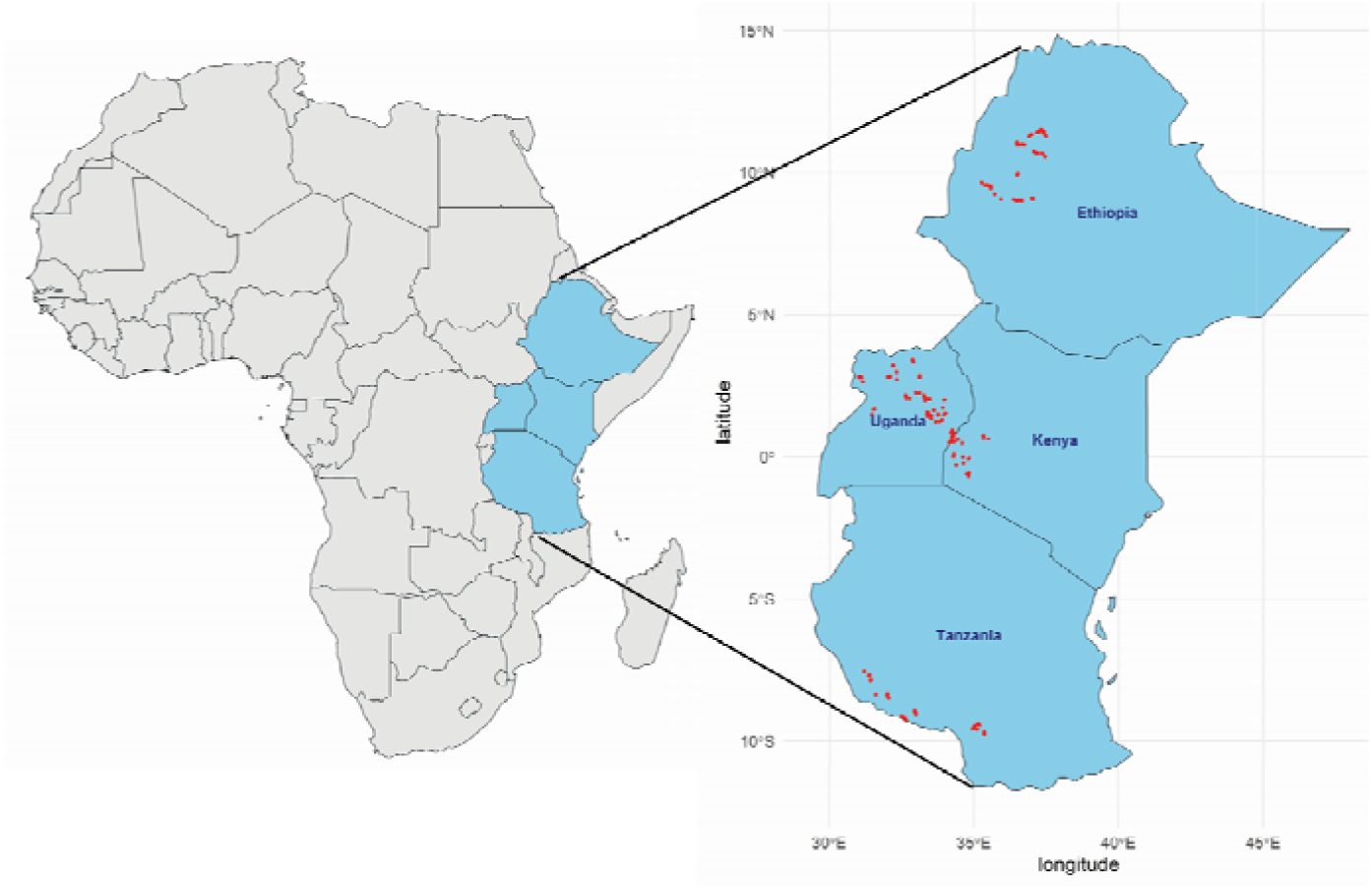
Map showing the collection sites (red dots) in eastern Africa of the M. oryzae strains sequenced

A total of 13,243 genes with a cDNA length ≥ 200 bp were annotated in E2. The gene set comprises 97.0% of Ascomycota BUSCO genes (**Figure S1**). An additional 588 non-redundant genes were *de novo* annotated across the 225 short-read assemblies. An estimated 11.5% of annotated genes were missing from at least one strain. This conservative estimate does not take into account partially deleted or inactive genes. The repeat content of E2 was estimated at 11.3% (**Table S4**).

### Genetic relationships between M. oryzae isolates revealed three MoE genetic groups

A total of 198,454 genome-wide SNPs identified across the 226 eastern African *MoE* strains distinguished 166 haplotypes with between 1 and 20 members with an identity >99.9% (**Table S5)**. The haplotype with the largest membership comprises strains from three counties in Kenya and 11 counties in Uganda, indicating likely spread of clonal isolates. Phylogenetic analysis of representative strains for the 166 haplotypes using a subset of 15,838 SNP that maximized the genetic diversity showed two distinct genetic groups, with the majority of the Kenyan and Ugandan isolates clustering in the KU group, and most Ethiopian and Tanzanian isolates forming the ET group (**Figure 2)**. Population structure analyses identified the same two groups with 12 haplotypes (18 isolates) being classified as admixed (<90% membership to a single population) (**Figure S2**). Interestingly, in the 11 cases where isolates were obtained from both the peduncle and the panicle from the same plant, only two (18.2%) yielded highly similar (> 99.9%) isolates (**Table S6**). While field isolates can be composed of multiple strains with varying levels of divergence, we cannot ascertain whether isolate composition varies by tissue because only a single isolate was purified from each tissue.

**Figure 2.**
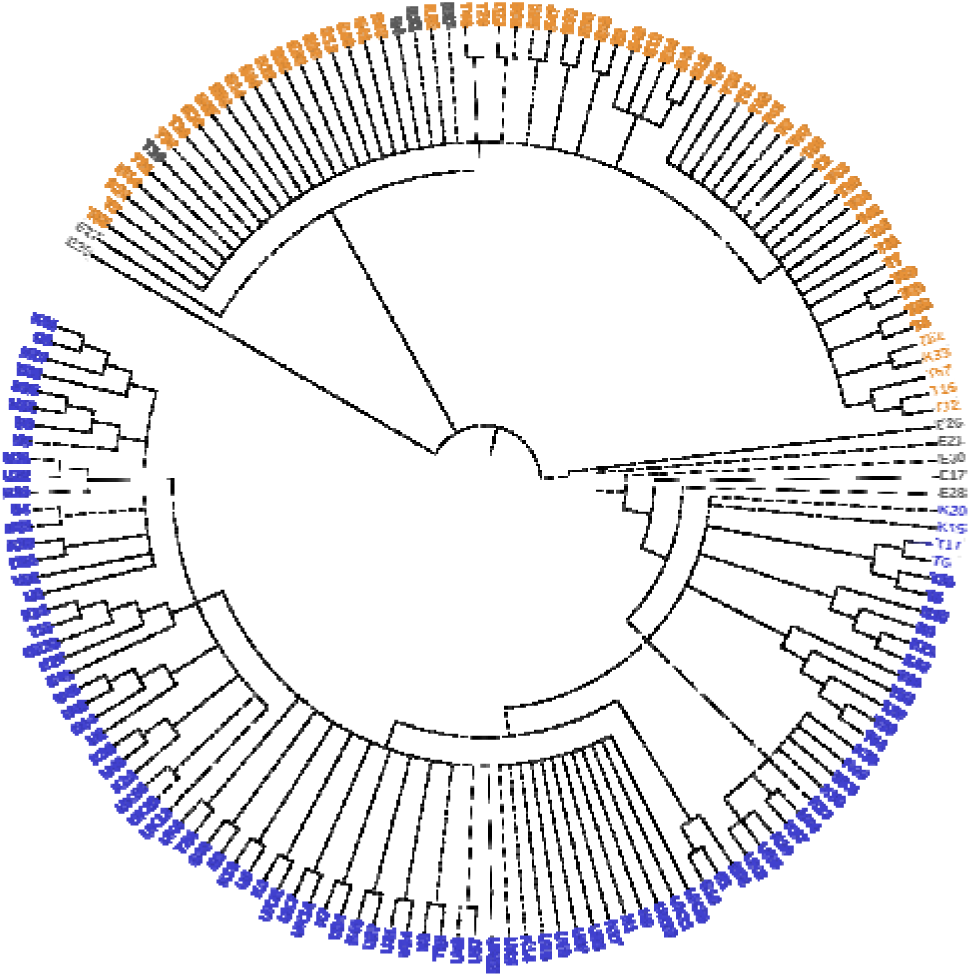
Maximum likelihood tree of representative isolates for 166 haplotypes identified from 226 MoE strains from eastern Africa. Strains are colored based on their membership to a genetic population (blue: KU population; orange: ET population; Grey: Admixed). Branches of the tree produced in <50% bootstrap replicates are collapsed.

The finger millet-infecting ET and KU groups are sister to a group of finger millet-infecting isolates from Asia (**Figure 3**). The majority of *E. indica* blast isolates (prefix Ei in **Figure 3**) are taxonomically most closely related to finger millet blast isolates from Kenya and Uganda, irrespective of their country and continent of origin. The geographic origin of *E. indica*, a worldwide weed, is not clear, but it was present in eastern Africa an estimated 1.3 million years ago (MYA) when it hybridized with an unknown species to form the wild tetraploid progenitor to finger millet, *E. coracana* subsp. *africana* (DEVOS *et al*. 2023). The close relationship of *E. indica* isolates from the Asian and South American continents with *E. coracana*-infecting isolates from Kenya and Uganda indicates dispersal of the *Eleusine* pathotype through infected seeds. An isolate from *Eragrostis curvula* (prefix Ecu; weeping lovegrass), a chloridoid species that can act as an alternative host to finger millet blast isolates lacking the avirulence genes *PWL1* and *PWL2* (MASAKI *et al*. 2023), was sister to the *Eleusine-*infecting isolates. In agreement with previous observations (GLADIEUX *et al*. 2018a), the chloridoid-infecting isolates were most closely related to blast isolates infecting species belonging to the subfamily Pooideae, including wheat, ryegrass and oats (**Figure 3**).

**Figure 3.**
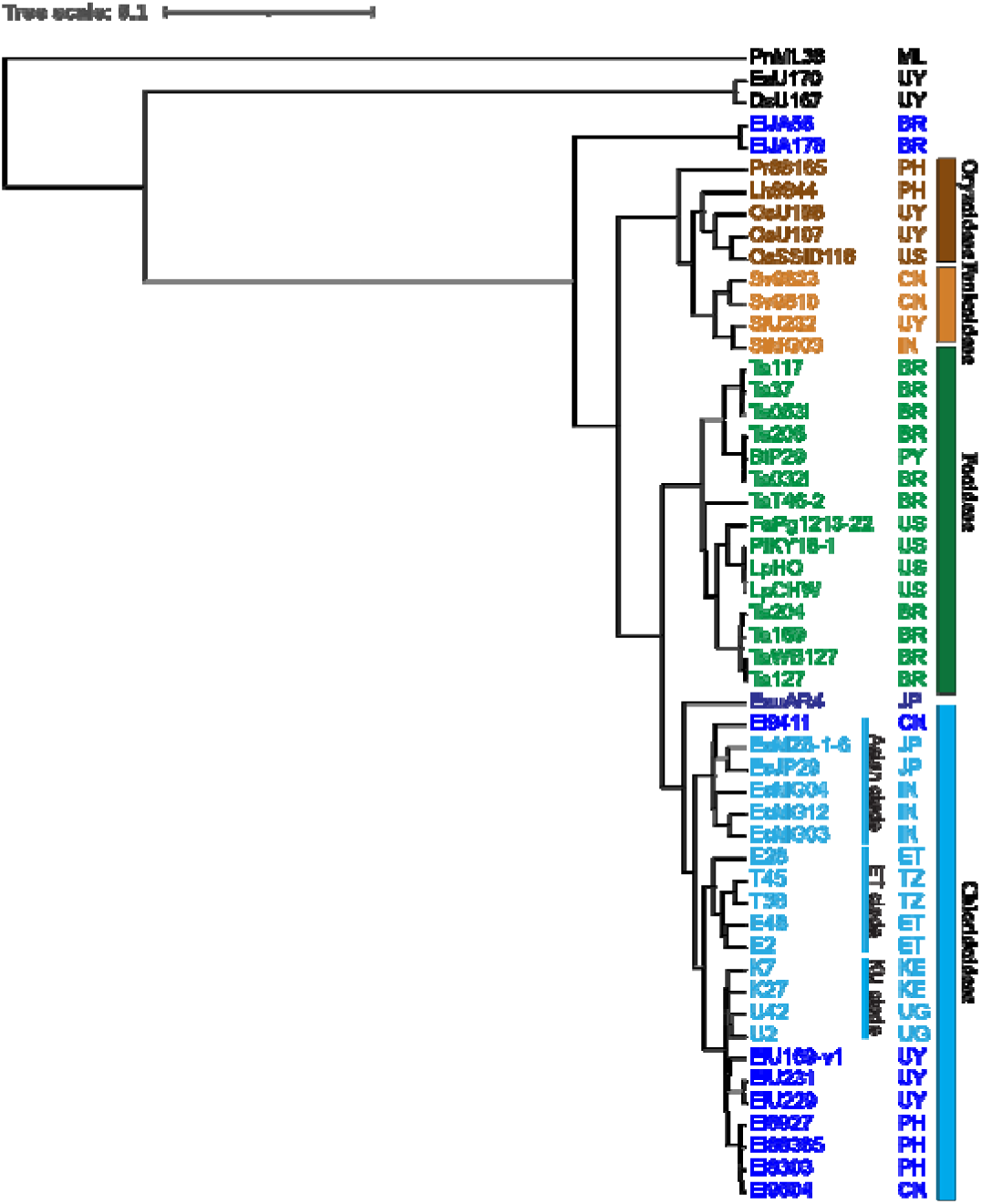
Dendrogram showing hierarchical clustering of M. oryzae isolates with different host specificity. Vertical bars indicate the subfamily taxonomy of the host species. Country of origin is indicated by their two-letter abbreviation (BR: Brazil; CN: China; ET: Ethiopia; IN: India; JP: Japan; KE: Kenya; ML: Mali; PH: Philippines; PY: Paraguay; TZ: Tanzania; UG: Uganda; US: United States; UY: Uruguay). Additional details, including host species and SRA accession number, are provided in **Table S7**.

### Differential expansion of TEs across the three groups of finger millet-infecting blast isolates

DOBINSON *et al*. (1993) uncovered the existence of two genetically distinct *MoE* pathotypes based on the presence of the long terminal repeat (LTR) retrotransposon *Grasshopper* (*Grh*). The *Grh* element was present in finger millet-infecting isolates from Japan, Nepal and India, and largely absent in isolates from Kenya and Uganda (DOBINSON *et al*. 1993; TAKAN *et al*. 2012). Repeat mining in the E2 (Ethiopia) and MZ5-1-6 (Japan) genome assemblies identified 45 and 76 full-length copies of *Grh*, respectively, making it the most prevalent full-length LTR-retrotransposon in both strains (**Table S8)**. *Grh* copy numbers estimated from the short-read assemblies (**Note S1**) averaged 4.5x for *MoE* isolates belonging to the KU group and 31.7x for ET isolates (**Table S9; Data S2**). These estimates demonstrate that the majority of Kenyan and Ugandan strains lack or only have few copies of the *Grh* element, as previously observed (TAKAN *et al*. 2012), while *Grh* underwent amplification in the ET group. All but one of the 121 full-length *Grh* elements in E2 and MZ5-1-6 have identical LTRs (**Data S3**), indicating they inserted in the last 80,000 years (threshold insertion time if LTRs were to differ by 1 bp). Greater than two-fold differences in copy number between the KU and ET subpopulations were also observed for the LTR-retrotransposon *Pyret* (17.7x in KU group *vs.* 39.7x in ET group) and the DNA transposon *Pot2* (10.6x in KU *vs.* 121x in ET) (**Figure S3; Table S9; Data S2**). Presence of the LTR-retrotransposon *Fosbury* (**Note S2**), on the other hand, largely differentiated Ethiopian (copy number of 34.4x) from Tanzanian strains (0.1x) within the ET population and from KU strains (0.5x) (**Figure S3; Data S2**).

### Comparative genomics of M. oryzae isolates with different host specificities

Synteny was high between isolates infecting finger millet (E2 and MZ5-1-6; *MoE* strains), wheat (B71; *MoT* strain), and rice (P131; *MoO* strain) (**Figure 4; Figure S4**). As expected, the number of interchromosomal translocations was higher between more divergent isolates. Interestingly, several isolate-specific rearrangements had common breakpoints, indicating the presence of insertion hot spots (**Note S3**).

**Figure 4.**
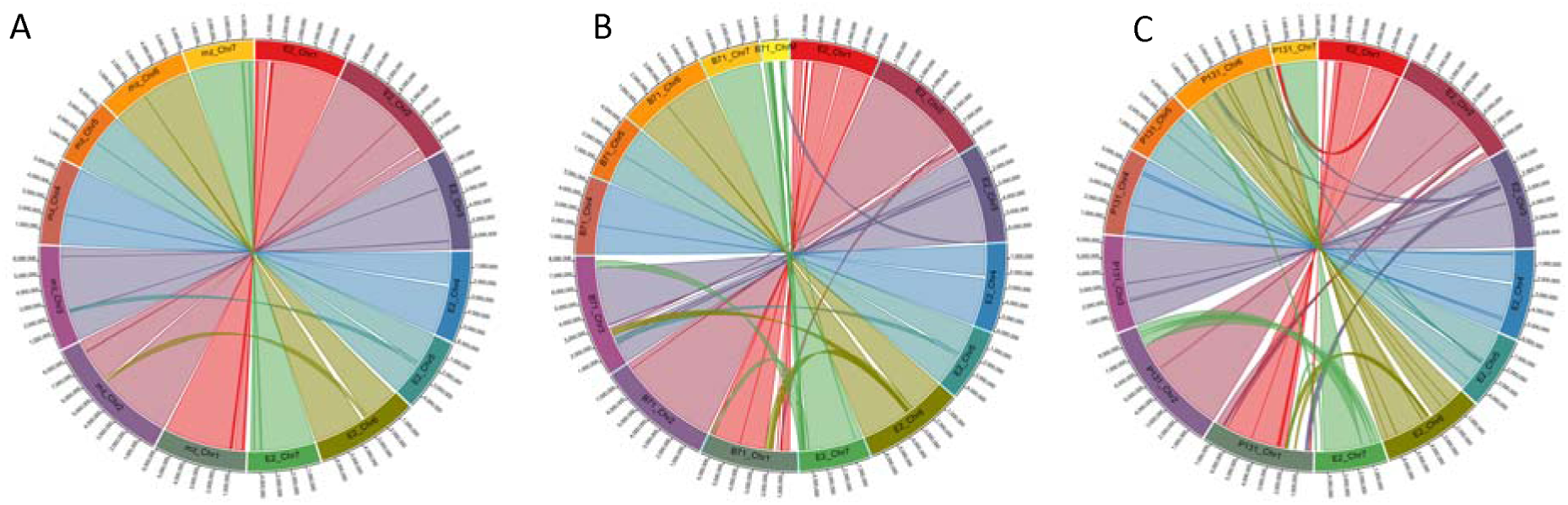
Comparative relationship of finger millet-infecting M. oryzae isolate E2 with MZ5-1-6 (finger millet-infecting) (A), B71 (wheat-infecting) (B) and P131 (rice-infecting) (C)

Reannotation of the repeats in the four strains using a common TE library showed an overall repeat content in the range 10.6% (B71) to 14.2% (P131) (**Table S4**). The P131 genome assembly is gapless (LI *et al*. 2024), likely accounting for the somewhat higher repeat identification. The four strains displayed differential presence and expansion of LTR-retrotransposon families with *Grh*, *Family 13* and *Fosbury* having the highest number of full-length elements in E2/MZ5-1-6, B71 and P131, respectively (**Table S8, Data S3, Note S4)**. The oldest activity of *Family 13* predates the acquisition of host specificity (**Figure 5**), suggesting its presence, likely in low copy numbers, in the common ancestor of rice, wheat and finger millet blast. Recent bursts of amplification following diversification are indicated by identity of the LTR sequences in more than 80% of the elements in both the *MoE* and *MoT* strains. In contrast, *Grh* and *Fosbury* are young elements, consistent with their presence in only the *MoE* and *MoO* strains, respectively (**Figure 5**). Interestingly, *Fosbury* or a closely related element was prevalent in Ethiopian strains in the ET group originating from the West Gojam (15 out of 17 strains), Awi (7/7) and East Wollega (5/5) regions (**Data S1**). Similar to *Family 13* elements, *Fosbury* may have been present in low copy numbers in ancestral blast lineages, and amplified independently in rice and, regionally, in Ethiopian finger millet blast strains after their divergence. Although the oldest insertion events in P131 are estimated to have occurred after the divergence of the rice and ancestral wheat/finger millet blast lineages (**Figure 5**), there is some uncertainty about both the divergence dates and transposon insertion dates due to the relatively low sequence divergence and evolutionary rates used for genes and TEs. Alternatively, the element may have been acquired more recently in Ethiopian finger millet-infecting blast isolates, potentially through horizontal gene transfer (BARRAGAN *et al*. 2024).

**Figure 5.**
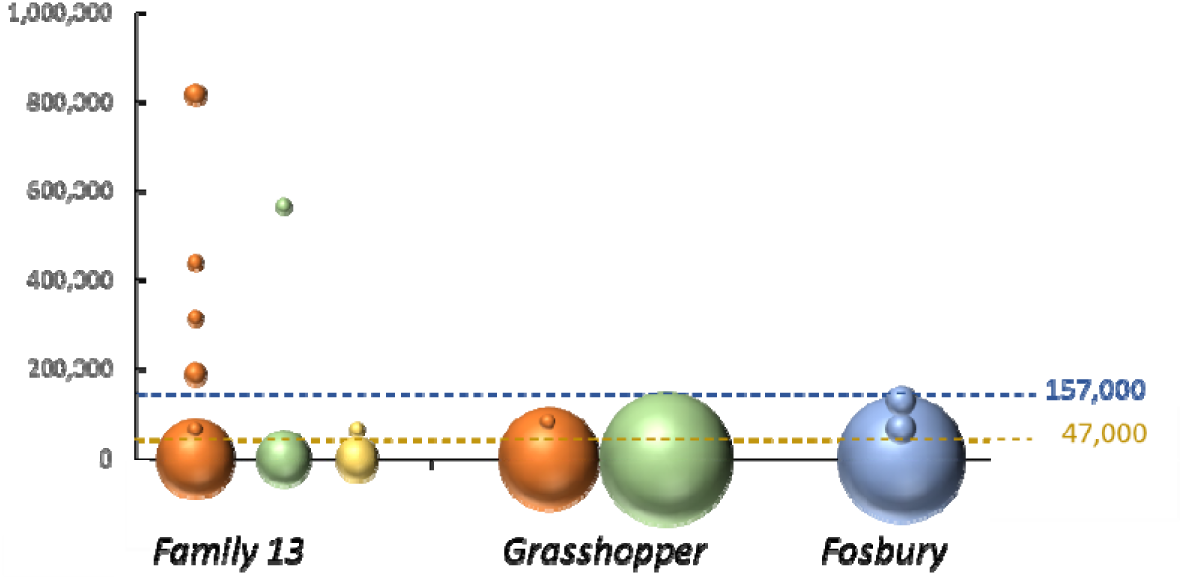
Prevalence and age distribution of LTR-retrotransposons ‘Family 13’, ‘Grasshopper’ and ‘Fosbury’ in E2 (orange), MZ5-1-6 (green), B71 (yellow) and P131 (blue). The Y-axis represents the estimated insertion time in years ago. The center of the dot corresponds to the insertion age, and the size of the dot is proportional to the number of elements with that insertion age. The blue and gold dotted lines indicate the estimated divergence age of E2 (MoE) and P131 (MoO), and E2 and B71 (MoT), respectively (**Table S10**).

### Homologous recombination between Pot2 transposable elements is a key mechanism leading to chromosomal deletions in Ethiopian MoE isolates

The presence/absence pattern observed for the 198,454 genome-wide SNPs across the 226 blast strains revealed the existence of both unique and shared deletions relative to the E2 reference strain with sizes ranging from ∼10 bases to several hundred kilobases (**Figure S5**). An initial manual analysis of 14 deletions for which the breakpoints could be defined at both ends (**Note S5**) showed that two were straightforward deletions, with one being flanked by 4 bp of microhomology (**Data S4**). This suggests that illegitimate recombination as part of the double strand break repair machinery (DEVOS *et al*. 2002) is one mechanism through which deletions occur in *M. oryzae*, albeit not the dominant one. Half of the 14 deletions analyzed were associated with the presence of a *Pot2* DNA transposon (KACHROO *et al*. 1994) and an additional four were associated with a LTR-retrotransposon (*Fosbury*: 2; *Pyret*: 1; *Grasshopper*: 1) (**Data S4**). Expanding this analysis to the genome-wide level (**Figure S6)** in 33 Illumina-sequenced Ethiopian strains with at least 85% membership to the ET population, and excluding strains clonal to the reference strain E2, identified 124 unique deletions >100 bp with breakpoints that could be delineated to a region <3 kb (**Data S5)**. A total of 32.3% of the deletions were associated with a *Pot2* DNA transposon (n=40) and an additional 12.9% with one of the six

LTR-retrotransposons analyzed (*Fosbury*: 6; *Grasshopper*: 4; *Family 13*: 2; *Pyret*: 2; *Family 1*: 1; *Inago2*: 1). Collectively, the data suggest that homologous recombination between TE family members is a major mechanism responsible for removal of DNA, including genes, in *M. oryzae*. A total of 56.5% (70/124) deletions carried at least one gene. The number of deleted genes averaged 24 per strain (range 11 – 47), 17.6% of which were predicted effector genes (range 0% - 34.8%).

### Effector gene repertoire and virulence in the KU and ET genetic groups

A total of 887 genes (874 in the E2 genome and 13 across the 225 short-read assemblies), representing 6.4% of annotated genes in the finger millet pan-genome, were predicted to encode effector proteins (**Data S6**). Of these, 50.4% (447/887) were homologous to effector genes predicted from blast strains infecting other grasses (mostly rice) while the remaining 49.6% (440/887) were newly predicted from our set of 226 *MoE* strains. Loss of effector genes provides a mechanism for pathogens to gain virulence on hosts carrying the cognate *R* genes (INOUE *et al*. 2017; HU *et al*. 2022). Effector gene loss, with loss defined as unambiguous absence (no reads identified corresponding to the gene in the raw sequence data) in at least one of the 226 strains, was seen for 11.6% of the predicted effectors (96/829 effectors for which presence/absence could be established unambiguously in ≥ 80% of strains; see M&M for filtering) (**Data S6**) compared to 8.6% for non-effector genes (**Note S6; Data S7**). The inclusion in this analysis of genes annotated in the 225 short-read assemblies should minimize the absence bias that could arise when only using the annotation from the E2 reference strain, which is an ET strain, in comparing loss between the ET and KU groups. Nevertheless, blast strains from Kenya and Uganda belonging to the KU population had fewer effector genes than strains from Ethiopia (p < 0.001) and Tanzania (p < 0.05) belonging to the ET population (**Figure 6; Table S11**) with 25.0% of effector genes (24/96) having a more than 25 percentage-point higher absence across strains in the KU group than in the ET group. A strong bias for absence in the ET group was observed for only 6.3% (6/96) of effector genes (**Data S6**). Further, 85.4% (82/96) of effector genes were present in at least 75% of the ET strains but only 6.3% (6/96) were absent. The corresponding percentages for the KU strains were 19.8% (19/96) for presence and 29.2% (28/96) for absence. While significant differences were also seen for the presence/absence profile of non-effector genes (**Table S11**), these differences were far less pronounced with 47.0% (484/1030) and 42.2% (435/1030) of genes being present and absent, respectively, in at least of 75% of the ET strains, and 55.9% (576/1030) and 48.3% (497/103) being present and absent, respectively, in at least 75% of the KU strains (**Note S6**).

**Figure 6.**
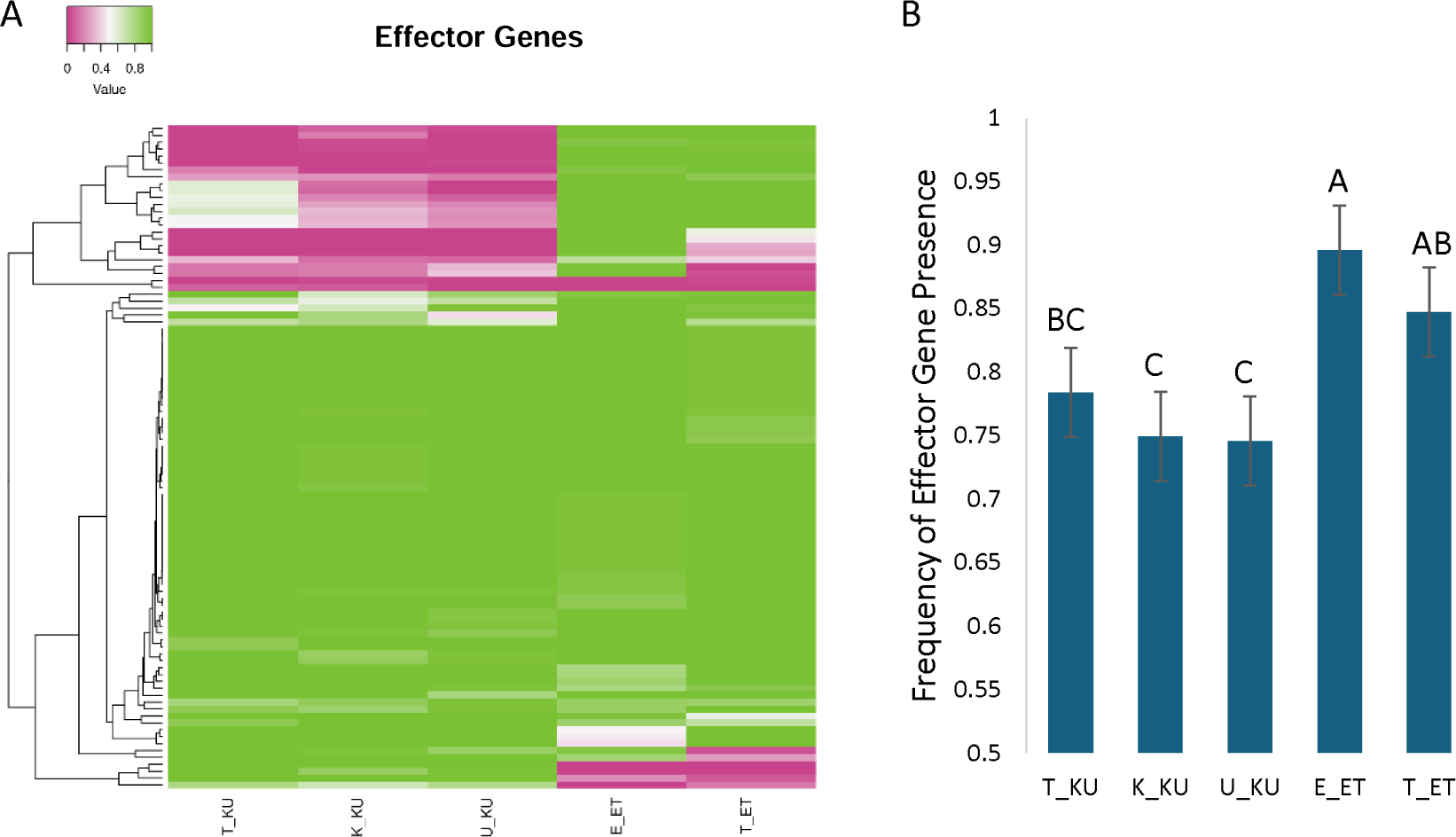
Relative presence of predicted effector genes shown in (A) a heatmap (generated with Heatmapper (BABICKI et al. 2016)) with the most intense green representing genes that are present in all strains within a country/population and the most intense purple representing genes that are absent in all strains within a country/population and (B) a bar graph representing the average frequency of effector gene presence by country/population. Significance was determined by Tukey corrected multiple comparison tests and identified as different letters. Genes that are uniformly present were not included. T_KU, K_KU, and U_KU are Tanzanian, Kenyan and Ugandan blast isolates that belong to the KU population; E_ET and T_ET are Ethiopian and Tanzanian blast isolates that belong to the ET population. For clonal isolates, only a single representative isolate was included in the analyses.

Despite having fewer effector genes, KU strains were significantly more infectious on the largely susceptible Ethiopian accession AAUFM-44 than ET strains (**Figure 7A**). Interestingly, infection levels were not significantly different between KU and ET strains on TZA1637, a relatively blast resistant variety from Tanzania (**Figure 7B**). This is likely because, within the ET population, Tanzanian *M. oryzae* isolates were significantly more virulent on TZA1637 than Ethiopian isolates (**Figure 7B**). While there was no statistically significant difference in the number of effector genes present between Ethiopian and Tanzanian ET strains (**Table S11**), the portfolio of effectors did vary (**Figure 6; Data S6**). Considering that TZA1637 is a Tanzanian accession, the higher virulence of Tanzanian *MoE* strains may reflect adaptation through loss of avirulence genes or gain of virulence genes.

**Figure 7.**
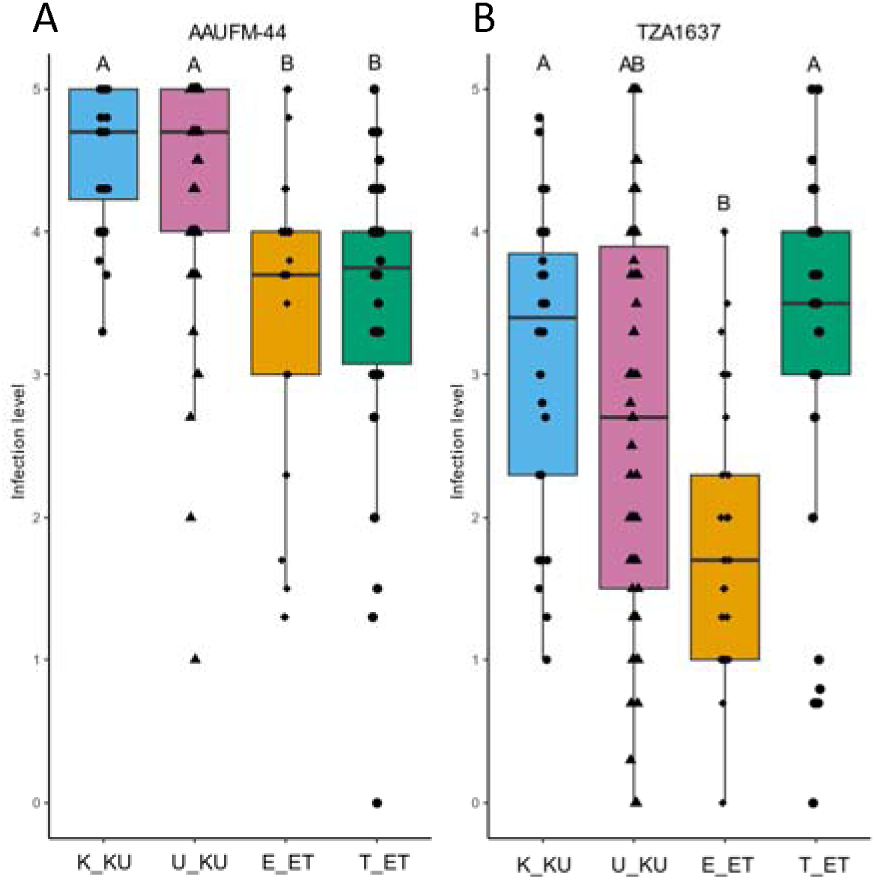
Infection levels (0-5 score; see **Figure S7** for scoring scale) of 207 blast isolates collected during the period 2015-2017 by population group and country of origin on finger millet accessions AAUFM-44 (A) and TZA1637 (B). K_KU and U_KU are Kenyan and Ugandan blast isolates that belong to the KU population; E_ET and T_ET are Ethiopian and Tanzanian blast isolates that belong to the ET population. The underlying data is provided in **Data S8**.

### M. oryzae *genes differentially regulated in compatible and incompatible interactions*

Transcriptome analyses were conducted using an enhanced green fluorescent protein (EGFP)-labeled *M. oryzae* transformant, CKF4046, which retains the virulence of its paternal E2 strain, on leaf sheaths of finger millet accessions TZA1637 and AAUFM-44 (**Note S7**). In contrast to rice, resistance to *M. oryzae* in finger millet is quantitative, and complete resistance is not commonly observed (KATO *et al*. 2000; TAKAN *et al*. 2012). Because TZA1637 is considerably more resistant to the *M. oryzae* CKF4046 strain than the Ethiopian accession AAUFM-44 (**Figure S8**) in both controlled environment and field conditions, we refer to the CKF4046 – TZA1637 interaction as incompatible and the CKF4046 – AAUFM-44 interaction as compatible. The notable difference in infections between compatible and incompatible interactions was consistent at cellular resolution when the growth of EGFP-labeled invasive hyphae was individually observed, or the infection area was quantified based on EGFP fluorescence reflecting hyphal colonization in sheath cells (**Figure 8 and Note S7**). That is, for compatible interactions at 30 hours post-inoculation (hpi), the majority of infections remained in the first infected plant cells, accounting for 3.6% of infection area. By 48 hpi, the invasive hyphae had spread to adjacent cells, increasing the infection area to 22.4%. Incompatible interactions, however, showed a stark restriction of fungal colonization. Even at 48 hpi, all invasive hyphae were retained in the first-invaded cells with the infection area reaching only 0.7%, up from 0.1% at 30 hpi. This increased over time, but contrasting fungal colonization between compatible and incompatible interactions was consistently reflected in the fungal transcript abundance in RNA-seq data of the infected leaf sheaths.

**Figure 8.**
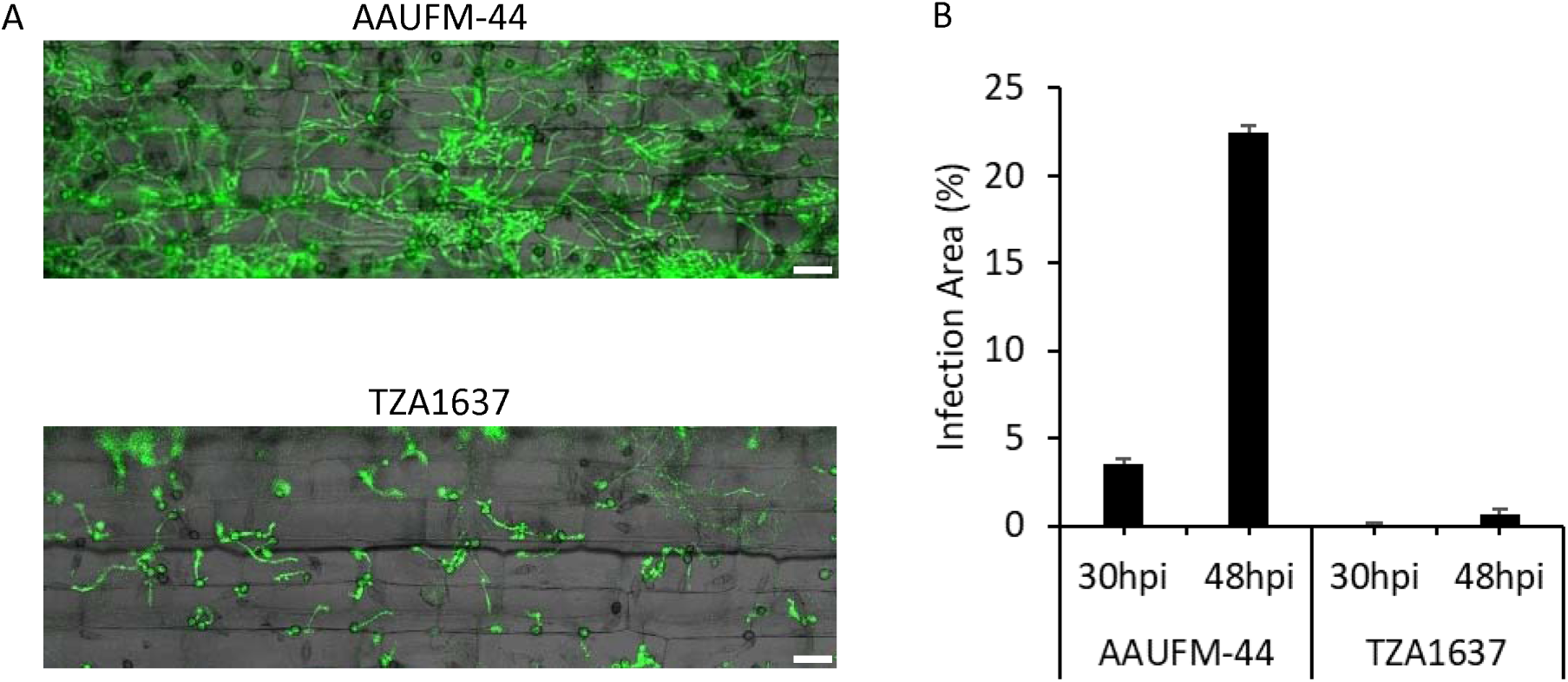
M. oryzae infection on compatible and incompatible finger millet cultivars. (A) Representative confocal microscopy images of finger millet AAUFM-44 (susceptible) and TZA1637 (resistant) sheath cells infected by EGFP-tagged M. oryzae isolate E2 (CKF4046) at 48 hpi. Bars = 50 μm. (B) Quantification of infection area (%) in AAUFM-44 and TZA1637 at 30 and 48 hpi, based on ImageJ analysis of two confocal images for each time point (30 and 48 hpi) and interaction (AAUFM-44 and TZA1637).

RNA-seq of the infected leaf sheaths yielded 1.7% and 13.3% fungal transcripts (transcripts aligning to the E2 genome assembly) at 30 hpi and 48 hpi, respectively, in the incompatible interaction, and 12.0% and 71.0% at 30 hpi and 48 hpi, respectively, in the compatible interaction (**Data S9**). A total of 7621 fungal genes were expressed (average normalized read count across three replicates ≥ 5) during infection (**Table S12**) with 2048 and 1257 genes being differentially expressed (Padj < 0.01; ≥ 2-fold transcript level difference) between the incompatible and compatible interactions at 30 hpi and 48 hpi, respectively (**Data S10, S11)**. Gene onthology (GO)-enrichment analysis identified ‘structural constituent of ribosome’ as the main molecular function (MF) term for genes upregulated during the compatible interaction at 30 hpi (4.7e^-61^) as well as for genes upregulated during the incompatible interaction at 48 hpi (4.7e^-6^), reflecting protein synthesis to support fungal growth (**Figure S9A, D)**. In agreement, 72% (36/50) of the top-50 (lowest Padj-value) functionally annotated genes with ≥ 2-fold higher transcript levels in the compatible *versus* incompatible interaction at 30 hpi (**Data S12**) were ribosomal proteins. However, no genes encoding ribosomal proteins were identified among the top-50 genes upregulated in the incompatible interaction at 48 hpi. The majority of the top-50 genes were annotated as ‘putative uncharacterized protein’, likely because less functional validation has been done of fungal proteins expressed at later infection time points in incompatible interactions. The term ‘oxidoreductase activity’ was the top MF term enriched in both the incompatible interaction at 30 hpi (1.4e^-5^) and the compatible interaction at 48 hpi (1.9e^-6^) (**Figure S9B, C)**. This term broadly covers enzymes contributing to pathogen virulence and survival. Of the top-50 genes with upregulated expression in the incompatible interaction at 30 hpi, several had previously been shown to encode proteins with a role in *M. oryzae* pathogenicity to rice, including *BAS3* (Biotrophy-associated secreted protein 3), *SPM1* (a subtilisin-like serine protease required for autophagy and pathogenicity), *MAS3* (*Magnaporthe* appressoria-specific protein 3; involved in suppressing rice’s innate immunity (GONG *et al*. 2022)) and *ALO1* (ALO1-D-arabinono-1,4-lactone oxidase; required for fungal growth, conidiogenesis and pathogenicity (WU *et al*. 2022). Salicylate hydroxylase and 2,3-dihydroxybenzoic acid decarboxylase, two enzymes that degrade salicylic acid (WRIGHT 1993; MARTINS *et al*. 2015; ROCHELEAU *et al*. 2019), were also upregulated in the incompatible response. Strikingly, while only 4% (2/50) and 0% of genes (0/50) with relative higher transcript levels in the compatible interaction at 30 hpi and 48 hpi, respectively, were predicted to be effector genes, the corresponding values were 22% (11/50) and 60% (30/50) for the genes upregulated in the incompatible interaction.

Co-expression analysis across genotypes and time points as well as a 4-day-old axenic culture grouped 10,355 gene models corresponding to annotated genes in 13 clusters (**Figure S10; Data S13**). The largest cluster (cluster 5; n=5992) consisted of genes with similar expression levels across the five conditions. Only 2.2% (130/5992) of the annotated genes in this cluster (compared to 6.8% across all clusters) are predicted to encode effectors (**Table S13; Data S13**). This cluster includes the known *Avr* gene *Avr-Pi54*. In contrast, 62.1% (151/243) of the annotated genes in cluster 3 were predicted to be effector genes. The overall expression pattern in cluster 3 is downregulation between 30 hpi and 48 hpi in a compatible interaction and upregulation in an incompatible reaction (**Figure S10**). Cluster 3 includes *Avr-Pi5, Avr-Pii, Avr-Pik* and *Avr1-CO39* (**Data S13**). Similarly, 26.8% (26/97) and 28.9% (115/398) of annotated genes in clusters 11 and 13, respectively, were predicted or validated effector genes (**Table S13**). As in cluster 3, fungal genes were more highly expressed during an incompatible interaction, but expression decreased between 30 and 48 hpi (**Figure S10**). This pattern was seen for the known *Avr* genes *Avr-Rmg8, Avr-Pi9* and *Avr-Pizt*.

### Two novel biosynthetic gene clusters are highly upregulated in the incompatible reaction

A total of 67 biosynthetic gene clusters (BGCs) were identified in the E2 genome (**Table S14, Data S14**), two of which are of particular interest. Cluster 5.01 (EcMO5g00088700 - EcMO5g00088800) on chromosome 5 encompasses 11 genes, eight of which were functionally annotated and highly upregulated (log2fold change in range 4.2 to 9.0) in the incompatible compared to the compatible interaction at 48 hpi. Six were also highly upregulated at 30 hpi (**Data S14; Note S8**). The second cluster of interest, 5.12, comprises six genes (**Note S9)** that are highly upregulated (log2fold change in range 7.7-11.0) at 30 hpi in the incompatible compared to the compatible interaction. The metabolite(s) generated by clusters 5.01 and 5.12 are unknown, but the presence of a polyketide synthase (5.01) or nonribosomal peptide synthetase (5.12) as well as an ent-kaurene synthase, gibberellin cluster GA14 synthase and geranylgeranyl pyrophosphate synthase 2, enzymes potentially involved in the gibberellic acid biosynthesis pathway (TUDZYNSKI AND HÖLTER 1998), in both clusters, suggest they could represent novel terpene-polyketide and terpene-amino acid hybrids (YEE *et al*. 2020; YAN AND MATSUDA 2024). The high upregulation in incompatible interactions and membership to co-expression cluster 11 strongly suggest a role in virulence. Intriguingly, differential loss of individual genes or, in some case, the entire gene cluster, can be observed across *M. oryzae* strains with different host specificities with loss not necessarily following phylogenic relationships between the strains (**Table S15)**.

## Discussion

### M. oryzae *isolates infectious on finger millet in eastern Africa diverged into two genetic groups*

Finger millet blast disease, caused by *Magnaporthe oryzae*, is the main biotic constraint to finger millet production in eastern Africa. Infection of the peduncle (neck blast) is the most detrimental to grain yield followed by infection of panicle (head blast). *MoE* strains from eastern Africa form two genetic groups that differ in their overall genetic make-up, composition of the repeat fraction of the genome, portfolio of effector genes and virulence level. The two eastern African groups are sister to *MoE* strains from Asia, demonstrating that there are at least three major *MoE* genetic groups. No phylogenetic separation was seen between isolates collected from Eastern Uganda and Western Kenya after a 10-year hiatus, indicating that KU isolates had undergone little genetic divergence during that time span.

Admixture (<90% membership to a single population) was predominantly (17 out of 18 strains) observed in the East Wollega (n=9) and West Wollega (n=8) regions in western Ethiopia (**Data S1**). Eight out of the nine strains (three of which were clonal) isolated from West Wollega were admixed with about equal membership to the ET and KU groups, potentially indicating higher fitness or virulence of the admixed strains in this region. Co-occurrence of ET strains (n=5) and KU strains (n=3) in East Wollega provided opportunity for intercrossing, as demonstrated by the presence of admixed strains (n=9; five clonal). However, in other areas where ET and KU strains co-occurred such as the West Gojam region in NW Ethiopia, Busia district in Kenya and Njombe district in Tanzania, admixed strains were not observed. Sexual reproduction in the blast fungus only occurs when two strains with opposing mating type meet (KANG *et al*. 1994). Mating type-specific PCR assays of 224 eastern African *MoE* isolates, including 127 from the current study (**Data S1)**, revealed near-equal levels of *MAT1-1* and *MAT1-2* mating types overall, although the pattern of co-occurrence varied among the countries and districts surveyed (SHITTU 2018). Further, mating cross assays showed that only 61% of the 224 isolates collected during 2015-2017 were fertile (K: 51%; E: 75%; T: 50%; U: 66%), compared to 89% of Kenyan (n=97) and 85% of Ugandan isolates (n=175) obtained from infected *E. coracana* tissues collected in Kenya and Uganda during 2000-2004 (TAKAN *et al*. 2012). The overall pattern of the mating type distribution and the fertility status of the *M. orzyae* isolates associated with finger millet in eastern Africa suggest predominance of asexual reproduction with limited potential for episodic sexual reproduction and recombination contributing to the stability of the pathogen genetic groups.

The presence of genetically related *MoE* strains in Kenya and Uganda is anticipated considering the geographic proximity of finger millet cultivation areas in Eastern Uganda and Western Kenya (**Figure 1**), and the free movement of finger millet seeds between the two countries. The close relationship between Tanzanian and Ethiopian isolates is more difficult to explain. The cultivars grown in Ethiopia and Tanzania are distinct (DEVOS *et al*. 2023), excluding host adaptation as the likely cause. Köppen climate zones, however, are temperate for the finger millet growing areas of Tanzania and Ethiopia (class C), while Kenya and Uganda are largely tropical (class A) (**Data S1**). Pathogen diversity may thus be driven by adaptation to climate variables (CABALLOL *et al*. 2024). Differential expansion of transposable element families, some of which were likely present at low copy numbers in the common ancestor of rice and wheat blast, combined with varying levels of asexual reproduction, contributed to the formation of distinct blast populations.

### Origin and spread of the finger millet blast fungus

Blast disease was first recorded on finger millet in Uganda in 1933 (EMECHEBE 1975) but was known as a major disease of rice in Uganda as early as 1920 (SMALL 1922). Rice, introduced into Uganda during British colonial rule by Indian traders, reached peak cultivation in the mid-1920s before declining, most likely due to blast disease (KIKUCHI *et al*. 2013). Despite the fact that the first recording of finger millet blast in Uganda followed a major outbreak of rice blast disease, the time frame of the divergence of rice- and finger millet-infecting blast strains some 157,000 years ago (**Table S10**) clearly indicates that finger millet blast in eastern Africa did not originate through a host jump from rice. This is supported by the closer phylogenetic relationship of *MoE* isolates to wheat blast (**Figure 3**; GLADIEUX *et al*. 2018a) with the estimated *MoE* – *MoT* divergence dating back to around 47,000 years ago (**Table S10**). Adaptation of *M. oryzae* to finger millet therefore predates the crop’s domestication which occurred around 5,000 years ago (DEVOS *et al*. 2023). In Africa, finger millet blast may have been present on *Eleusine indica*, the A-genome donor to tetraploid finger millet, or on *Eleusine coracana* subsp. *africana*, the wild progenitor to finger millet, long before the disease was recorded on the crop. Alternatively, finger millet blast may have been introduced into eastern Africa in more recent times, potentially through contamination of grain lots with blast-infected weed seeds. The potential for contaminated seed to spread blast disease has been demonstrated in both rice and wheat (FAIVRE-RAMPANT *et al*. 2013; MARTINEZ *et al*. 2021). The close relatedness between *E. indica-* and finger millet-infecting blast isolates in the KU group with *E. indica* isolates from Asian and South American countries where finger millet is not cultivated as a crop (**Figure 3**), supports spread through blast-contaminated seed rather than host jump but also suggests lowland eastern Africa as the area of adaptation to the *Eleusine* host.

### Homologous recombination between transposable elements is a key mechanism for DNA loss

Loss of effector genes is an effective way to gain virulence on plant hosts carrying the cognate *R* genes. Effector genes are often located within repetitive DNA (**Figure S11**; KIM *et al*. 2019b), and our data demonstrate that homologous recombination between members of the same TE family is a main mechanism in *M. oryzae* for removing DNA. A total of 194 non-redundant genes were removed in 33 Ethiopian *MoE* strains as part of 124 deletions, ∼45% of which were caused by homologous recombination between TEs (**Data S5**). Despite effector genes representing only ∼6.4% of the total gene number in the genome, 18.6% (36/194) of the genes located in the 124 analyzed deletions were predicted effector genes. The frequency with which homologous recombination between TE family members occurs is almost certainly higher than what was observed in our study. Our genome-wide analysis only identified deletion-associated TEs if at least one of the two TEs involved in the homologous recombination was absent from the reference strain (**Figure S6**). Active (young) elements are more likely to be differentially present in closely related strains and will therefore preferentially be identified as associated with a deletion. Other factors that can drive the frequency of homologous recombination are TE copy number and their distribution across the genome. While all TEs analyzed clustered in the same areas of the genome (**Figure S11**), *Pot2* DNA transposons had the highest copy number (**Table S9**), explaining their considerably higher association with deletions (32.3%) than the six LTR-retrotransposons combined (12.9%). We postulate that homologous recombination between TEs is a key mechanism for DNA removal in *Magnaporthe oryzae*, but that the elements driving the DNA loss will vary by lineage and genetic groups within a lineage concomitant with the TE composition.

### MoE *isolates in the KU group have fewer effectors and higher levels of virulence*

The observation that KU strains carry fewer effector genes (**Figure 6; Data S6**) but exhibit higher virulence on AAUFM-44 (**Figure 7A**) is intriguing. Given that some effectors can function as avirulence factors, triggering host immunity (VALENT AND KHANG 2010), it is possible that KU strains lack such genes that are likely present in ET strains. Since higher virulence can also result from virulence effectors targeting host susceptibility factors (GORSHKOV AND TSERS 2022), alternatively, KU strains, despite having a smaller effector repertoire, may carry such virulence factors that are absent in ET strains. In either case, our infection results suggest the presence of host immunity genes in AAUFM-44 recognizing avirulence factors likely absent in KU strains but present in ET strains, or the presence of host susceptibility factors targeted by virulence factors in KU but not ET strains.

### *Transcript profiling suggests avirulence activity during incompatible* MoE-finger millet *interactions*

Identifying effector genes with avirulence activity in *MoE* is instrumental in discovering their cognate host *R* genes in finger millet, and for the strategic deployment of these *R* genes in finger millet breeding programs. Although several avirulence genes have been identified in *MoE*, they confer avirulence on other hosts such as wheat and weeping lovegrass (ASUKE *et al*. 2021; ASUKE *et al*. 2023; MASAKI *et al*. 2023). No avirulence genes effective in finger millet have been identified.

Our comparative transcriptomics analyses combined with live cell imaging of infections suggest that avirulence genes are specifically induced during the incompatible interaction, which paves the way for identifying these genes. Some 30% of predicted effector genes were upregulated in the incompatible interactions with TZA1637 compared to the compatible interaction with AAUFM-44. More than 65% of the effector genes in the three co-expression clusters most highly enriched for effector genes (clusters 3, 11 and 13) had significantly higher transcript levels at 48 hpi in the incompatible compared to the compatible interaction (**Table S13**). More than half of the effector genes in clusters 11 and 13, which showed overall downregulation between 30 hpi and 48 hpi, also had significantly higher transcript levels in the incompatible compared to the compatible interaction at 30 hpi. These clusters include seven of the eight homologs of known avirulence genes from rice-infecting *M. oryzae*, supporting the hypothesis that these induced effectors may function as avirulence factors in *MoE* – finger millet interactions.

Importantly, live-cell fluorescent microscopy at 48 hpi revealed a clear restriction of invasive hyphal growth in the resistant TZA1637, in contrast to continued invasive growth in the susceptible AAUFM-44 (**Figure 8 and Note S7**). The presence of EGFP fluorescence in these restricted hyphae inside TZA1637 cells indicates that the hyphae remained viable. We predict that the presumed resistance response in TZA1637 is likely distinct from a typical hypersensitive response, which is often associated with the death of both host cells and invasive hyphae (KANKANALA *et al*. 2007). The viable short invasive hyphae in the resistant TZA1637 cells at 48 hpi may explain the seemingly counterintuitive finding of higher expression levels of effector genes, including avirulence genes, in the incompatible interaction. As well documented in *MoO*, where effector genes are highly expressed from short invasive hyphae during early biotrophic invasion of rice cells, *MoE* may similarly continue to express effector genes from short, restricted hyphae in TZA1637. It raises the possibility that effectors induced during incompatible interactions may follow similar delivery routes – via Biotrophic Interfacial Complex (BIC) and translocation into host cells – as seen in compatible interactions (**Note S7**; (KHANG *et al*. 2010; GIRALDO *et al*. 2013; OLIVEIRA-GARCIA *et al*. 2023), but with distinct functional outcomes. This study also opens an exciting opportunity to functionally validate the effectors identified by generating isogenic strains differing by a single avirulence or virulence gene and assessing their infection potential.

### Non-effectors encoded by genes with increased expression in an incompatible interaction may attenuate host plant defense

Genes that are not predicted to encode effector proteins, and that are upregulated at early time points in the incompatible *vs.* the compatible reaction may, similarly to effectors, play a role in virulence through modulation of plant defense. The function of some of the most significantly upregulated genes at 30 hpi in clusters 3, 11 and 13 that were also significantly upregulated at 48 hpi (**Data S12**) provide examples of fungal proteins that can increase virulence by assisting the pathogen with host cuticle penetration by degrading plant cutin and waxes, and attenuating host defense systems by catabolizing or detoxifying defense molecules such as salicylic acid and phytoalexins (**Note S10**). Some fungal proteins also increase tolerance to host-produced toxins by transporting them out of fungal cells. Upregulation of these genes as early as 30 hpi in the incompatible interaction and the action mode of five of the six encoded proteins described suggest secretion before or during hyphal invasion of host cells.

## Conclusions

Sequence analysis of 226 finger millet-infecting *M. oryzae* strains from eastern Africa demonstrate their organization in two genetic groups that are sister to an Asian *MoE* group. Group members differed in their overall genetic make-up, transposable element composition, effector gene content and virulence level. Diversification into the two genetic groups may have been driven by climate factors. DNA deletions caused by homologous recombination between transposable elements encompassed a higher-than-expected number of effector genes, demonstrating that this is an important mechanism through which strains lose effector genes to enhance virulence. For Ethiopian strains, the element most frequently associated with deletions was the DNA transposon *Pot2,* which was the highest copy number element analyzed in our study. The preferential deletion of effector genes can be attributed to their predominant location in repeat-rich genomic regions. The portfolio of more than 800 predicted effector genes identified in our study provides an important resource for future identification of avirulence genes and their cognate host *R* genes. The observation that a subset of effector genes, including seven out of the eight known avirulence genes that were present in the effector gene set, are significantly upregulated in an incompatible compared to a compatible interaction provides an important criterion for prioritizing genes for functional validation. This paves the way to identify the cognate *R* genes in finger millet germplasm to improve the crop for resistance to blast disease.

## Materials and Methods

### Finger millet-infecting Magnaporthe oryzae strains

A total of 207 single-spore *Magnaporthe oryzae* isolates were obtained from finger millet panicles and peduncles, collected in Ethiopia, Kenya, Uganda and Tanzania during 2015-2017, that displayed symptoms of blast disease. Strain isolation was performed following the standard protocol in the Khang lab. Briefly, infected tissues were cut into 1 cm pieces, surface sterilized in 70% ethanol, and rinsed with sterile distilled water. After repeating the surface sterilization twice, the sterilized pieces were placed on V8 medium containing penicillin and streptomycin antibiotics (Hyclone Cat# SV30010). Conidia produced in 3-5 days were observed under a microscope at 400x magnification to confirm *M. oryzae* based on its well-documented, uniquely shaped conidia (MAKAJU *et al*. 2016). A single conidium was isolated and cultured on oatmeal agar (OMA) with filter discs. The filter discs fully colonized by the fungus were dried in a desiccator for four weeks and stored at −20 °C for long-term preservations. The fungal strains were cultured on OMA plates at 25 °C under continuous light. An additional 19 finger millet-infecting blast isolates had previously been isolated from finger millet leaf, panicle and peduncles collected during 2000 and 2002 in Kenya and Uganda (TAKAN *et al*. 2012). We refer to the latter strains as ‘historical isolates’. Detailed information on the 226 isolates is provided in **Data S1**.

### Finger millet accessions

Accessions IE 2555 (Kenya), 214988 (Zambia), Gulu-E (Kenya), AAUFM-44 (Ethiopia) and TZA1637 (Tanzania) were used in infection assays. AAUFM-44 and TZA1637 were used for RNA-Seq. Information on the accessions is available from DEVOS *et al*. (2023).

### *Whole-genome Illumina sequencing of finger millet-infecting* M. oryzae *strains*

Fungal conidia were harvested from ∼10-day-old cultures on OMA plates. A suspension of 1×10^5^ spores/ml in distilled water was inoculated into liquid complete medium and shaken at 25 °C, 100 rpm for four days in the dark. Fungal mycelia were collected, washed by filtration to remove extra water, frozen immediately in liquid nitrogen and stored at −80 °C until used for DNA extraction. Frozen mycelia (∼0.5 g) were ground into a fine powder in liquid nitrogen, and DNA was extracted for one hour at 50 °C using a modified Cetyltrimethylammonium Bromide (CTAB) protocol with the CTAB buffer containing 100 µg/mL proteinase K, 2% PVP-40, and 0.2% beta-mercaptoethanol. Proteins were removed through two gentle extractions with one volume of chloroform:isoamyl alcohol (24:1 v:v). DNA was precipitated with 2.5 volumes of ethanol after the addition of 1/10th volume of 3M KAc. The DNA pellet was washed with 70% ethanol, air dried for 20 minutes, and dissolved in 100 µL Tris-EDTA (TE) buffer at room temperature. The concentration and purity of the resulting DNA were determined using a Qubit fluorometer (Thermo Fisher Scientific, Carlsbad, CA, USA). Library preparation and short-read sequencing of all strains on an Illumina NovoSeq6000 platform (400 bp insert libraries; 2×150 bp) was done at the HudsonAlpha Institute for Biotechnology in Huntsville, Alabama, using standard protocols.

### *Whole-genome PacBio sequencing of the Ethiopian* M. oryzae *isolate E2*

Approximately 0.5 g mycelium of *M. oryzae* isolate E2 was harvested from fresh cultures and ground to a fine powder in liquid nitrogen. The ground material was resuspended in 10 mL of extraction buffer [2% (w/v) CTAB, 100 mM Tris-HCl (pH 8.0), 20 mM EDTA, 1.4 M NaCl, and 1% (w/v) polyvinylpyrrolidone]. The suspension was incubated at 65°C for 60 min, with intermittent mixing. After incubation, an equal volume of chloroform:isoamyl alcohol (24:1 v:v) was added, followed by vigorous shaking and centrifugation at 4,000 rpm for 10 min. The aqueous phase was transferred to a new tube, and DNA was precipitated using 0.6 volumes of cold isopropanol. After a 15-minute incubation at room temperature, the DNA was pelleted by centrifugation at 1,000 rpm for 10 min, washed with 70% ethanol, air-dried, and finally resuspended in 100 µL of 1/10^th^ TE buffer. Library preparation and long-read sequencing of isolate E2 was done on a PACBIO SEQUEL II platform at the HudsonAlpha Institute for Biotechnology in Huntsville, Alabama.

### *Assembly of the* M. oryzae *strain E2* genome

The E2 version 1.0 assembly was generated by assembling the 1,020,525 PacBio reads (**Table S2**; 235x sequence coverage) using the MECAT v1.4 assembler (XIAO *et al*. 2017) and subsequently polished using ARROW v2.1 (CHIN *et al*. 2013). This produced an initial assembly of 29 scaffolds (29 contigs), with a contig N50 of 6.1 Mb, and a total genome size of 44.3 Mb (**Table S3**).

The *M. oryzae* isolate MZ5-1-6 genome obtained from GÓMEZ LUCIANO *et al*. (2019) was broken into 3,884 nonredundant, non-overlapping 1,000 bp syntenic markers which were used to identify misjoins in the initial MECAT assembly. Misjoins were characterized as a discontinuity in the MZ5-1-6 linkage groups. A total of four breaks were identified and resolved. The resulting broken contigs were then oriented, ordered, and joined together into seven chromosomes using *M. oryzae* MZ5-1-6 synteny. A total of 11 joins were made during this process. Each chromosome join was padded with 10,000 Ns. Significant telomeric sequence was identified using the (TTAGGG)_n_ repeat, and care was taken to make sure that it was properly oriented in the release assembly. The remaining scaffolds were screened against bacterial proteins, organelle sequences, GenBank nr and removed if found to be a contaminant.

Finally, homozygous SNPs and INDELs were corrected in the release consensus sequence using ∼50x coverage by Illumina reads (2×150, 400 bp insert) by aligning the reads using BWA-mem 0.7.17-r1188 (LI 2013) and identifying homozygous SNPs and INDELs with the UnifiedGenotyper tool in GATK v3.6-0 (MCKENNA *et al*. 2010). A total of three homozygous SNPs and 941 homozygous INDELs were corrected in the release. The final version 1.0 release contains 44.35 Mb of sequence, consisting of 33 contigs with a contig N50 of 5.8 Mb and a total of 99.1% of assembled bases in chromosomes.

Completeness of the euchromatic portion of the version 1.0 assembly was assessed using protein sequences from *M. oryzae* (strain 70-15 / ATCC MYA-4617) obtained from UniProt (https://www.uniprot.org/proteomes/UP000009058). The aim of this analysis was to obtain a measure of completeness of the assembly, rather than a comprehensive examination of gene space. The transcripts were aligned to the assembly using BLAT v35.0 (KENT 2002) and alignments with ≥ 90% identity and ≥ 85% coverage were retained. The screened alignments indicate that 93.53% of the *M. oryzae* strain 70-15 proteins aligned to the E2 strain version 1.0 release.

### Semi-de novo *assembly of the Illumina-sequenced* M. orzyae *strains*

Low quality reads and adaptors were removed from the raw Illumina reads using fastx_trimmer (FASTX-Toolkit v0.0.14) (HANNON 2010) using the parameter *-q 30*. The preprocessed reads of isolates sequenced to a depth >15x were assembled using SOAPdenovo2 version r240 (LUO *et al*. 2012) with a k-mer length of 41 bp (parameters *avg_ins 500, asm_flag 3, rank 1, pair_num_cutoff 3, map_len 32, -D 1, -K 41*). The resulting contigs were aligned to the whole-genome assembly of E2 using MUMmer v4.0.0 (MARÇAIS *et al*. 2018) with default parameters. The position and orientation of the contigs were extracted from the MUMmer output, and this information was used to order and orient the contigs using in-house scripts.

### Repeat annotation and characterization

A combined repeat library for E2, MZ5-1-6, B71 and P131was constructed through *de novo* identification and classification of repetitive elements using the pipeline ‘Extensive *de-novo* TE Annotator’ (EDTA v2.2.0; (OU *et al*. 2019). Default parameters were used for all softwares unless stated otherwise. Genomic repeat annotation was performed via homologous sequence searches against the repeat library using RepeatMasker (SMIT *et al*. 2013-2015).

Full-length LTR elements within the four genomes were identified using LTR_FINDER v1.07 (XU AND WANG 2007). Solo-LTRs were detected by homologous genomic searches against the full-length LTR elements using LTR_retriever v2.9.9 (OU AND JIANG 2018) . Solo_LTRs with target-site duplications (TSDs) were retained for further analyses. The insertion dates of LTR elements were estimated based on an evolutionary rate of 3.1e^-8^ substitutions/site/year (1.5x the average rate of 2.16e^-8^ (LATORRE *et al*. 2020) and 1.98e^-8^ (GLADIEUX *et al*. 2018b)) to account for the faster evolutionary rate of TEs). LTR families were defined by sequence similarity of the 50 bp LTR terminal regions. If the similarity between two LTR ends exceeded 90%, the elements were classified within the same family. Copy number and dating analyses were conducted for families that contained at least two full-length LTR elements or at least one full-length and one solo-LTR elements in at least one of the four genomes.

To estimate TE copy numbers in the Illumina-sequenced strains, the *semi-de novo* assemblies of the 226 blast isolates were used as queries in BLASTN searches against the LTRs from the retrotransposons *Grasshopper, Fosbury, Family1, Family13, Inago2* and *Pyret* and the full-length DNA transposon *Pot2* using default parameters. The number of BLAST hits ≥ 25 bp and with ≥ 95% similarity were recorded for each strain. The coverage of each base position in a TE by BLASTN alignments was extracted using a custom python script.

### Gene annotation of the E2 and pan-genomes

Putative protein-coding sequences (CDS) in the *M. oryzae* E2 genome were predicted from the repeat-masked E2 genome using MAKER v.3.01.03 (HOLT AND YANDELL 2011) with default parameters, incorporating reference proteins from the rice-infecting *M. oryzae* strain 70-15 (DEAN *et al*. 2005) and the finger-millet infecting strain MZ5-1-6 (GOMEZ LUCIANO *et al*. 2019), and 33,498 *M. oryzae* protein sequences hosted by NCBI (Retrieved Oct 13^th^, 2020). The predicted gene structures were then manually curated using RNA-seq data generated as part of this manuscript (PRJNA1279063) using Web Apollo v2.0.8 (LEE *et al*. 2013). Putative functions of the annotated proteins were assigned using BLAST2GO based on homology to BLASTP hits (pvalue 1e-5, other parameters default value) to NCBI’s nonredundant fungal protein database.

To identify isolate-specific genes in the *semi-de novo* genome assemblies of the 225 Illumina-only sequenced *M. orzyae* strains, we first used RepeatMasker 4.0.9 (SMIT *et al*. 2013-2015) to mask the repetitive elements and genes identified in the E2 genome assembly. We then used the MAKER pipeline to annotate the unmasked genome sequence using the same datasets and parameters as for the E2 annotation. The resulting gene models were filtered to obtain a non-redundant gene set. The criteria used were ≥70% identity at the protein level over ≥70% of the length of the protein. We also removed genes that, based on raw read alignments, were present in E2 or were absent from the strain in which they were annotated as likely artefacts. Further, because short genes were greatly overrepresented in the short-read annotations, genes with a cDNA length < 200 bp were removed. For consistency, we also removed genes with cDNA length < 200 bp from the E2 annotations for all data analyses.

### De novo *prediction of effector genes*

We used EffectorP v2.0 (SPERSCHNEIDER *et al*. 2018) to predict effector candidates from the set of annotated E2 and pan-genome genes, SignalP v5.0 (ALMAGRO ARMENTEROS *et al*. 2019) to predict proteins with a signal peptide, and TMHMM v2.0 (KROGH *et al*. 2001) to predict proteins without a transmembrane domain. Additionally, we used Phobius v1.01 (KÄLL *et al*. 2004) to predict signal peptides and transmembrane domains separately, and GPIPredictor (CAO *et al*. 2009) to predict proteins with a glycosylphosphatidylinositol (GPI) anchor. Because effector proteins are typically short, we excluded proteins with a sequence length longer than 300 amino acids. Finally, we used dbCAN v3.0.3 (ZHENG *et al*. 2023) to identify CAZymes and BLASTP against the MEROPS protease database (RAWLINGS *et al*. 2013) to identify proteases. The resulting set of EffectorP-identified proteins with a sequence length smaller than 300 amino acids, a signal peptide, no transmembrane domains or GPI anchor, and without either a CAZyme or protease domain, were considered potential effector candidates.

Additionally, a list of 435 validated and predicted effector genes (**Data S15**) from other grass-infecting *Mo* strains was compiled, and used as queries in a BLASTN search against the E2 annotated genes with an e-value threshold of 1e^-5^. The resulting hits were used to label effector genes identified through the *de novo* pipeline as well as discover effectors that were missed by the pipeline.

### Determining gene loss from Illumina sequence data

A reference gene set consisting of all cDNA sequences ≥ 200 bp annotated in E2 and the short-read assemblies plus 1 kb of upstream and downstream sequence was generated. The cleaned Illumina reads for each strain were aligned to this reference using Bowtie2 v.2.2.9 (LANGMEAD AND SALZBERG 2012). For each gene in the reference gene set, the coverage length and depth in each blast strain were extracted from the Bowtie alignment file (SAM file) using in-house scripts. A gene was considered present in a blast isolate if ≥ 90% of the mRNA length was covered by Illumina reads at an average depth ≥ 2, excluding the flanking sequence. Genes not covered by any reads were considered absent. Coverage over < 90% of the mRNA length was recorded (**Data S6, S7**), but was considered ambiguous with regards to presence/absence. Genes with ambiguous scores in more than 20% of the strains were excluded when determining the number/percentage genes that displayed presence/absence across the Illumina-sequenced strains. Admixed strains (<90% membership to a single population) were also excluded and clonal isolates were represented by a single isolate.

### SNP calling

Read processing, and single nucleotide polymorphism (SNP) and insertion/deletion (INDEL) calling and filtering were performed following the pipeline described by QI *et al*. (2018). In brief, the cleaned Illumina reads were aligned to the E2 reference genome using Bowtie2 v2.2.9 (LANGMEAD AND SALZBERG 2012) with the parameters *--maxin 900 --no-discard --no-mixed*. Three isolates were removed from further analysis because they had low alignment rates to the E2 reference genome and were likely contaminated. UnifiedGenotyper within the Genome Analysis Toolkit (GATK) version 3.4 (MCKENNA *et al*. 2010) was used to identify SNPs and INDELs with the parameters *-dcov 1000 -glm BOTH*. SNP filtering consisted of the removal of tri- and tetra-allelic SNPs, SNPs with a minor allele frequency ≤ 5% and SNPs that were heterozygous in more than 5% of the isolates. Within individual samples, SNPs with a read depth < 4x were entered as missing data. Considering that the *M. oryzae* genome is haploid, seven isolates with ≥ 1% of heterozygous SNPs were removed from further analyses. All remaining heterozygote SNPs were converted to missing data. Finally, SNP loci with more than 20% of missing data were removed from the data set.

### Strain similarity

To evaluate the genetic similarity between strains, we conducted pairwise comparisons on the filtered SNP data using an in-house script. The percent similarity between any two strains was calculated as (100 * number of identical SNPs between two isolates) / total number of SNPs for which a genotypic score was available in both isolates. Strains with a similarity ≥99.9% were considered clonal.

### Phylogenetic analyses

A subset of 15,838 SNPs was selected to maximize both Modified Roger’s distance (with weight 0.7) and Shannon’s diversity (with weight 0.3) using Core Hunter v2.0 (DE BEUKELAER *et al*. 2012). A Maximum Likelihood (ML) phylogenetic tree of the 166 multilocus haplotypes was generated based on the 15,838 SNP set with 1,000 bootstrap replicates under Mega X (KUMAR *et al*. 2018) using the Tamura-Nei model (TAMURA AND NEI 1993). The ML tree was visualized and color-coded using iTOL 6 (LETUNIC AND BORK 2024). Branches of the tree produced in <50% bootstrap replicates were collapsed.

To determine the phylogenetic placement of the finger millet-infecting *M. oryzae* isolates from eastern Africa in relation to isolates with different host specificities and geographic origins, sequencing data from 52 isolates from 16 host species (**Table S7**), including a representative selection of nine eastern African finger millet-infecting isolates from the current study, were analyzed. For strains sequenced using short-read technologies, low-quality sequences (Phred score <30) were removed using Trimmomatic v0.39 (BOLGER *et al*. 2014). For strains with full genome assemblies (*Lolium perenne* strains LpHO and LpCH2, *Bromus tectorum* strain BtP29 and *Festuca arundinaceum* strain Pg1213), short reads were generated *in silico* by breaking up the assembly into consecutive 150 bp fragments using a random seed position. This process was repeated 10 times to increase the number of sampled regions. Bowtie2 v2.4.2 (LANGMEAD AND SALZBERG 2012) was used to align the preprocessed reads to the single-copy Ascomycota BUSCO genes (ascomycota_odb12.2025-04-11 dataset) (MANNI *et al*. 2021). SNP calling was performed on the BAM file of each genotype with HaplotypeCaller from GATK v3.4.0 (MCKENNA *et al*. 2010), and VCF files were combined using GenotypeGVCFs from GATK. The final VCF file, consisting of 76,529 SNPs from 1415 genes that were present in all 52 isolates, was used as input for the SNPrelate R package (ZHENG *et al*. 2012). The VCF file was converted to GDS format using the snpgdsVCF2GDS() function with the option “biallelic.only“, and the GDS file was used to calculate pairwise identity by state using snpgdsIBS() using the parameters maf = 0.05, missing rate = 0.25. Hierarchical cluster analysis was performed on the dissimilarity matrix using the snpgdsHCluster() function, and snpgdsCutTree() was used to determine subgroups of individuals. The tree file in Newick format was visualized using ITOL v7 (LETUNIC AND BORK 2021).

### Population structure analyses

A population structure analysis of the 226 isolates was conducted on the 15,838 SNP dataset using the admixture model in STRUCTURE v2.3.5 (PRITCHARD *et al*. 2000). The analysis was performed using a burn-in period of 100,000 replications, followed by 100,000 Markov Chain Monte Carlo (MCMC) iterations. The number of putative subpopulations (K) was set to range from one to 10, and 10 runs were conducted for each K-value. The optimal value of K was determined based on the Delta K estimate using Structure Selector (LI AND LIU 2018).

### Genome-wide analysis of deletions associated with TEs

Genomic deletions were defined as a minimum of five consecutive SNPs missing in the genome-wide SNP file (198,454 SNPs), and were determined for each strain using a custom script. The genome locations of the SNPs present immediately adjacent to a deletion on both sides and the first and last SNPs missing in a deletion were recorded. Only deletions larger than 100 bp for which the breakpoints could be delineated to a region less than 3 kb were included in our analyses.

The first 25 bp of the inverted repeat of the *Pot2* element (Genbank accession Z33638), and the first and last 25 bp of the LTRs of the retrotransposons *Grasshopper* (Genbank acc. BK061831), *Fosbury* (AH005360), *Pyret* (AB062507), *Inago2* (AB334125), *Family 1* (**Figure S12A)** and *Family 13* (**Figure S12B)** were used as queries in BLASTN v2.2.26 analyses against the Illumina reads of Ethiopian *M. oryzae* strains using the parameters *-task blastn -reward 2 -penalty -3 -gapopen 6 -gapextend 2 -wordsize 4*. Reads were filtered to only retain those for which the terminal region of the TE was located either at the beginning or the end of a read with the direction indicating that the remainder of the TE was absent from the read (**Figure S13**).

This approach did retain some reads that consisted entirely of LTR-retrotransposon sequence because no differentiation was made between the first 25 bp of the 3’ and 5’ LTR and, similarly, between the last 25 bp of the 3’ and 5’ LTRs (**Figure S13)**. The alignment location of reads with homology at their 5’ or 3’ end to the ends of a TE on the E2 reference genome was obtained from the BAM alignment files generated using Bowtie2 v2.4.5 (LANGMEAD AND SALZBERG 2012). CIGAR (Concise Idiosyncratic Gapped Alignment Report) strings identified reads that transitioned from a matched alignment (sequence present in E2) to alignments with a combination of matched bases, insertions and deletions (sequence adjacent to matched alignment absent from E2) (**Figure S6**). The location of the transition point on the E2 genome assembly was estimated from the CIGAR string. If the transition point was located within a deletion breakpoint region ± 200 bp, the read was retained. A minimum of three reads with a transition point located in a deletion breakpoint was considered evidence that a deletion in an Illumina-sequenced strain was associated with a transposable element. A flow chart of the approach is shown in **Figure S14**, and scripts are available from Github (https://github.com/ysahin1/DevosLab-Pan_genome_fingermillet_infecting_MagnaportheOryzae)

### Comparative genome analyses

The annotated genome of the finger millet-infecting blast isolate MZ5-1-6 (GOMEZ LUCIANO *et al*. 2019) was downloaded from NCBI (GCA_004346965.1). The genomes of the wheat-infecting (B71; GCA_004785725.2; PENG *et al*. 2019) and rice-infecting (P131; GCA_000292605.2; LI *et al*. 2024) blast strains were obtained from the NCBI genome portal (https://www.ncbi.nlm.nih.gov/datasets/genome/) and annotated with the annotation pipeline MAKER v3.01.04 (CANTAREL *et al*. 2008) with default parameters, incorporating reference proteins from *MoE* strains E2 (this study) and MZ5-1-6 (GOMEZ LUCIANO *et al*. 2019). Protein sequences were used as queries in BLASTP searches against E2 primary proteins, retaining top hits with an e-value ≤ 1e^-5^. For syntenic block identification, E2-MZ5-1-6, E2-B71, E2-P131 and B71-P131 pairs were analyzed using MCScanX (WANG *et al*. 2012) with a match score of 50, match size of 5, gap penalty of -1, overlap window of 5, e-value threshold of 1e^-5^, and a maximum gap of 25. Protein pairs located within syntenic blocks were considered orthologs. Syntenic relationships were depicted using Circa v.1.2.2 (OMGenomics Labs).

### Divergence age of blast isolates

To estimate divergence times between *MoE* strains E2 and MZ5-1-6, *MoT* strain B71 and *MoO* strain P131, single-copy gene pairs with a one-to-one relationship between strains were identified using Orthofinder v2.5.5 (EMMS AND KELLY 2019) and the corresponding protein sequences aligned using Clustal Omega v2.1 (SIEVERS *et al*. 2011). The multiple sequence alignments were converted to codon-based DNA alignments using PAL2NAL v14 (SUYAMA *et al*. 2006). The synonymous (Ks) and nonsynonymous (Ka) substitution rates were estimated for each aligned homologous gene pair using the CODEML program from PAML v4.10.5 (YANG 2007) using the Nei and Gojobori method with the parameters CodonFreq = 2, model =0, omega = 0.5. Finally, the divergence time (T) between was estimated from the median Ks value using the formula: T = Ks / (2 * r).

### Pathogenicity assay

Whole-plant infections were performed with 1×10^5^ spores/ml in 0.25% gelatin solution to assess pathogenicity of the blast isolates (VALENT *et al*. 1991). Seven days after inoculation, symptoms on the sprayed plants were scored on a 0 to 5 scale (**Figure S7**), and recorded as described by MATSUNAGA *et al*. (2017). Briefly, the marked youngest and/or second youngest leaf that was expanded at the time of inoculation was scanned with an EPSON perfection 4870 Photo Scanner with 24-bit color and 600 dpi resolution. The diseased area of individual leaves was then measured by ImageJ (SCHNEIDER *et al*. 2012). Pixels were converted to centimeters using the Set Scale command. Color Threshold with HSB color space was used to select and measure appropriate thresholds to whole leaf area and specifically diseased leaf area, respectively.

### Leaf sheath infection assays

For leaf sheaths used for staining or confocal microscopy, the sheath of the second or third youngest leaf of finger millet plants (∼12 days after sowing) was peeled off, excised (∼ 2 cm), and laid horizontally on a support. Sheaths were injected with a spore suspension (1×10^5^ spores/ml in distilled water) harvested from ∼10-day old cultures, and incubated at 25 °C until observation. For transcriptome analyses, to minimize sampling time, sheaths were pretrimmed (0.3 to 0.5 cm wide), and taped to the bottom of a 5 cm petri dish. Two leaf sheaths were generated from each plant (both sides) and six leaf sheaths were placed per petri dish. A total of 15 plants were used for each treatment at each time point. Sheaths were flooded with 10 ml of inoculum consisting of 1×10^5^ spores/ml and incubated at 25 °C until sampling. Control sheaths were incubated with distilled water. At 30 or 48 hpi, the inoculated sheaths were gently swabbed to remove spores, appressoria and mycelia on the sheath surface, and immediately analyzed for infection density using confocal microscopy or frozen in liquid nitrogen and stored at -80 °C for transcriptome analysis.

### Confocal microscopic analyses of pathogen infection

Fluorescein diacetate (FDA) and propidium iodide (PI) staining of finger millet sheaths were performed as described (JONES *et al*. 2016). Confocal microscopy was carried out on a Zeiss LSM 880 Confocal Microscope with an upright microscope stand. Excitation/emission wavelengths were 488 nm/496 to 544 nm for EGFP/FDA and 543 nm/565 to 617 nm for mCherry/PI to assess host cell viability and *M. oryzae* effector localization. Confocal images were processed with Zen Black software (version 10.0, Zeiss). Infection areas (%) were quantified using ImageJ from confocal images of finger millet sheaths infected with EGFP-labeled *M. orzyae* strain CKF4046 at 30 hpi and 48 hpi.

### Sequencing and analyses of the fungal transcriptome

Mycelia of strain CKF4046 (EGFP-labeled E2 strain), grown in liquid complete medium in the dark for four days (three biological replicates), and leaf sheaths of finger millet accessions TZA1637 and AAUFM-44 at 30 hpi and 48 hpi following infection with strain CKF4046 (three biological replicates, each consisting of 30 leaf sheaths) were collected and flash frozen in liquid nitrogen.

Total RNA was extracted from ground tissue with Trizol followed by purification using the RNA Clean and Concentrator-5 kit (Zymo) according to the manufacturer’s instructions for three biological replicates per sample. Genomic DNA was removed using the Turbo DNA-free kit (Invitrogen). Barcoded RNA-seq libraries were generated with the KAPA Stranded mRNA-Seq Kit (Kapa Biosystems) using 1.5 µg of total RNA as input. The manufacturer’s protocol was followed using half reaction volumes through step 10.13 and full reaction volumes from step 10.14 to the end of the protocol. Fragmentation was carried out at 94 °C for eight minutes, and 11 library amplification steps were used. Library concentrations were quantified using a Qubit Fluorometer and a Qubit 1X dsDNA High Sensitivity Assay Kit (Invitrogen). The size distribution of randomly selected libraries was assessed on a Fragment Analyzer^TM^ Automated CE System at the Georgia Genomics and Bioinformatics Core (GGBC). The barcoded libraries were combined in equal amounts except for the mycelia libraries for which only 50% quantities were added to the pool. The pooled libraries were sequenced on an Illumina NextSeq2000 platform (75 bp paired-end reads) at the GGBC.

The raw sequencing reads were split by barcode and trimmed to remove adapter sequences and low-quality sequences (Phred score < 33) using the paired-end mode of Trim Galore v0.6.0 (https://github.com/FelixKrueger/TrimGalore). The trimmed reads were aligned to the E2 reference genome using HISAT2 v2.2.1 (KIM *et al*. 2019a) with default parameters. The aligned reads were assembled into transcripts using StringTie v2.2.1 (PERTEA *et al*. 2015). The assembled transcripts were merged across samples and used as a reference for transcript quantification. The resulting GTF files with quantified transcripts were converted to a gene count matrix using prepDE.py (http://ccb.jhu.edu/software/stringtie/index.shtml?t=manual). Lowly expressed genes (< 10 reads across all samples) were removed, and the remaining genes were then analyzed for differential expression using DESeq2 v1.42.0 (LOVE *et al*. 2014) in R v4.3.2. Pairwise analyses were conducted using a single main effect to identify differentially expressed genes for each treatment (host accession and time).

Gene onthology (GO) enrichment was conducted with gProfiler version e113_eg59_p19_f6a03c19 (https://biit.cs.ut.ee/gprofiler/gost; KOLBERG *et al*. 2023) using the *M. oyzae* strain 70-15 gene IDs homologous to the finger millet genes (best BLASTN hit at evalue threshold of 1e^-5^) as input, all genes expressed (average normalized read count ≥5) in at least one of the infection conditions as background, and the g:SCS algorithm for multiple testing correction. Statistically significant driver terms were downloaded, and results plotted using ggplot2 in R (WICKHAM 2011).

Co-expression analysis was conducted using the *co-seq* R package v1.26.0 (RAU AND MAUGIS-RABUSSEAU 2018) using default model and data transformation parameters (*K-*means algorithm and log centered log transformation) and the DESeq2 size factors to normalize read counts for expressed genes (gene count ≥ 10 across all samples).

Identification of biosynthetic gene clusters (BGCs) in the E2 genome was conducted by Fungi AntiSMASH v7.1.0 (MEDEMA *et al*. 2011) using default parameters.

## Supporting information

all supplemental data

## Data availability

E2 genome assembly: JBPPCE000000000; Illumina sequence of 226 *MoE* strains and RNA-seq reads: PRJNA1279063; Scripts are available from Github (https://github.com/ysahin1/DevosLab-Pan_genome_fingermillet_infecting_MagnaportheOryzae); All other data are provided as Supplementary Data.

## Author contributions

SdV designed and oversaw the collection of blast-infected finger millet tissues. Collections were made by MMD, TA, KT, JHR and JT. DWK conducted *MoE* isolation from the infected tissues. SS, SM and TAS provided the historical isolates and conducted the mating type analyses. PQ conducted DNA extractions, generated the short-read assemblies, conducted gene annotation and SNP calling, determined presence/absence of genes and strain similarity of the 226 strains, and participated in the comparative analyses of strains with different host specificities and in the transcriptome analyses. JG constructed the PacBio and Illumina libraries and conducted the sequencing. JJ generated the PacBio assembly. HW conducted the repeat analyses. JZ performed the leaf sheath assays, pathogenicity assays, confocal microscopy and RNA extractions. HCW conducted the transcriptome analyses. BAB, YS and JMA conducted the phylogenetic and population genetic analyses. KAP conducted the infection assays and quantitative analysis of infection. BM performed the confocal microscopy. YS participated in the comparative analyses, analyzed the divergence age of blast isolates, and conducted the genome-level deletion analyses. K-TK and Y-HL conducted effector gene mining. THP conducted the statistical analyses. CHK and KMD designed and coordinated the study, assisted with data analysis and interpretation, and drafted the manuscript with input from the co-authors. All authors reviewed and approved the manuscript.

