## Supplementary material for "Pan-genome analyses of 226 finger millet-infecting *Magnaporthe oryzae* strains from eastern Africa": all supplemental data

**Supplementary Notes**

**Note S1**: **Estimating transposable element copy numbers from short-read assemblies**

Although we cannot accurately determine the copy number of transposable elements in short-read assemblies because repeats are likely collapsed or lacking, the assumption is that the distal 5’ and 3’ regions of a TE are present in the assemblies, in particular if the TEs are inserted in low copy DNA. Therefore, the number of BLASTN hits ≥ 25 bp and with ≥ 95% similarity that cover each base of the TE using the *semi-de novo* assemblies as queries provides an estimate of the TE copy number.

**Note S2**: **The closely related LTR-retrotransposons *MAGGY* and *Fosbury***

The LTR-retrotransposons *MAGGY^2^* and *Fosbury^3^* are 99% similar across LTRs and, throughout the paper, we refer to the element as *Fosbury*.

**Note S3**: **Comparative analyses between *MoE, MoT* and *MoO* strains reveal insertion hot spots**

Insertion hot spots are exemplified by insertions (relative to E2) in Chr5 and Chr6 that differed in origin in MZ5-1-6, B71 and P131 (**Figure 3**). The region on Chr5 in E2 (location 2,193,241 – 2,460,482) is colinear with a region on Chr3 in MZ5-1-6, a different Chr3 region in B71, and a Chr6 region in P131. Similarly, the region on Chr6 in E2 (location 2,657,339 – 3,200,001) corresponds to regions on Chr2 in MZ5-1-6, and non-syntenic Chr1 regions in B71 and P131.

**Note S4**: **Comparative analyses of the repeat content in representative *MoE, MoT* and *MoO* strains**

*Grasshopper* (*Grh*) is the highest copy number element present in *MoE* strains E2 and MZ5-1-6 (**Table S8**), and is of recent origin (identical LTRs in 120/121 full length elements; **Data S3**). The highest copy number transposable element in the *MoT* strain B71 is an uncharacterized LTR-retrotransposon (*Family 13*) with nine full-length copies (**Table S8**), eight (89%) of which have identical LTRs and one that is estimated to have inserted around 63,000 years ago (**Data S3**). *Family 13* was also the second most prevalent element in the *MoE* isolates E2 and MZ5-1-6 with 83% (39/47) of the full-length elements having identical LTRs while the others had estimated insertion dates ranging from 63,000 to 816,000 years ago. The highest copy number LTR-retrotransposon in the rice blast strain P131 is *Fosbury* with 78 full-length copies (**Table S8**). Around 90% of the full-length elements had identical LTRs, and the oldest transposition events were dated to around 127,000 years ago (**Data S3**). Neither full-length elements nor solo LTRs of *Fosbury* were identified in E2, MZ5-1-6 or B71 (**Table S8**) although *Fosbury*, or a related element, was present in the majority of ET strains originating from the West Gojam (88.2% of isolates), Awi (100%) and East Wollega (100%) regions (**Data S1**). The absence of *Fosbury* from the E2 reference strain and lack of full-length element assemblies for other Ethiopian strains precluded dating of the divergence of the *Fosbury* elements.

**Note S5**: **Analysis of chromosomal deletions in Ethiopian *MoE* strains**

To determine the mechanism(s) by which chromosomal deletions in *M. oryzae* occurred, the precise breakpoints and flanking sequences from a small set (n=14) of randomly selected deletions in Ethiopian strains belonging to the ET population were manually extracted from the Illumina read alignments against the E2 reference genome. The analysis was limited to Ethiopian strains because of the need for high sequence similarity to the E2 reference strain in order to identify reads that span a deletion breakpoint. A locally high SNP density adjacent to a deletion is indicative of reads that extend into a region that is present in E2, but is deleted in the resequenced strain (**Figure S6**). This characteristic was exploited to conduct genome-wide analyses in Ethiopian ET strains of deletions associated with transposable elements.

**Note S6**: **Presence/absence polymorphisms of non-effector genes**

The level of loss observed for non-effector genes amounted to 8.6% (1030/11,937 genes for which presence/absence could be established unambiguously in ≥ 80% of strains; 11,447 annotated in E2 and 490 in the 225 short-read assemblies) (**Data S7**). While loss of non-effector genes was highest in KU strains with 18.2% of gene absences (187/1030) being strongly biased towards the KU clade compared to 12.6% of gene absences (130/1030) being strongly biased towards the ET clade (**Figure N6**), this difference was far less pronounced than what was observed for effector genes. The reduced bias is also demonstrated by the comparable numbers of non-effector genes that are present or absent (75% threshold) within each clade. In the ET clade, 47.0% of genes (484/1030) were present in at least 75% of isolates and 435/1030 (42.2%) were absent. Presence and absence in the KU clade amounted to 55.9% (576/1030) and 48.3% (497/1030), respectively.

Figure N6. (A) Heatmap (generated with Heatmapper ^1^) showing the relative presence of non-effector genes with the most intense green representing genes that are present in all strains within a country/population and the most intense purple representing genes that are absent in all strains within a country/population. (B) Graph showing the average frequency of non-effector gene presence by country/population. Significance was determined by Tukey corrected multiple comparison tests and identified as different letters. Genes that are uniformly present were not included. T_KU, K_KU, and U_KU are Tanzanian, Kenyan and Ugandan blast isolates that belong to the KU population; E_ET and T_ET are Ethiopian and Tanzanian blast isolates that belong to the ET population. For clonal isolates, only a single representative isolate was included in the analyses.

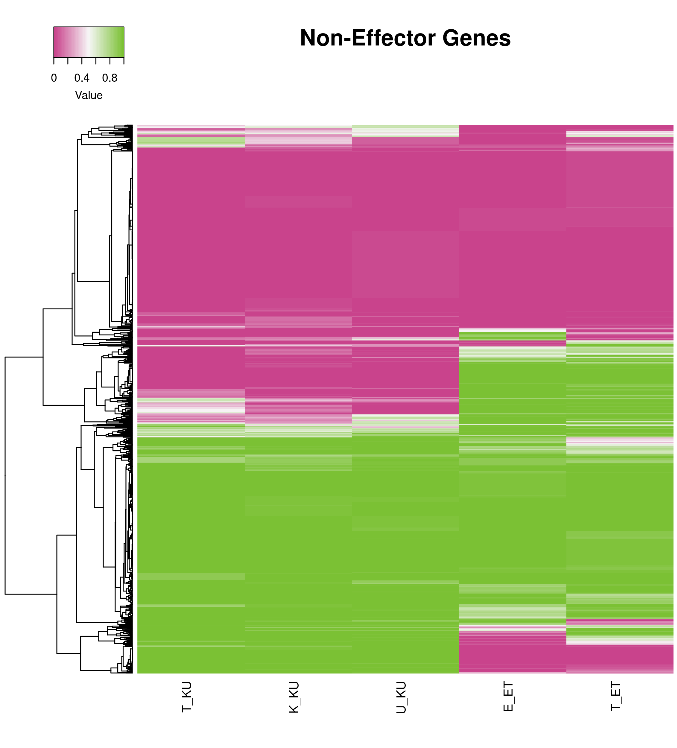

**Non-Effector Genes**

A

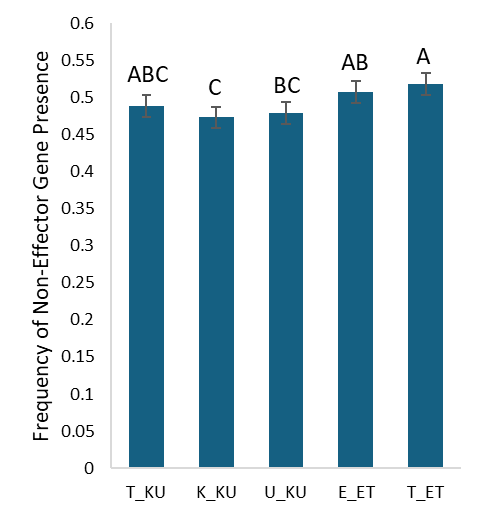

B

Non-effector genes with presence/absence polymorphisms are significantly smaller than genes that are uniformly present across the strains (average/median gene size of 930/632 bp *vs.* 1804/1533 bp for genes annotated in E2). It is possible that some are effector genes (average/median size of 702/516 bp) that were missed by the prediction pipeline which has a false-negative rate of 33% based on known rice effectors. The non-effectors with presence/absence variation also had a paralog (as determined by orthofinder) at a higher frequency than genes that were uniformly present (5.0% *vs.* 1.3%), a difference that was not seen for effector genes (2.3% *vs.* 3.0%), suggesting gene loss following duplication. Single copy genes that are lost in select strains may be non-essential, provide a selective advantage when lost, or could have structurally different but functionally redundant gene copies ^4,5^.

**Note S7**: **Finger millet leaf sheath assay**

We developed a finger millet leaf sheath infection assay by modifying the rice sheath infection assay ^6^. Similar to rice sheaths, finger millet sheaths are optically clear, facilitating direct observation of the fungal infection process under microscopy. Infected leaf sheaths also have a higher ratio of fungal to plant transcripts compared to infected leaves originating from whole-plant spray inoculations. First, we confirmed that in whole-plant spray inoculation, the *M. oryzae* transformant CKF4046, expressing cytoplasmic enhanced green fluorescent protein (EGFP), retains the virulence of its parental wild-type E2 strain on finger millet accessions AAUFM-44 (susceptible) and TZA1637 (resistant) (**Figure N7.1**).

E2

CKF4046

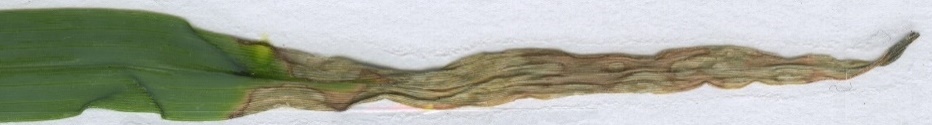

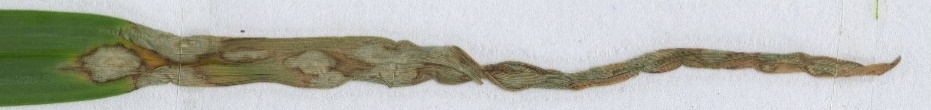

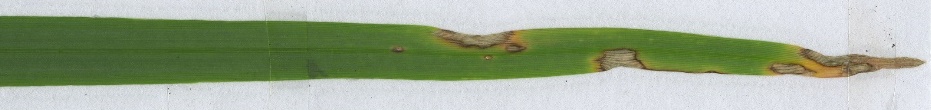

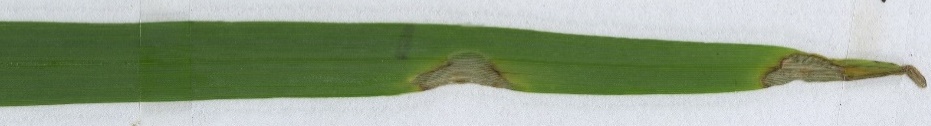

AAUFM-44

AAUFM-44

TZA1637

TZA1637

***Figure N7.1.*** *Consistent infection levels of finger millet accessions AAUFM-44 (susceptible) and TZA1637 (resistant) following whole-plant spray inoculation with the wild-type strain E2 and its EGFP transformant CKF4046 at 7 days post-inoculation.*

Comparative transcriptomic analyses were conducted using M. oryzae strain CKF4046 on leaf sheaths of finger millet accessions AAUFM-44 and TZA1637 at 30 hours post inoculation (hpi) and 48 hpi. Before harvesting infected sheaths for RNA extraction, we conducted live-cell confocal microscopy to quantify fungal colonization of sheath cells. Two confocal images for each time point (30 and 48 hpi) and interaction (AAUFM-44 and TZA1637) were analyzed using ImageJ to quantify infection area. Representative confocal and ImageJ selection images are shown in **Figure N7.2.** See **Figure 8** for quantification of infection area along with representative confocal images of CKF4046-infected AAUFM-44 and TZA1637 at 48 hpi.

***Figure N7.2.*** *Green channel and ImageJ selection images of resistant and susceptible finger millet sheaths inoculated with CKF4046 conidia, expressing cytoplasmic EGFP. Arrowheads indicate some invasive hyphae in green, while asterisks mark areas of autofluorescence that were manually excluded during ImageJ analysis. Panel A (top left) shows the distribution of invasive hyphae on a section of AAUFM-44 sheath at 30 hpi. The top right image in Panel A is the same green channel image processed in ImageJ to color-select the green hyphae, which are outlined in yellow. The bottom left image in Panel A shows an AAUFM-44 sheath at 48 hpi, where green hyphae have expanded and spread through the sheath cells. The bottom right image in Panel A is the ImageJ selection of the 48 hpi AAUFM-44 sheath. Panel B (top left) shows a 30 hpi green channel image of a TZA1637 sheath with very little green signal due to the lack of invasive hyphae; note the two bands of autofluorescence marked with asterisks. The top right image in Panel B is the ImageJ selection of the green signal in that image, manually adjusted to exclude the autofluorescent bands. The bottom left image in Panel B shows a TZA1637 sheath at 48 hpi, with some branched hyphal growth localized in vein-associated cells and additional autofluorescence marked with asterisks. The bottom right image in Panel B is the ImageJ selection of the 48 hpi image, with autofluorescence noise excluded in quantitation. Bars = 100 μm.*

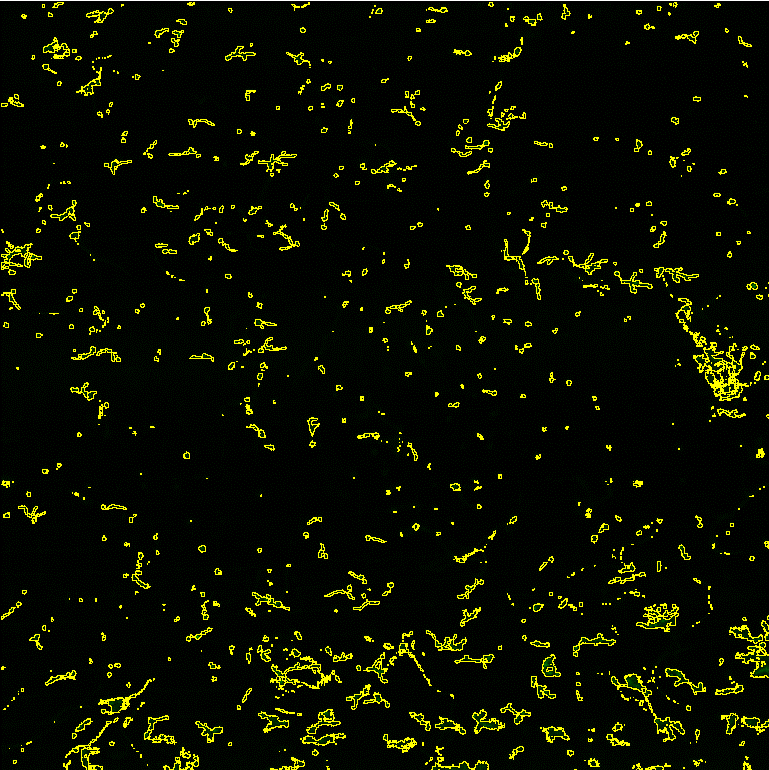

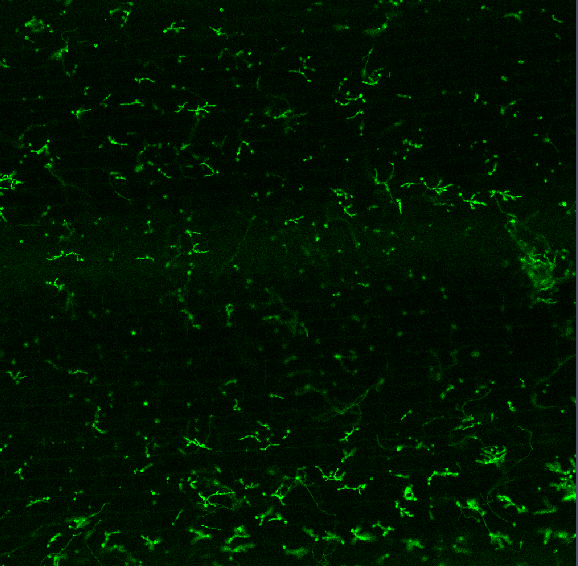

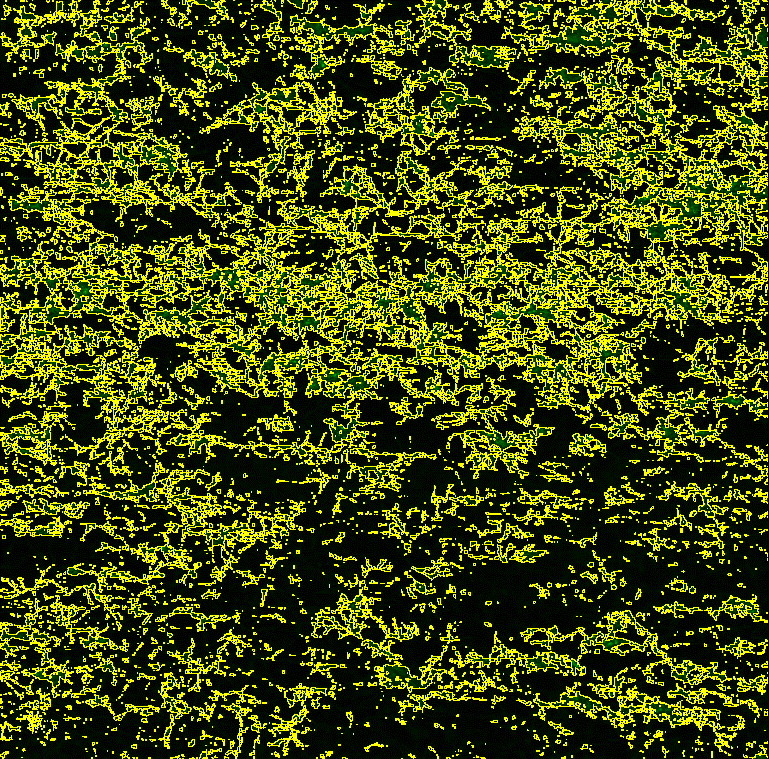

GFP Channel

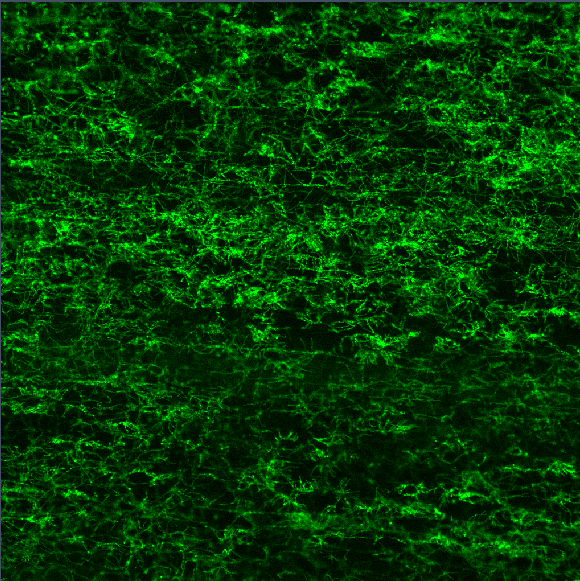

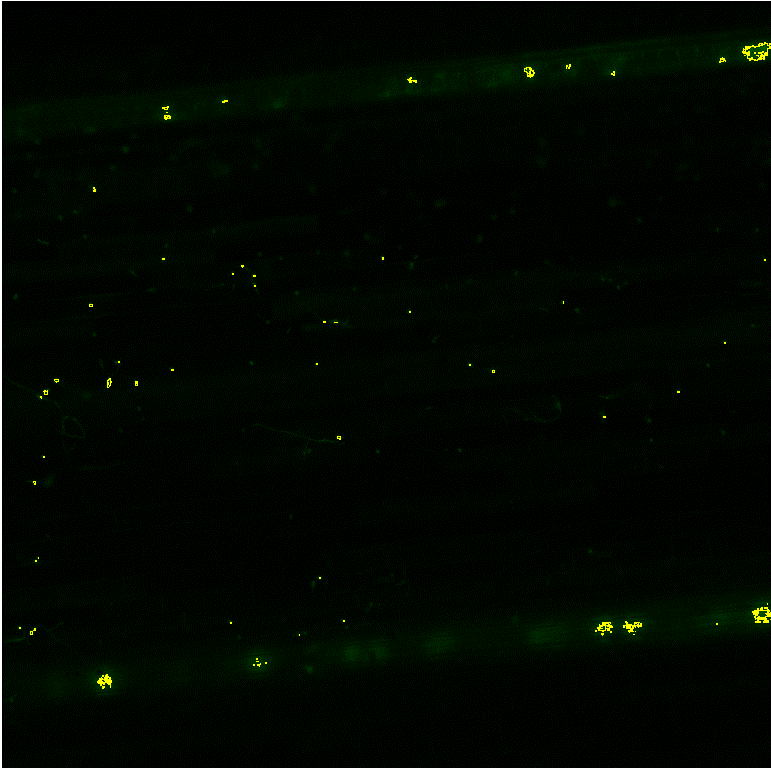

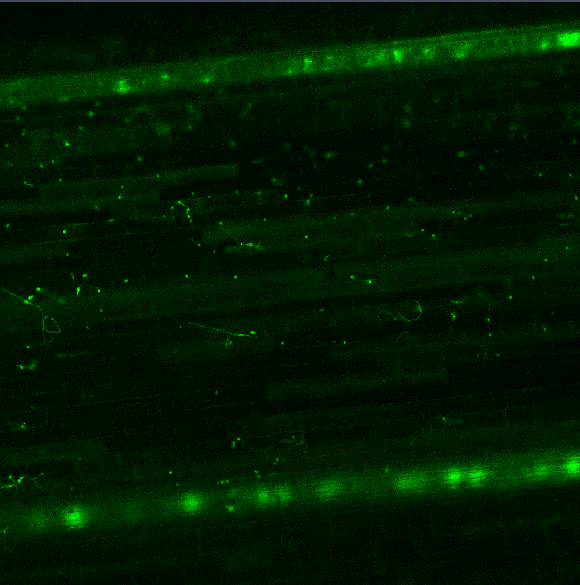

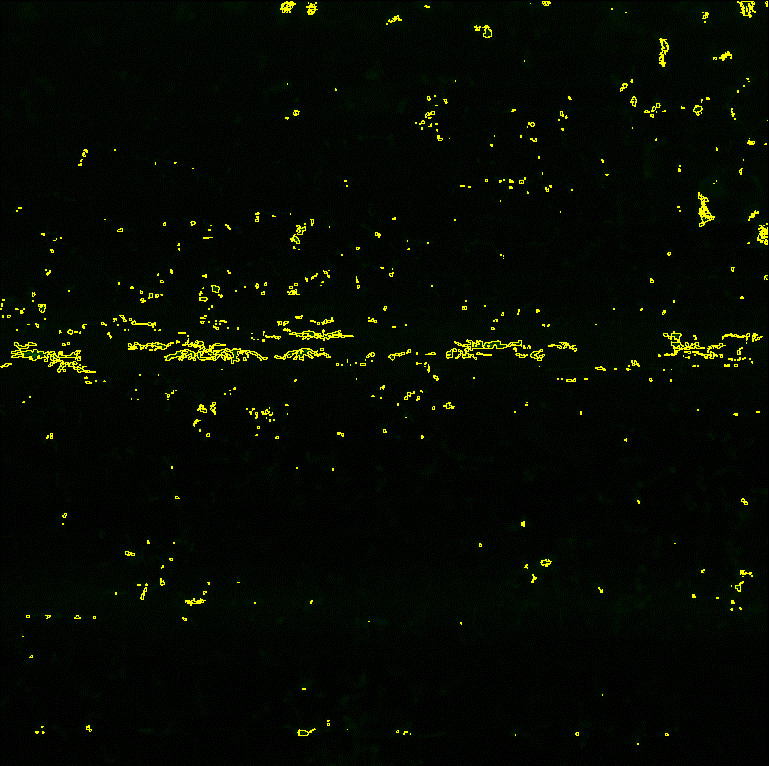

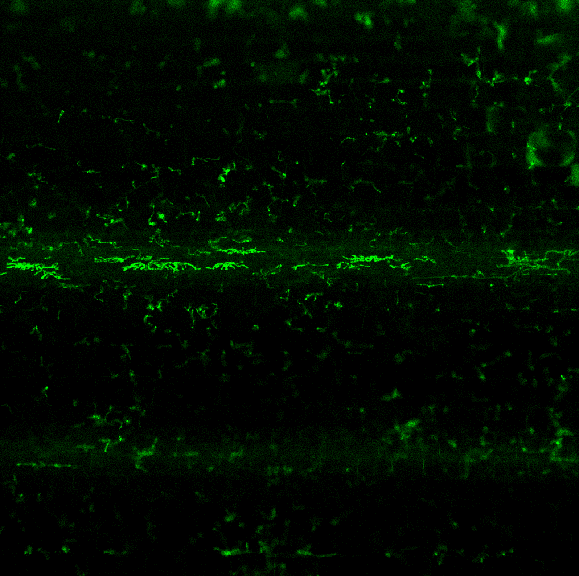

A

B

ImageJ Selection

30hpi

30hpi

30hpi

30hpi

48hpi

48hpi

48hpi

48hpi

*

*

*

GFP Channel

ImageJ Selection

**AAUFM-44**

**TZA1637**

Live-cell confocal microscopy demonstrated the viability of finger millet sheath cells colonized by the *M. oryzae* strain CKF4046 at 30 hpi ^7^ (**Figure N7.3**). We further used the *M. oryzae* E2-transformant CKF4196, expressing PWL2:mCherry:NLS and BAS4:GFP, to show the formation of the extra-invasive hyphal membrane (EIHM) and the biotrophic interfacial complex (BIC) in *M. oryzae*-infected finger millet cells (n = 21 infected cells) (**Figure N7.3**). Both BIC and EIHM are plant-derived structures associated with biotrophic invasion and implicated in effector trafficking, well documented in rice cells infected by rice-adapted *M. oryzae* strains ^6,8^. Thus, the successful formation of BIC and EIHM in viable invaded finger millet cells suggests the establishment of biotrophic invasion and effector delivery by *M. oryzae* on finger millet at the early infection stage.

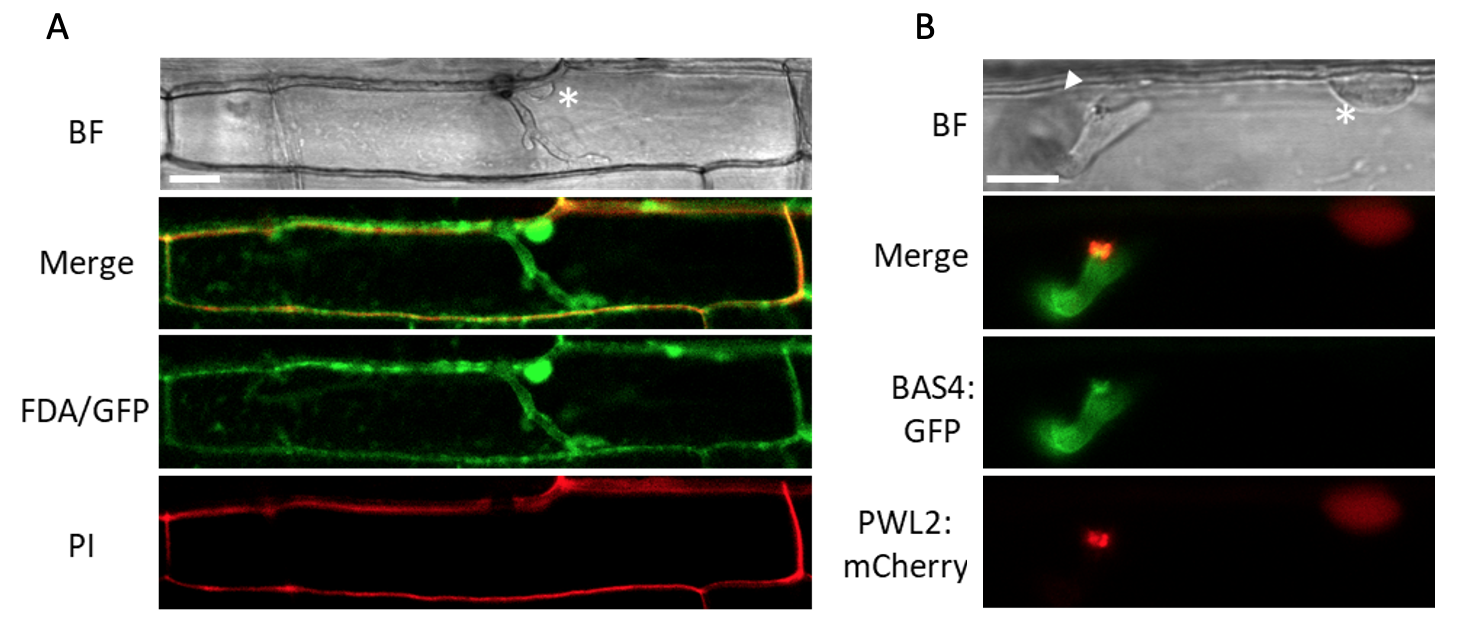

***Figure N7.3.*** *(A) Confocal image of a susceptible finger millet sheath (AAUFM-44) epidermal cell infected with M. oryzae transformant CKF4046 expressing cytoplasmic EGFP (shown in green inside the plant cell) at 30 hpi and stained with fluorescein diacetate (FDA) (green around the plant cell and invasive hypha) and propidium iodide (PI) (red around the plant cell). The asterisk indicates the plant nucleus stained with FDA. Bar = 20 μm (B) Cellular localization of PWL2:mCherry:NLS (red) and BAS4:GFP (green) expressed from M. oryzae strain CKF4196, colonizing a finger millet (AAUFM-44) epidermal cell at 30 hpi. BAS4:GFP showed an outline pattern around the invasive hypha, indicating the presence of EIHM, as in rice cells. PWL2:mCherry:NLS was observed accumulating preferentially at the BIC and translocating into the plant nucleus as indicated by asterisk. The arrowhead indicates the appressorium. Bar = 10 μm.*

**Note S8: BGC 5.01**

AntiSMASH identified cluster 5.01 as a type I polyketide synthase based on the presence of EcMO5g00088770, a reducing polyketide synthase. Interestingly, the cluster also contains three genes encoding proteins with homology to ent-kaurene synthase, gibberellin cluster GA14 synthase and geranylgeranyl pyrophosphate synthase 2 of *Fusarium fujikuroi* that are potentially involved in the gibberellic acid biosynthesis pathway ^9^. The four remaining genes in the cluster encode a cytochrome P450 monooxygenase, two short chain dehydrogenase/reductases and a putative transcription factor with a GAL4-like Zn_2_Cys_6_ binuclear cluster DNA-binding domain (motif CysX_2_CysX_6_CysX_6_CysX_2_CysX_6_Cys) (**Data S14**).

**Note S9**: **BGC 5.12**

Cluster boundaries are difficult to predict ^10^. Although the bioinformatically predicted cluster 5.12 comprises 14 genes, the expression profile suggests that 5.12 consists of two clusters, with the six functionally annotated genes EcMO5g00103950 – EcMO5g00104000 making up one BGC. The six genes in this region encode an ent-kaurene synthase, gibberellin cluster GA14 synthase and geranylgeranyl pyrophosphate synthase 2, in addition to a cytochrome P450 monooxygenase, a nonribosomal peptide synthetase and an uncharacterized protein (**Data S14)**

**Note S10**: **Top-two most significantly upregulated genes in clusters 3, 11 and 13**

A role in virulence of non-effector proteins is exemplified by the functions of the top-two most significantly upregulated genes at 30 hpi with functional annotations in clusters 3, 11 and 13 that were also significantly upregulated at 48 hpi (**Data S12**). Salicylate hydroxylase (EcMO6g00121820) and, potentially, 2,3-dihydroxybenzoic acid decarboxylase (EcMO6g00121630) (cluster 3) may be involved in the catabolism of host salicylic acid, an important defense signaling molecule ^11-13^. Secretory lipases (EcMO1g00013860) (cluster 13) can assist the pathogen with penetration of the cuticle by degrading plant cutin and waxes ^14^. *Ent*-kaurene oxidase (EcMO2g00029440) (cluster 13) converts *ent*-kaurene to *ent*-kaurenoic acid and is part of the gibberellic acid pathway, but may also react with related diterpenes to produce specialized diterpenoids ^15^. Fungal terpenoids can contribute to virulence through toxicity or manipulation of host defense responses ^16^. Pisatin demethylase (EcMO3g00066070) (cluster 11) is a cytochrome P450 family 57A1 member. In the pathogenic fungus *Nectria haematococca*, pisatin demethylase detoxifies the phytoalexin pisatin produced by the host in response to infection ^17^. We hypothesize that the CYP57A1 produced by *M. oryzae* similarly modulates the finger millet defense response by detoxifying host phytoalexins. P-glycoprotein ABC transporters (EcMO4g00086900) (cluster 11) export a variety of substrates, including toxins. Plant pathogens may use ABC transporters to remove plant defense factors. ABC1 in *N. haematococca*, for example, can transport pisatin out of fungal cells which, combined with the detoxification of pisatin by pisatin demethylase, confers tolerance to this phytoalexin.

**Supplementary Figures**

**Figure S1**. Ascomycota BUSCO genes identified in the E2 genome assembly (‘Assembly’) and among the annotated genes with cDNA length ≥ 200 bp (‘Annotation’). S = complete & single copy; D = complete and duplicated; F = fragmented; M = missing

Ascomycota BUSCO representation

|  | S | D | F | M |
| --- | --- | --- | --- | --- |
| Annotation | 1582 | 74 | 9 | 41 |
| Assembly | 1653 | 14 | 10 | 29 |

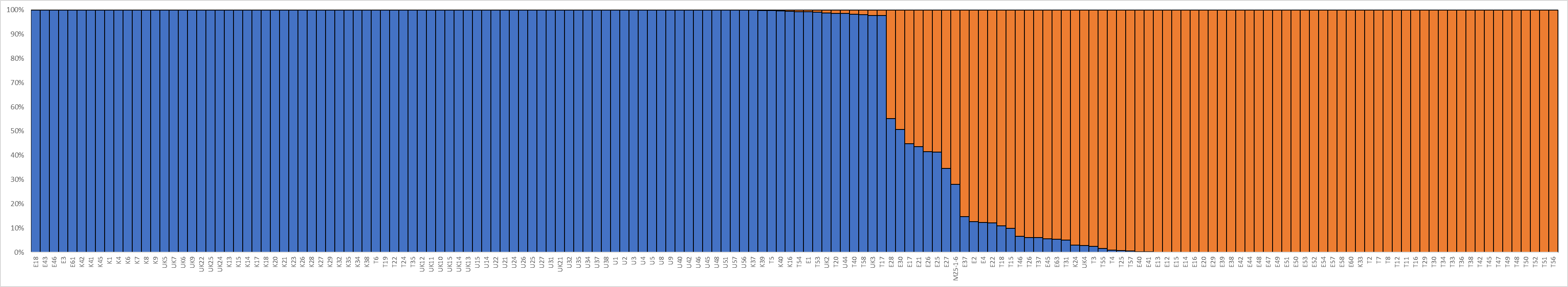

*************

**Figure S2**. Population structure analysis (K=2) of 226 finger millet-infecting *M. oryzae* isolates from eastern Africa. K = strains collected from Kenya during 2015-2017; U = Uganda (2015-2017); E = Ethiopia (2015-2017); T = Tanzania (2015-2017); UK = strains collected from Eastern Uganda/Western Kenya during 2000-2002. The horizontal axis gives the strain IDs ordered by population membership. The vertical axis gives the percentage membership to a single subpopulation for each strain. Admixed strains (<90% membership to a single population) are indicated with an asterisk (*).

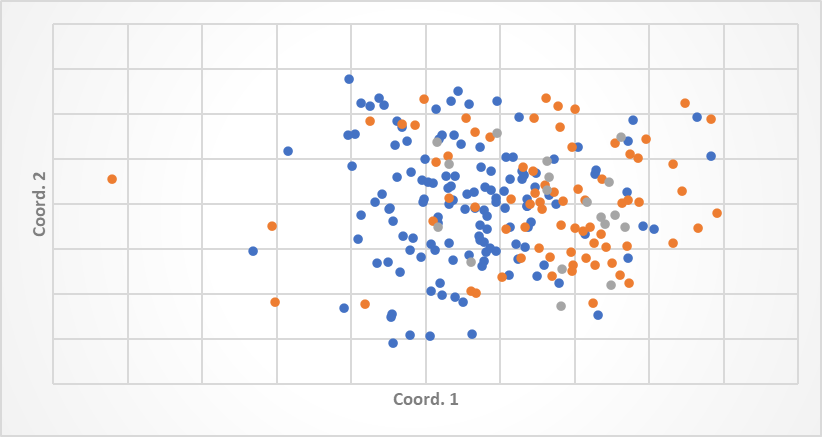

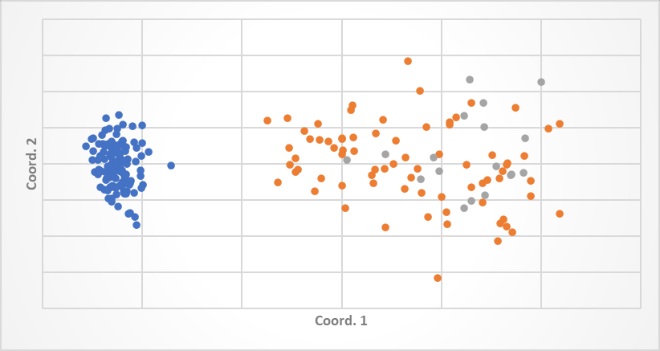

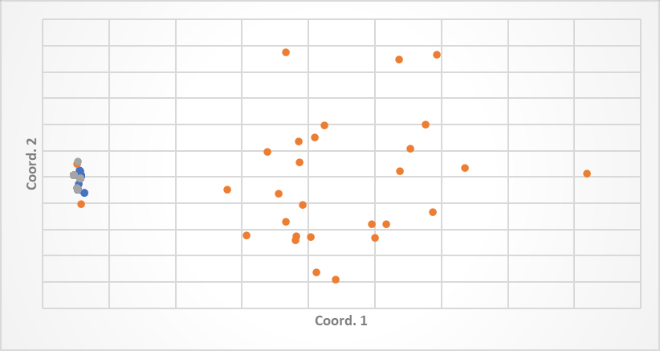

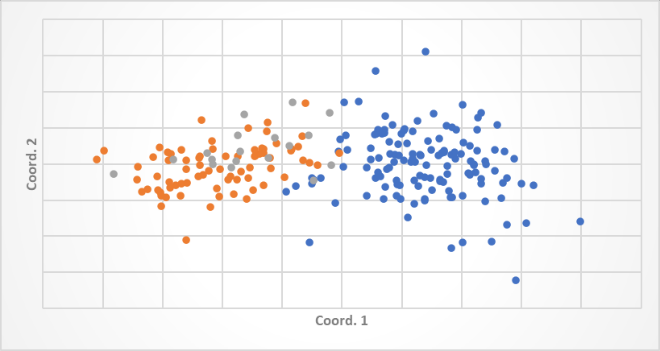

*Family 13*
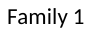
*3*

*Family 1*

*Grasshopper*

*Fosbury*

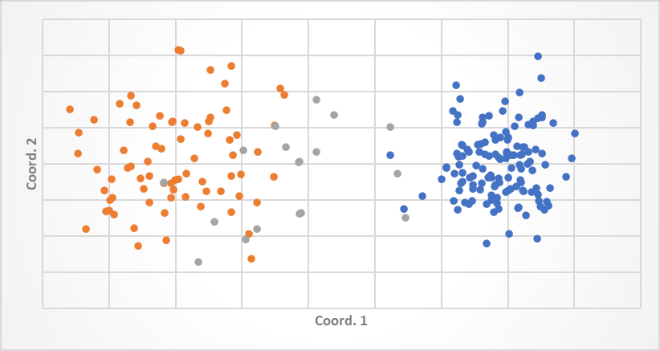

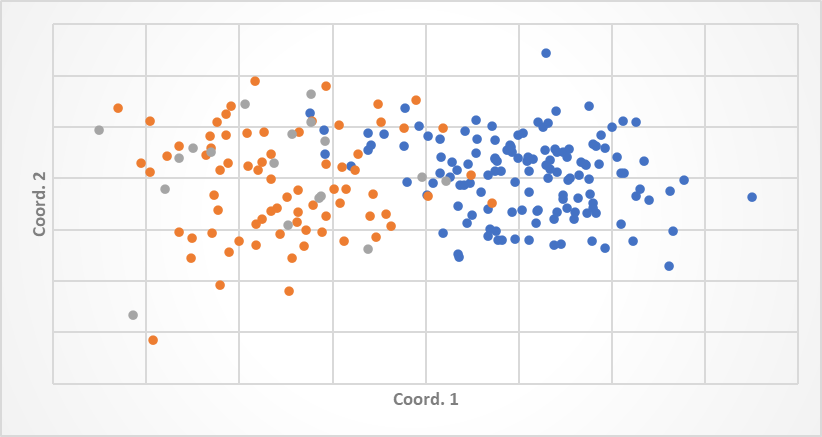

*Pyret*
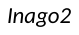

*Inago2*
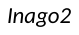

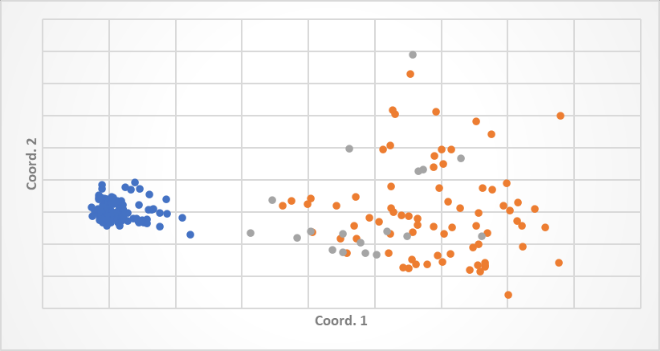

**Figure S3**. PCoA of relative copy number of select transposable elements. KU strains are in blue, ET strains in orange, and Admixed strains in grey.

*Pot2*
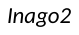

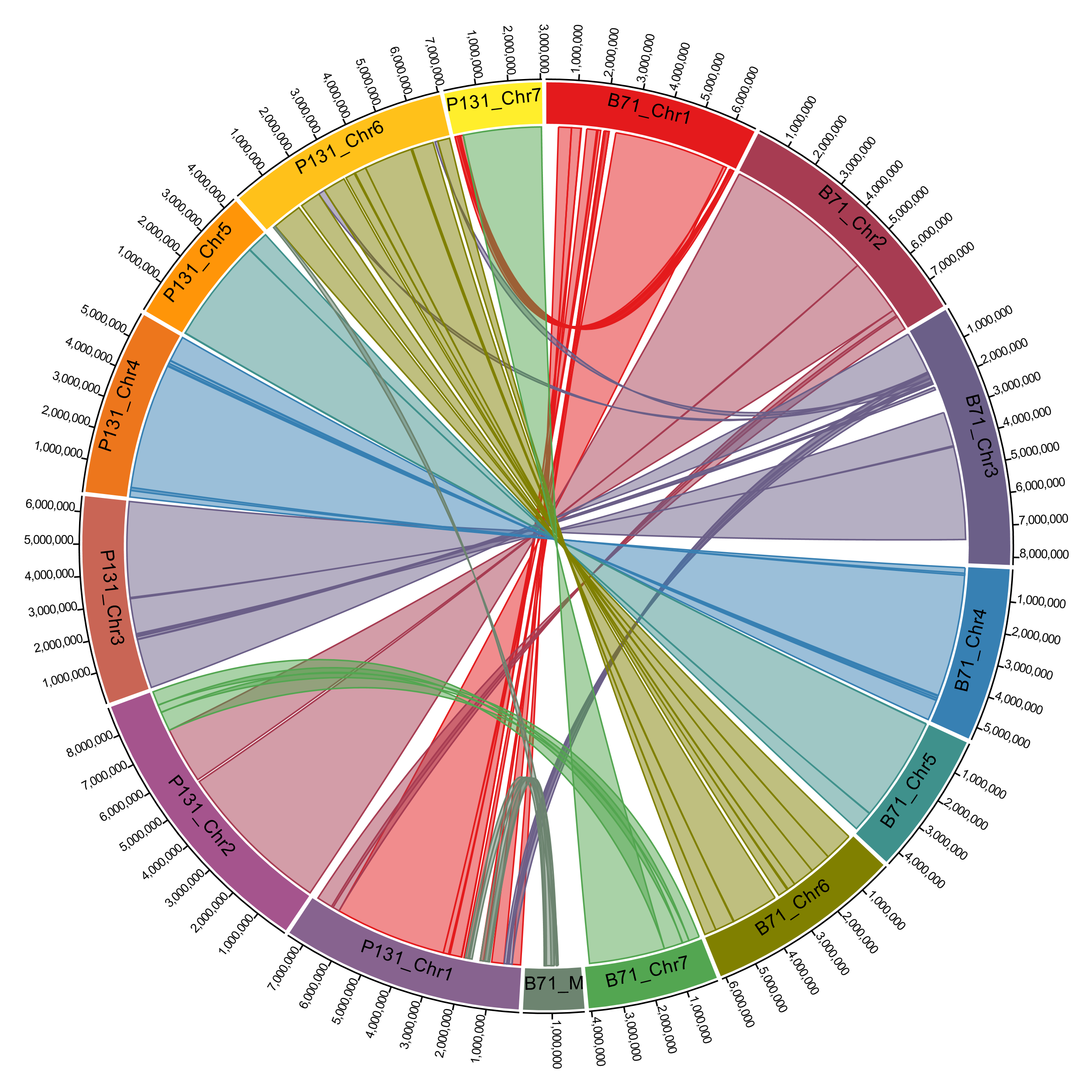

**Figure S4**. Comparative relationship between wheat-infecting isolate B71 and rice-infecting isolate P131.

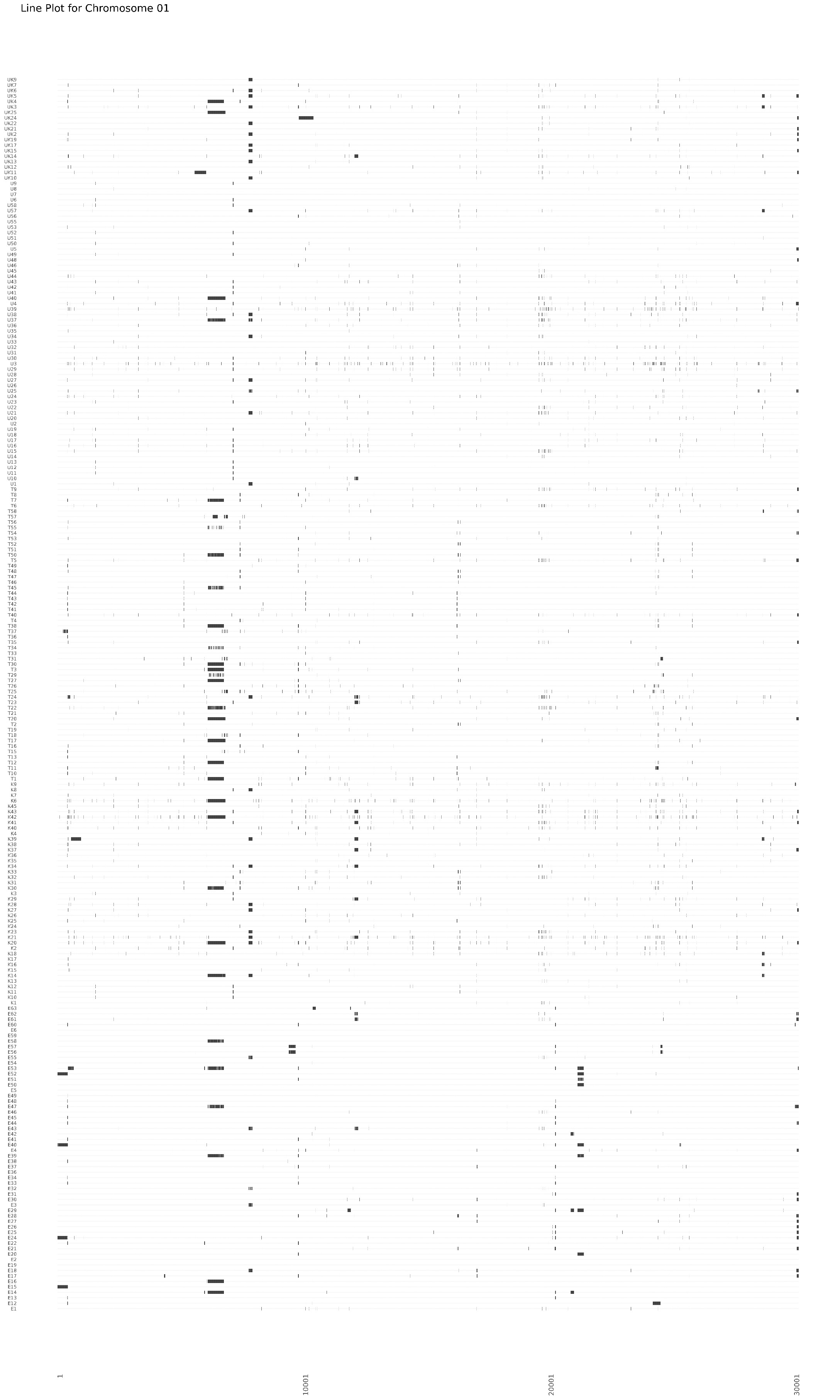

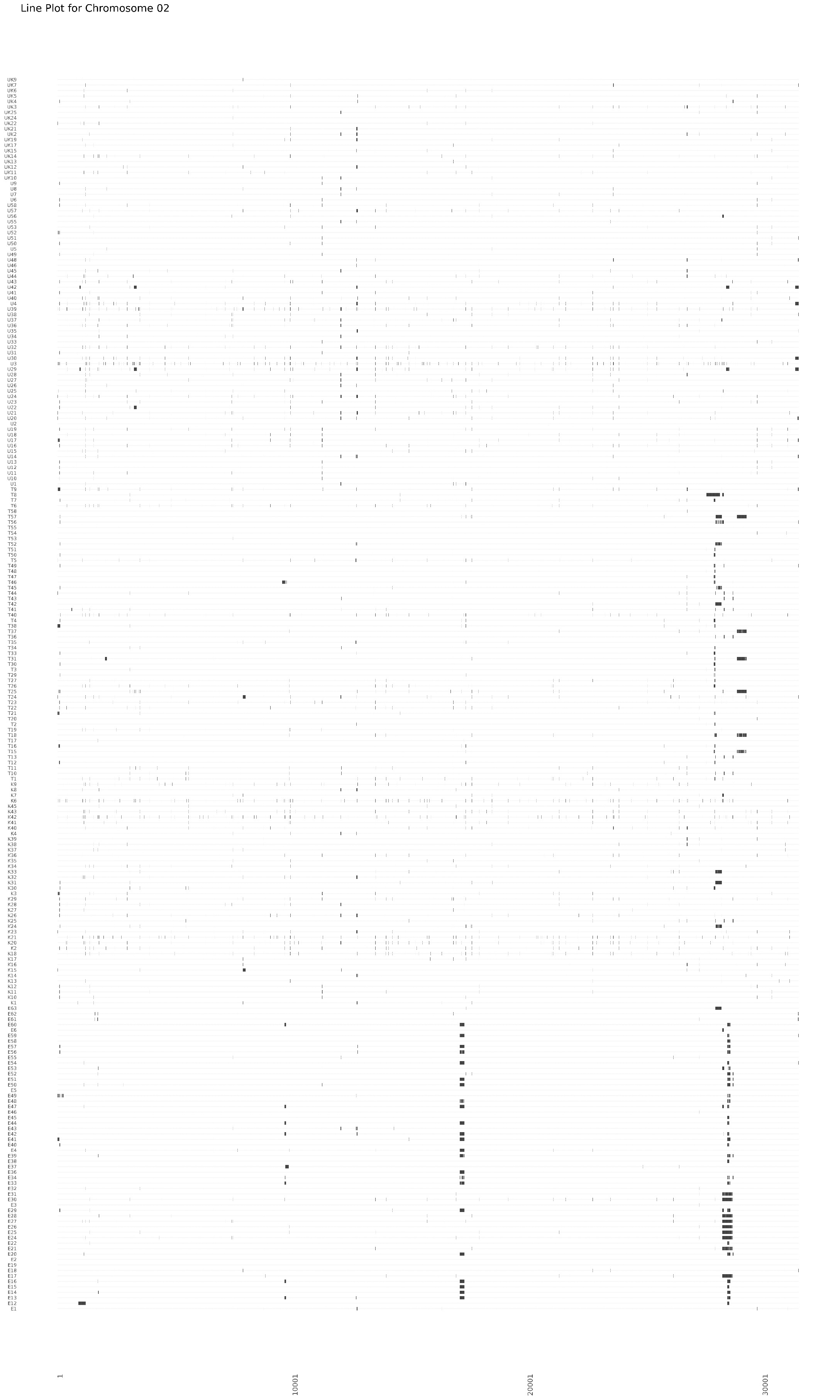

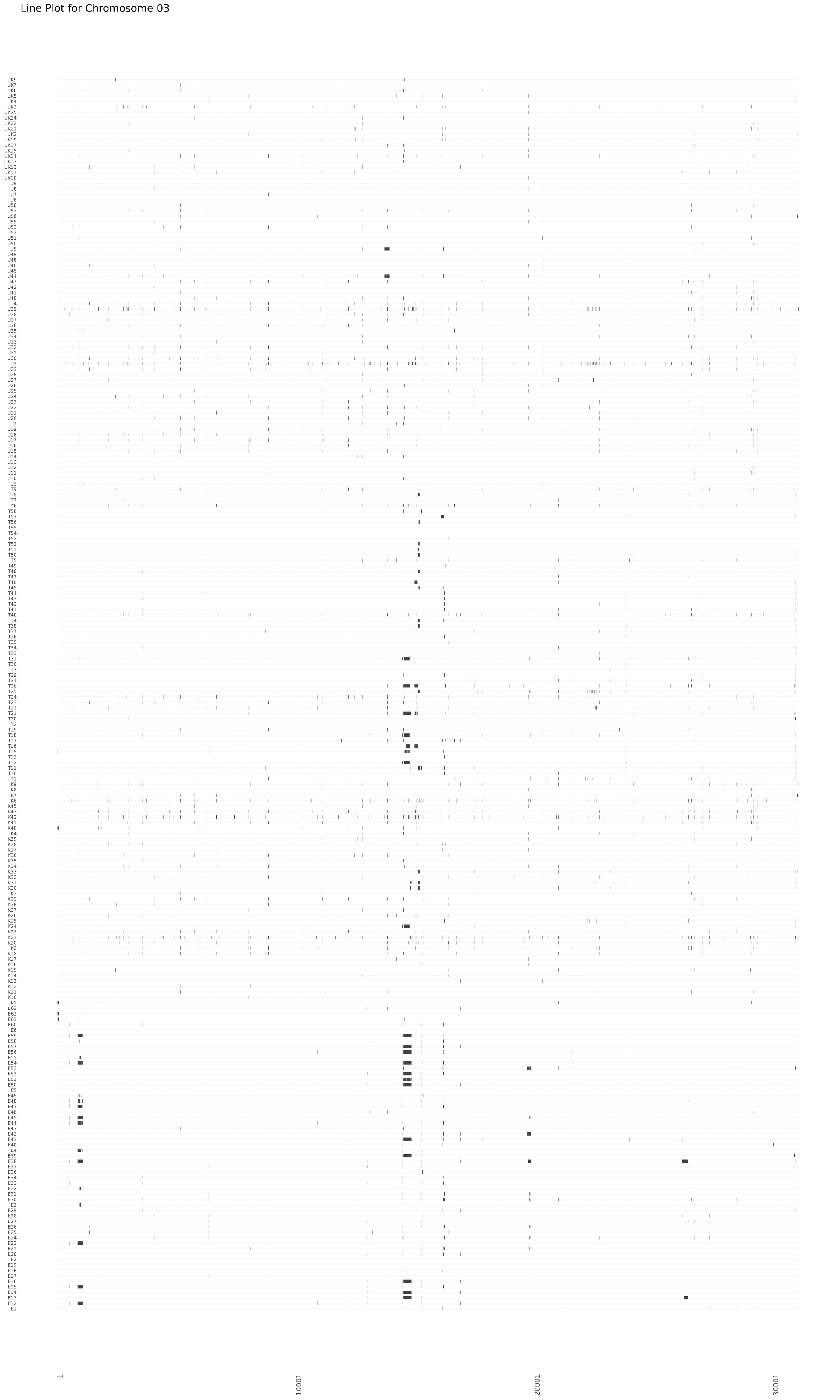

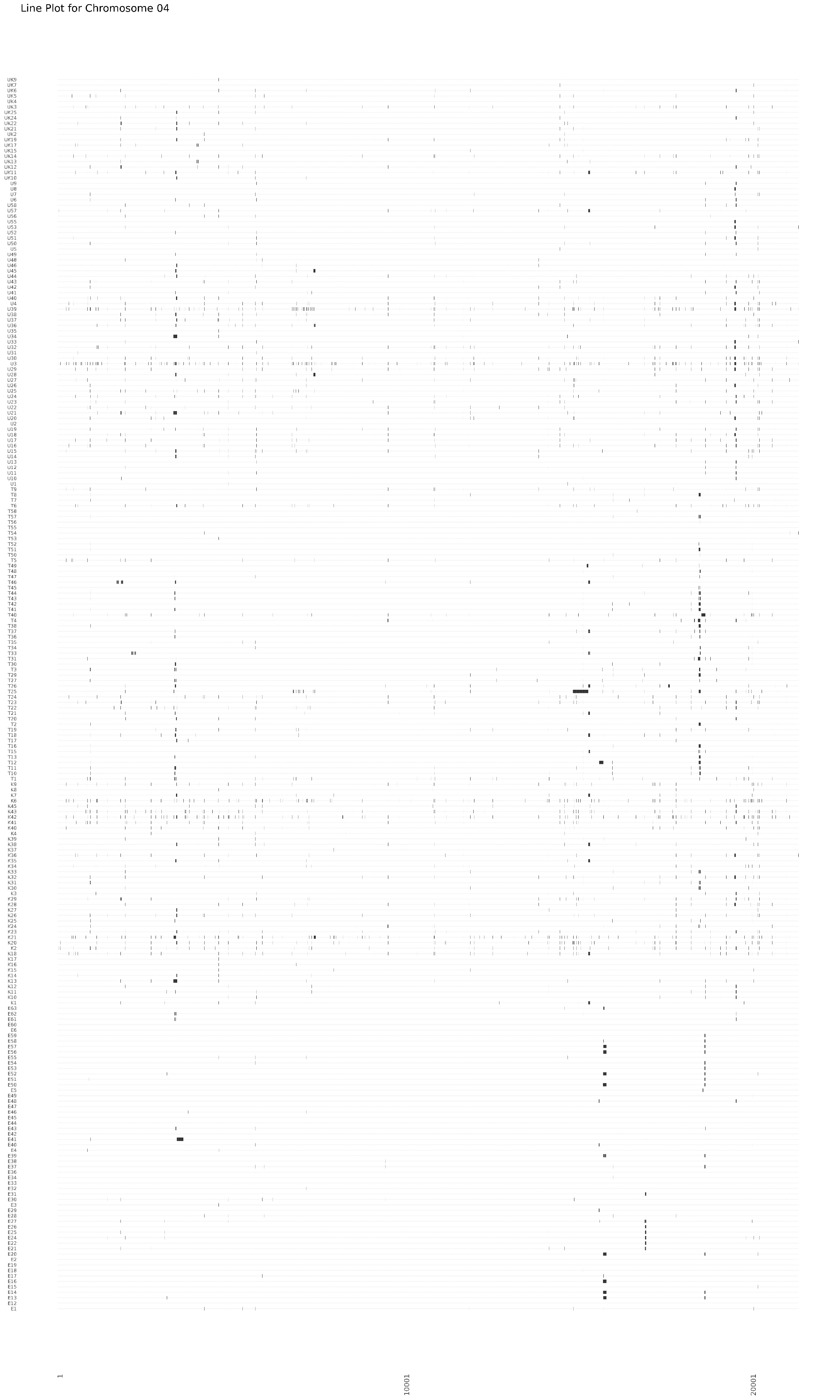

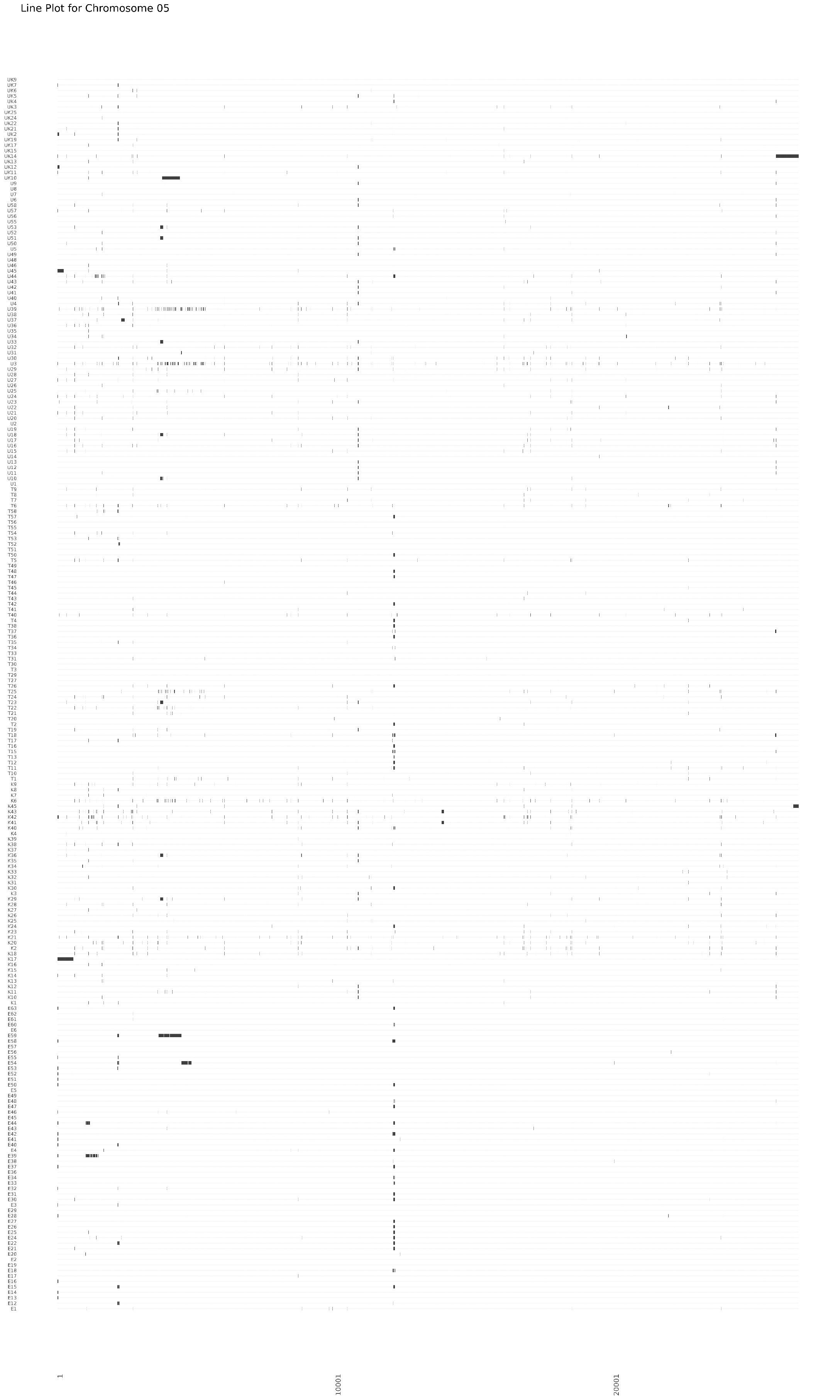

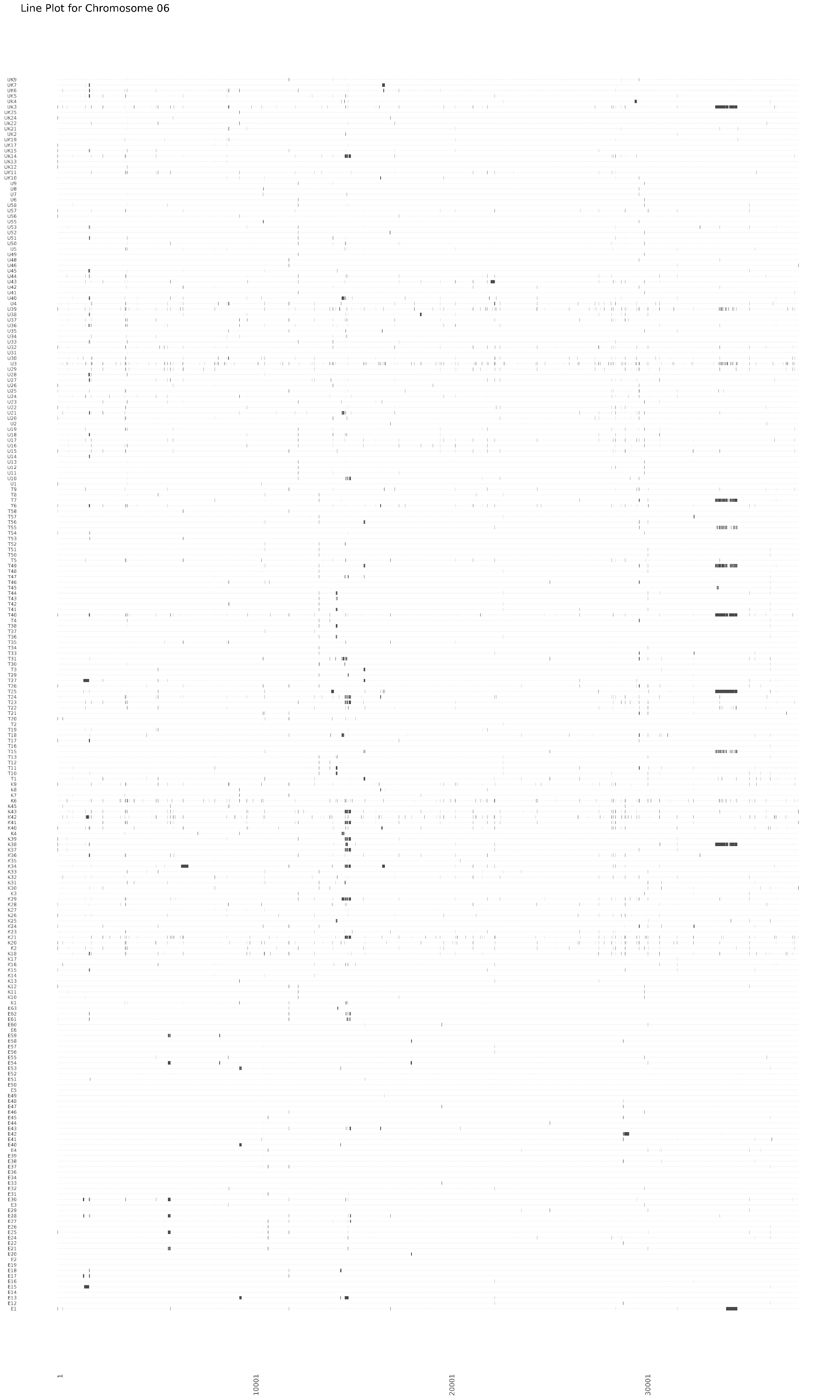

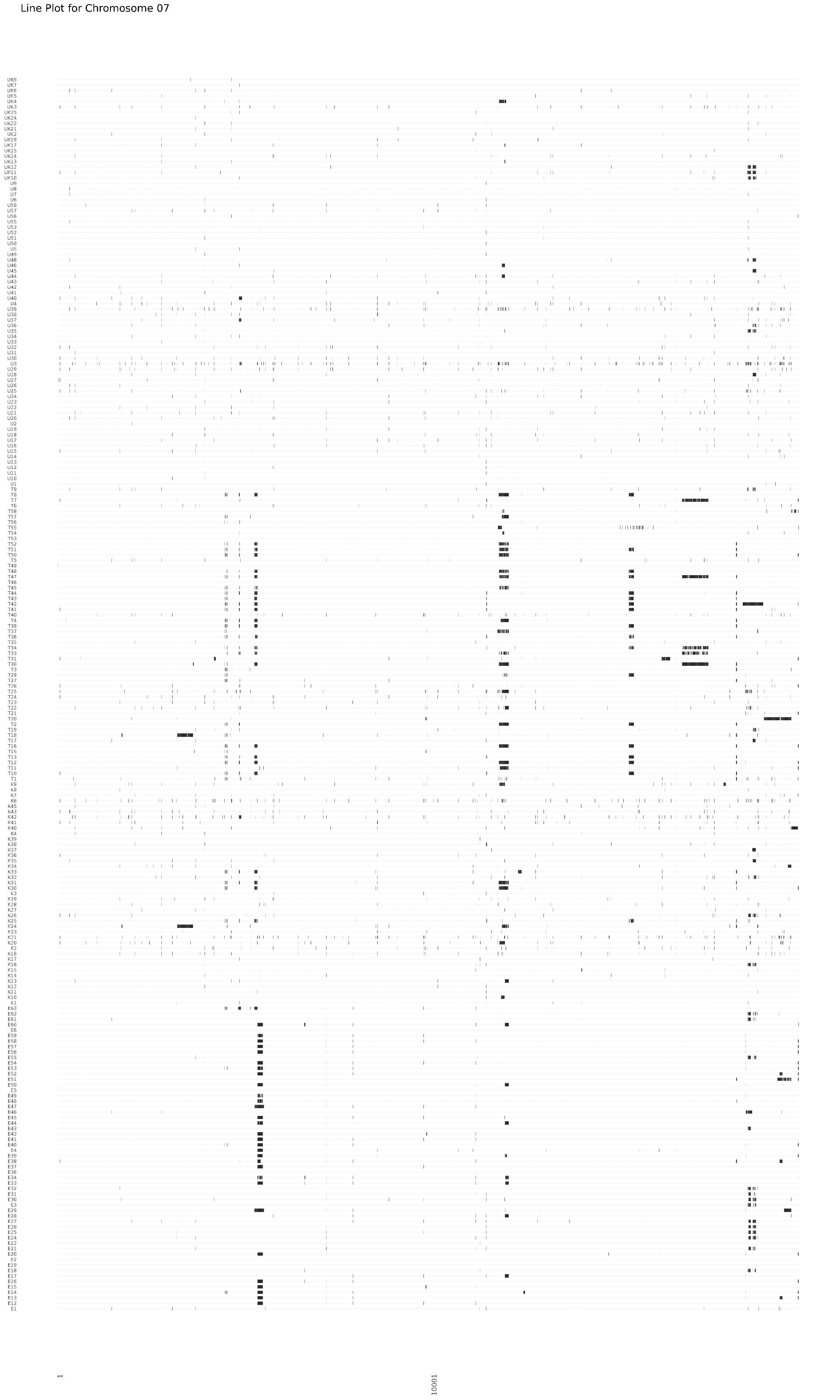

**Figure S5**. Graphical genotypes of 226 finger millet-infecting *M. oryzae* strains (Y-axes) based on 198,454 SNPs (X-axes). Small vertical lines indicate missing data with stretches of missing data likely corresponding to deletions.

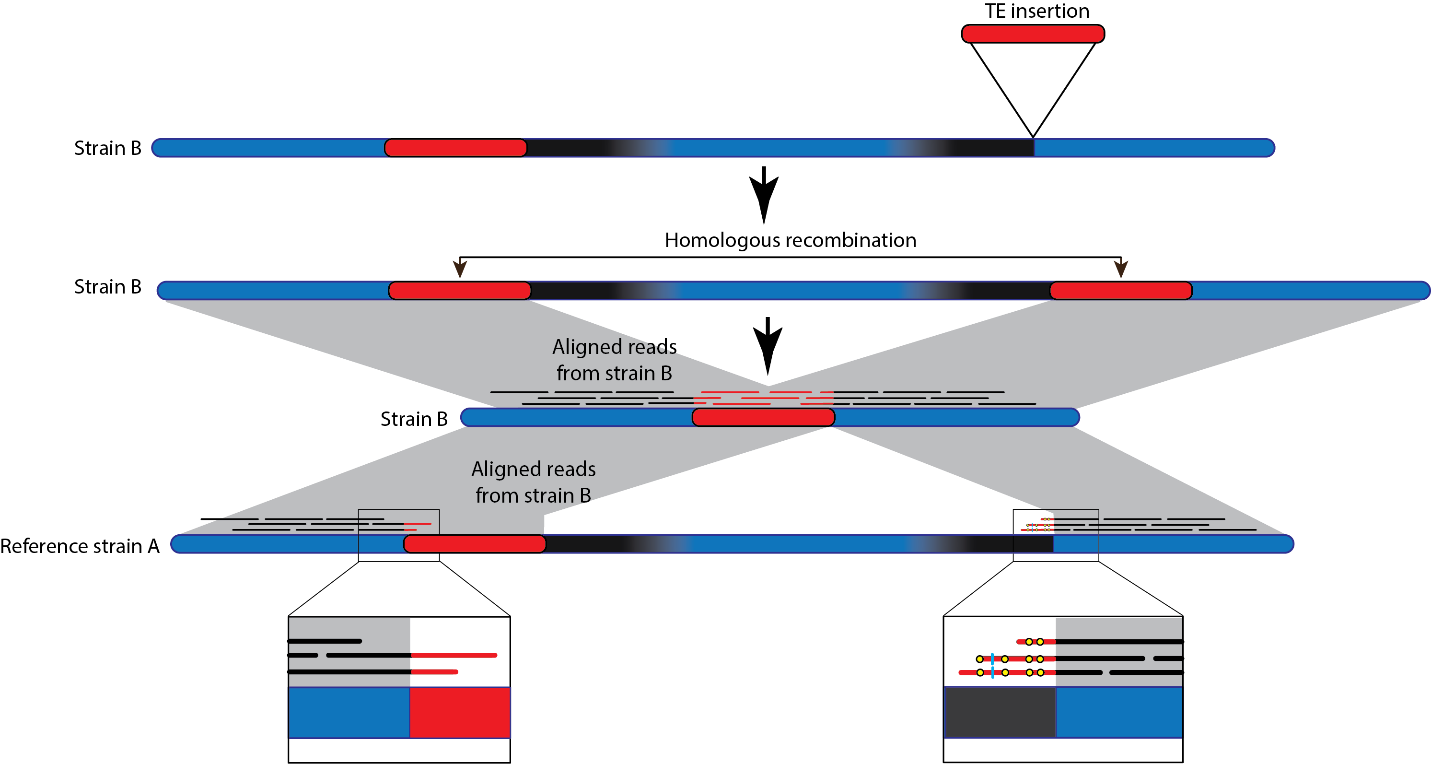

**Figure S6**. Schematic of the mechanism that leads to deletions associated with a transposable element, and how these were identified on a genome-wide scale. A PacBio-based genome assembly is available for Reference strain A. Illumina short-read data is available for strain B. Strain B carries two transposable elements (TEs) from the same family (in red) with at least one of the insertions being absent from the reference strain A (here, one TE is shared with reference strain A). The region flanked by black areas is deleted when homologous recombination takes place between the transposable elements in strain B. Grey regions connect homologous DNA segments. The bottom left inset shows Illumina reads generated in strain B with good alignment over their entire length to the reference assembly which occurs when the reference strain A and resequenced strain B carry the same TE. The bottom right inset shows that reads from strain B extending into a transposon that is differentially present between the resequenced and reference strain will have mismatches (depicted in yellow) and indels (depicted as light blue vertical lines) when aligned to the reference assembly of strain A. If strain B underwent TE-based homologous recombination leading to a deletion, the transition point in the reads will correspond to the deletion breakpoint.

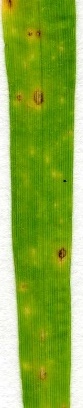

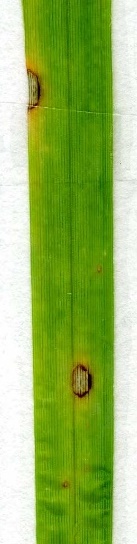

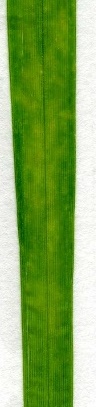

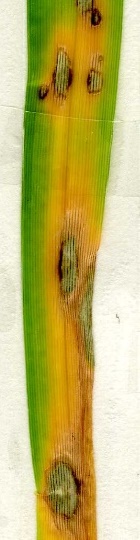

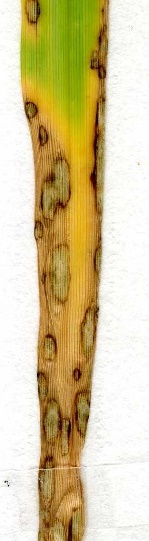

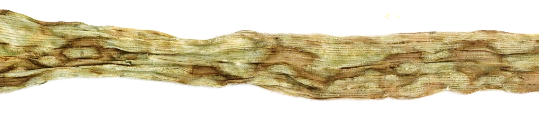

0

1

2

3

4

5

**Figure S7.** Representative examples of infection scores 0-5 following whole-plant spray inoculation assays with *MoE* strain CKF4046 (E2 transformant labeled with enhanced green fluorescent protein (EGFP)) on finger millet cultivars TZA1637 (scores 0 and 1), IE 2555 (score 2), 214988 (score 3), Gulu-E (score 4) and AAUFM-44 (score 5). The six lesion types were defined as follows for finger millet seedlings inoculated under conditions described in Materials and Methods. Type 0, no infection; Type 1, very few small lesions with minimal chlorosis; Type 2, small to medium lesions with few having visibly grey centers, minimal chlorosis; Type 3, several medium to large lesions with some merging, large areas of chlorosis; Type 4, many lesions with several merged lesions, large areas of chlorosis and some collapsing of the leaf; Type 5, complete infection of the exposed leaf, large merged lesions, collapse of leaf and often complete chlorosis of leaf.

AAUFM-44

TZA1637

Gelatin

Percentage of lesion area per leaf (%)

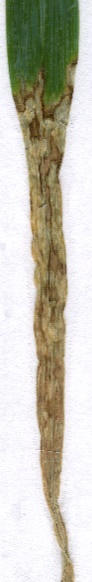

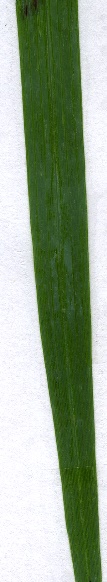

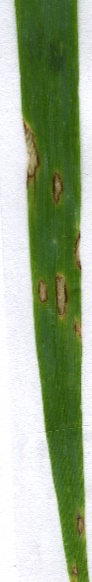

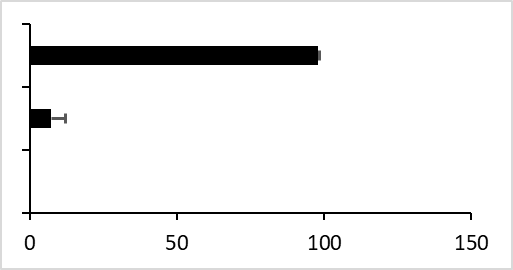

****

A

B

**Figure S8.** Leaf images (A) and bar chart (B) showing level of pathogenicity on susceptible and resistant finger millet cultivars. Inoculation with 0.25% gelatin on AAUFM-44 was used as the negative control. Bar chart shows the percentage of lesion area per marked leaf after infection with fungal spores. Error bar equals standard deviation of the mean (n=3); **** indicates p<0.0001. One-way ANOVA with Tukey's multiple comparisons test was used. Two experiments were performed with similar results.

|  |  |
| --- | --- |
| Gelatin | 0.0607903 |
| TZA1637 | 7.26791 |
| AAUFM-44 | 98.1198 |

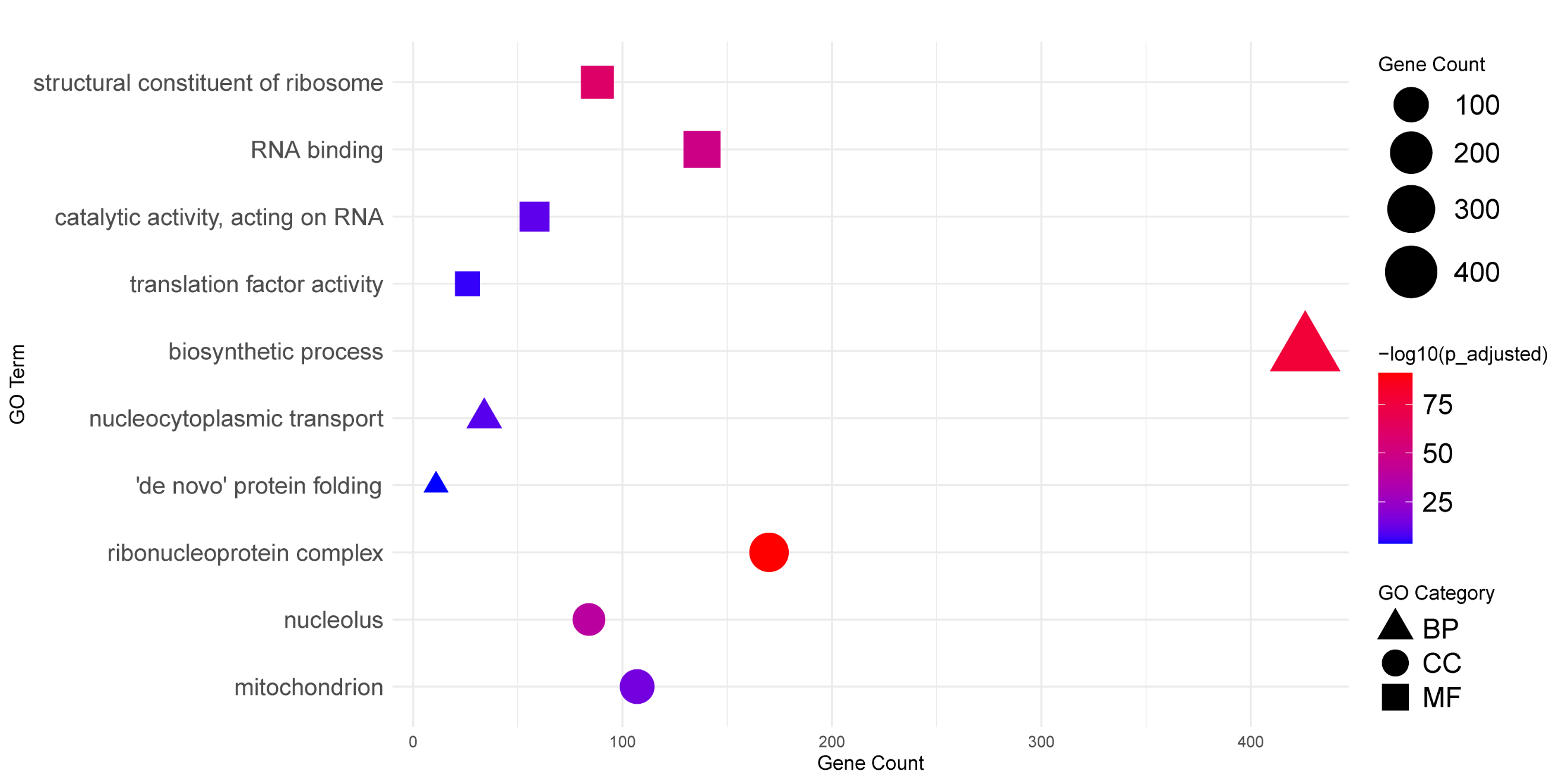

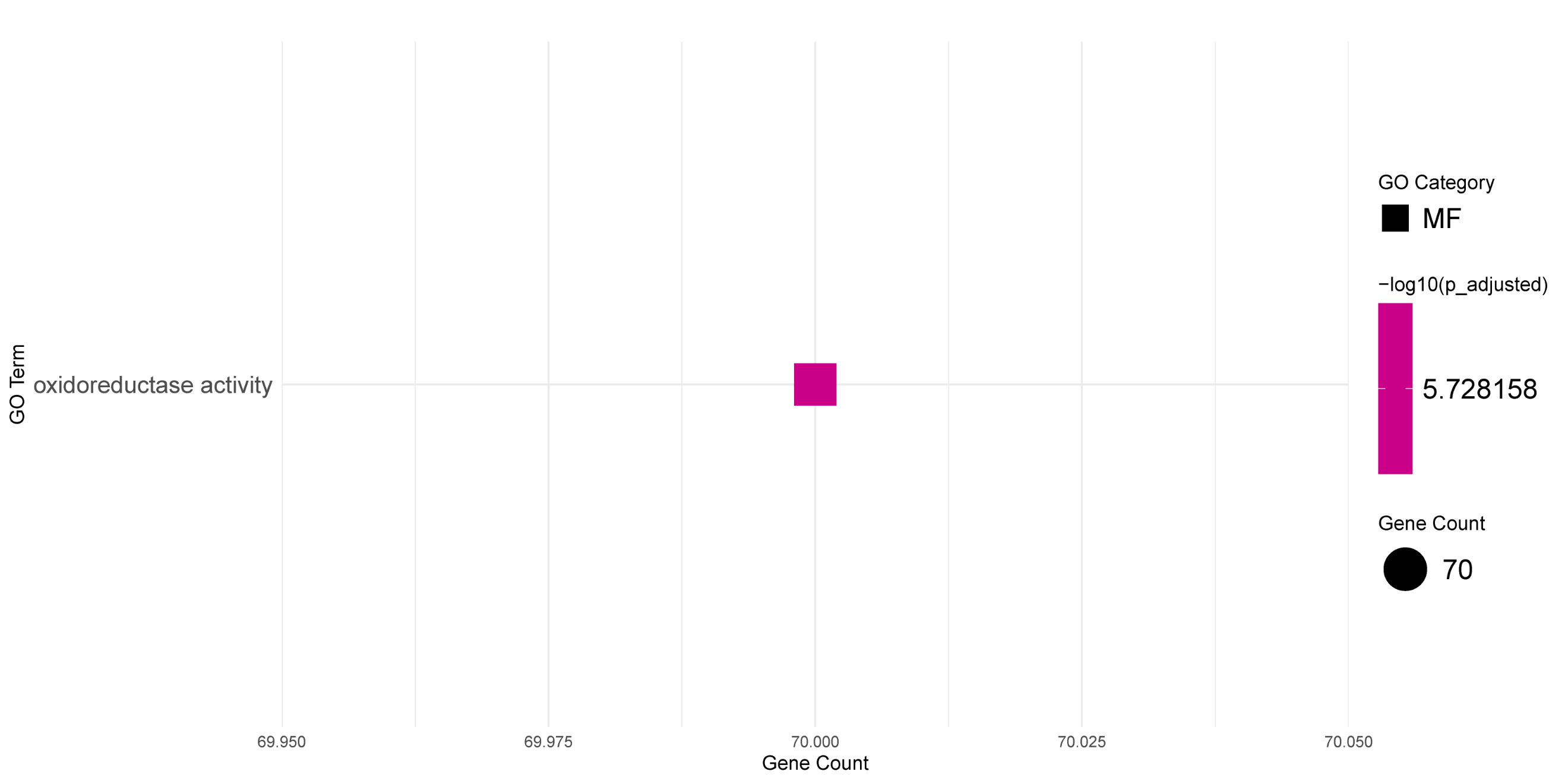

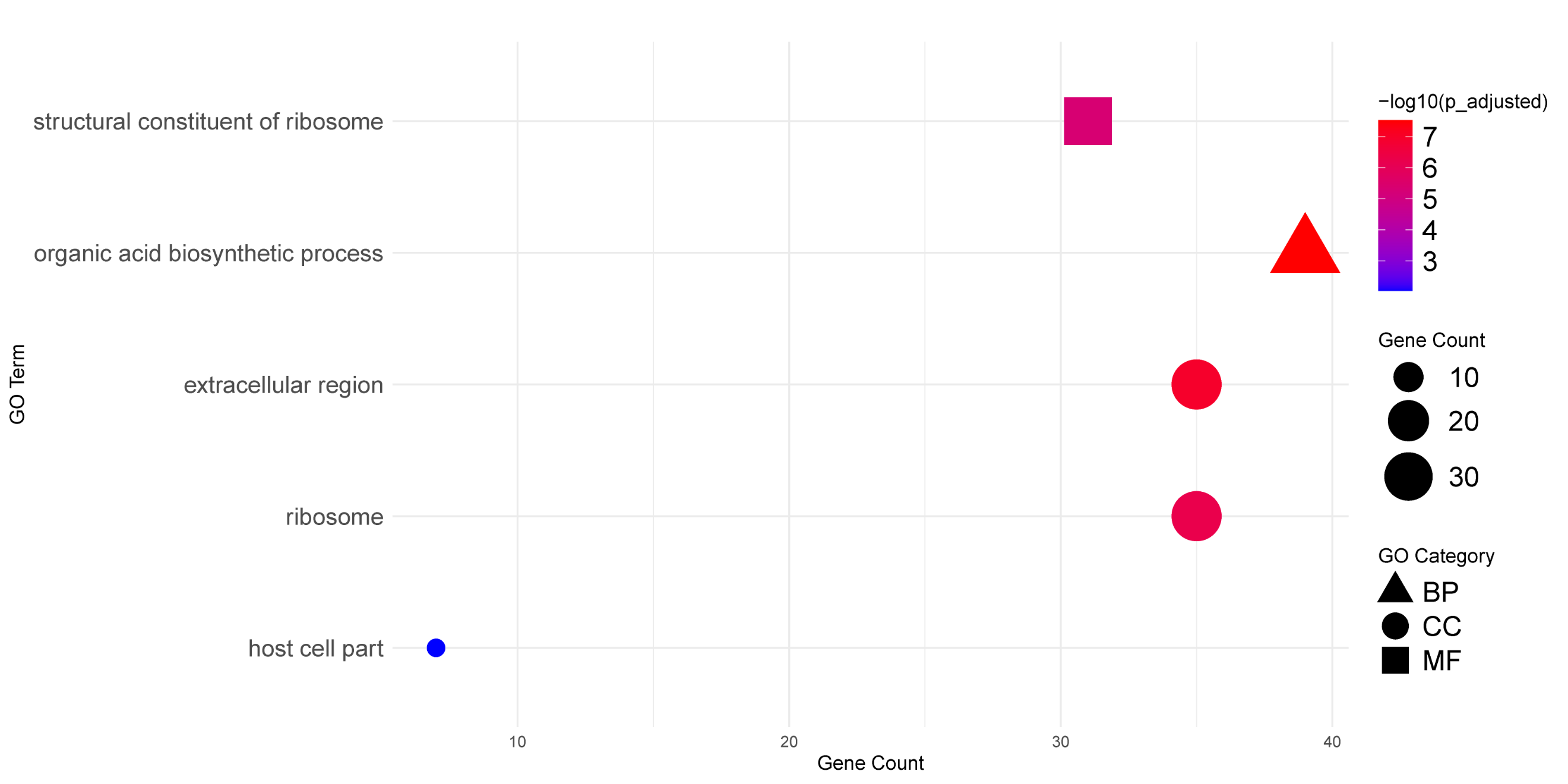

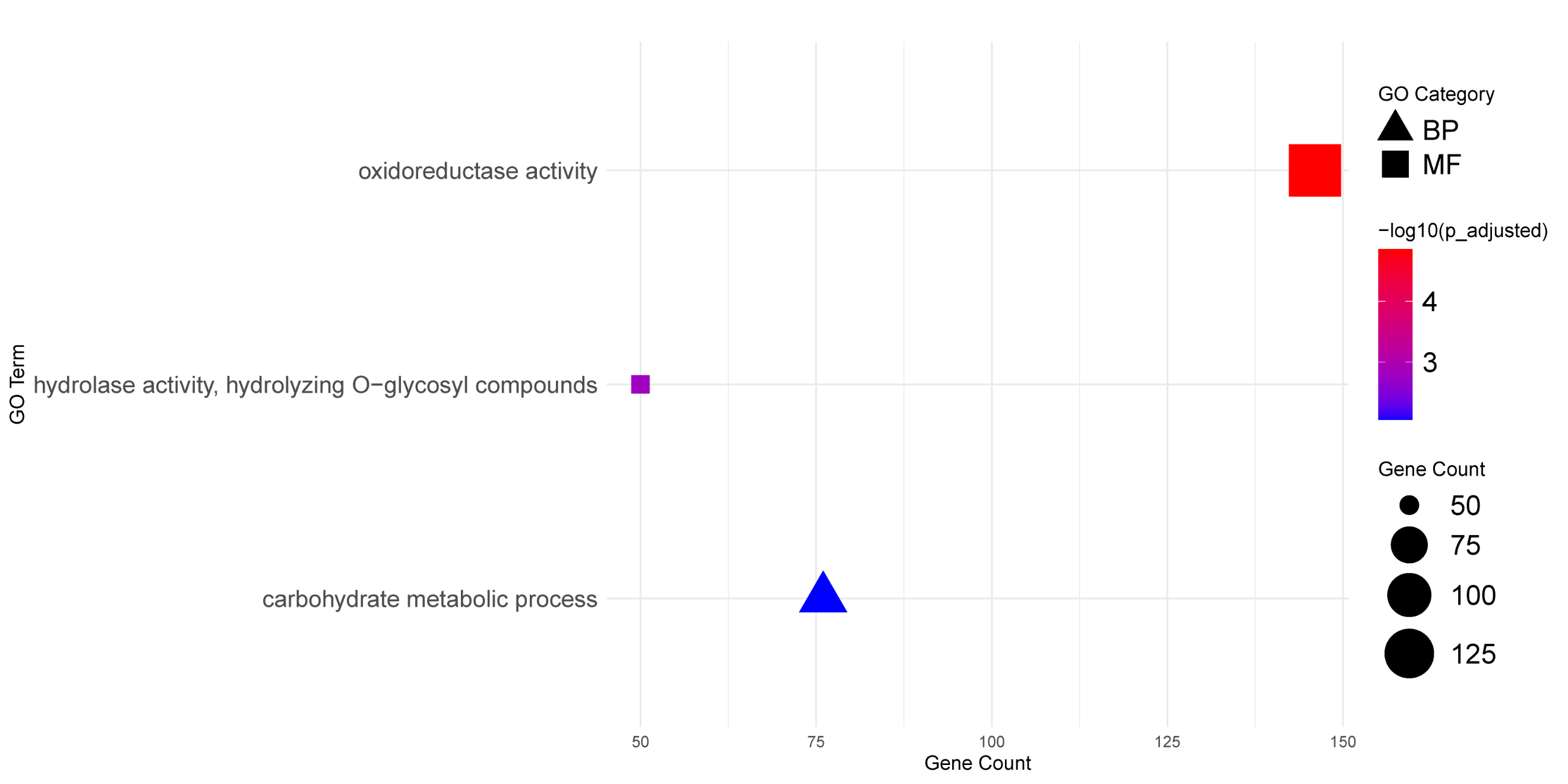

**Figure S9.** Enriched GO driver terms for (A) genes upregulated in the compatible interaction at 30 hp, (B) genes upregulated in the compatible interaction at 48 hpi, (C) genes upregulated in the incompatible interaction at 30 hpi and (D) genes upregulated in the incompatible interaction at 48 hpi.

D

C

A

B

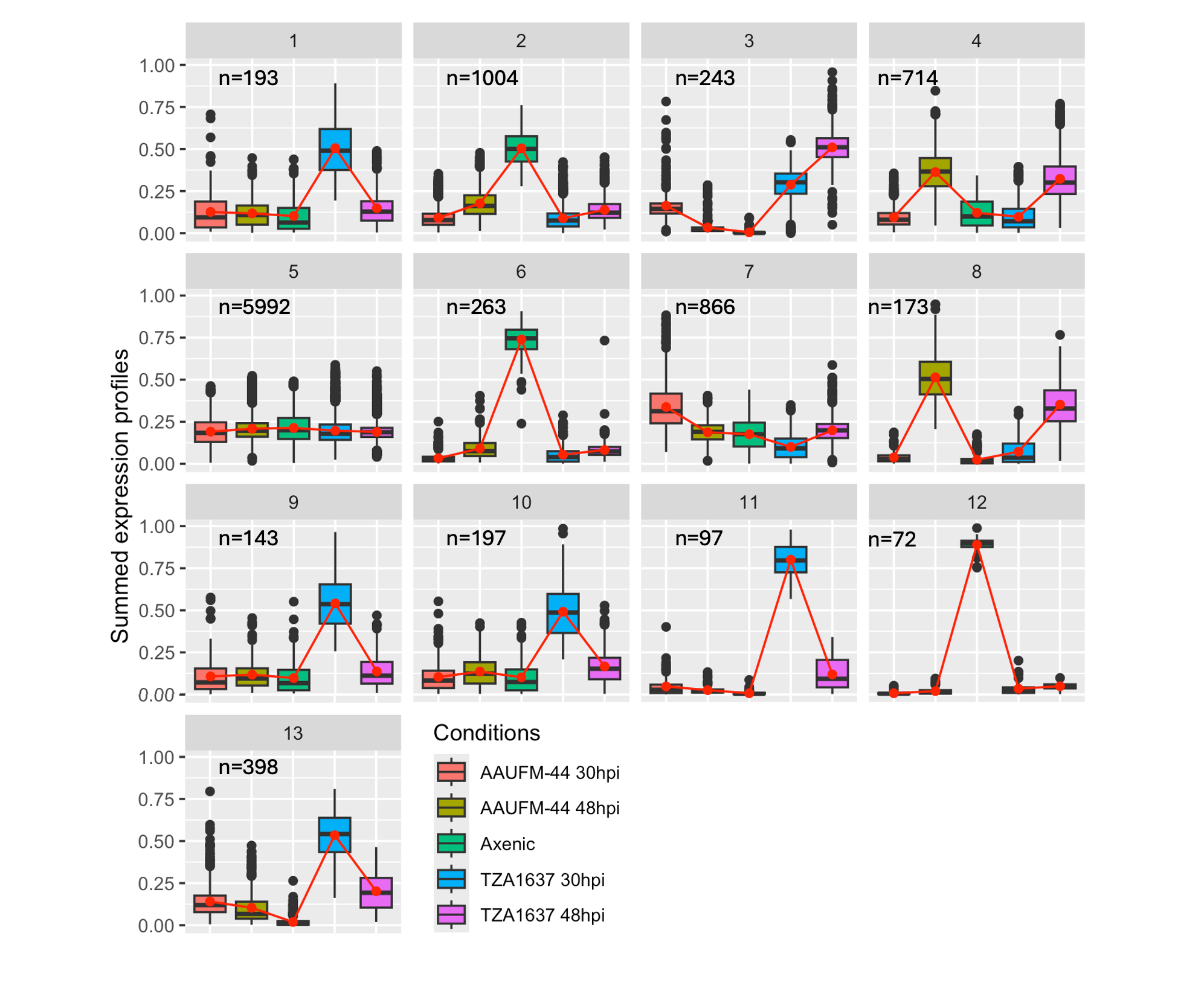

**Figure S10.** Results of co-seq analysis of transcripts expressed in the Ethiopian strain E2 at 30 hpi and 48 hpi in a compatible interaction (AAUFM-44) and an incompatible interaction (TZA1637) as well in axenic culture. A total of 10,355 gene models corresponding to annotated genes were grouped in 13 clusters. The number of genes per cluster is designated by ‘n’.

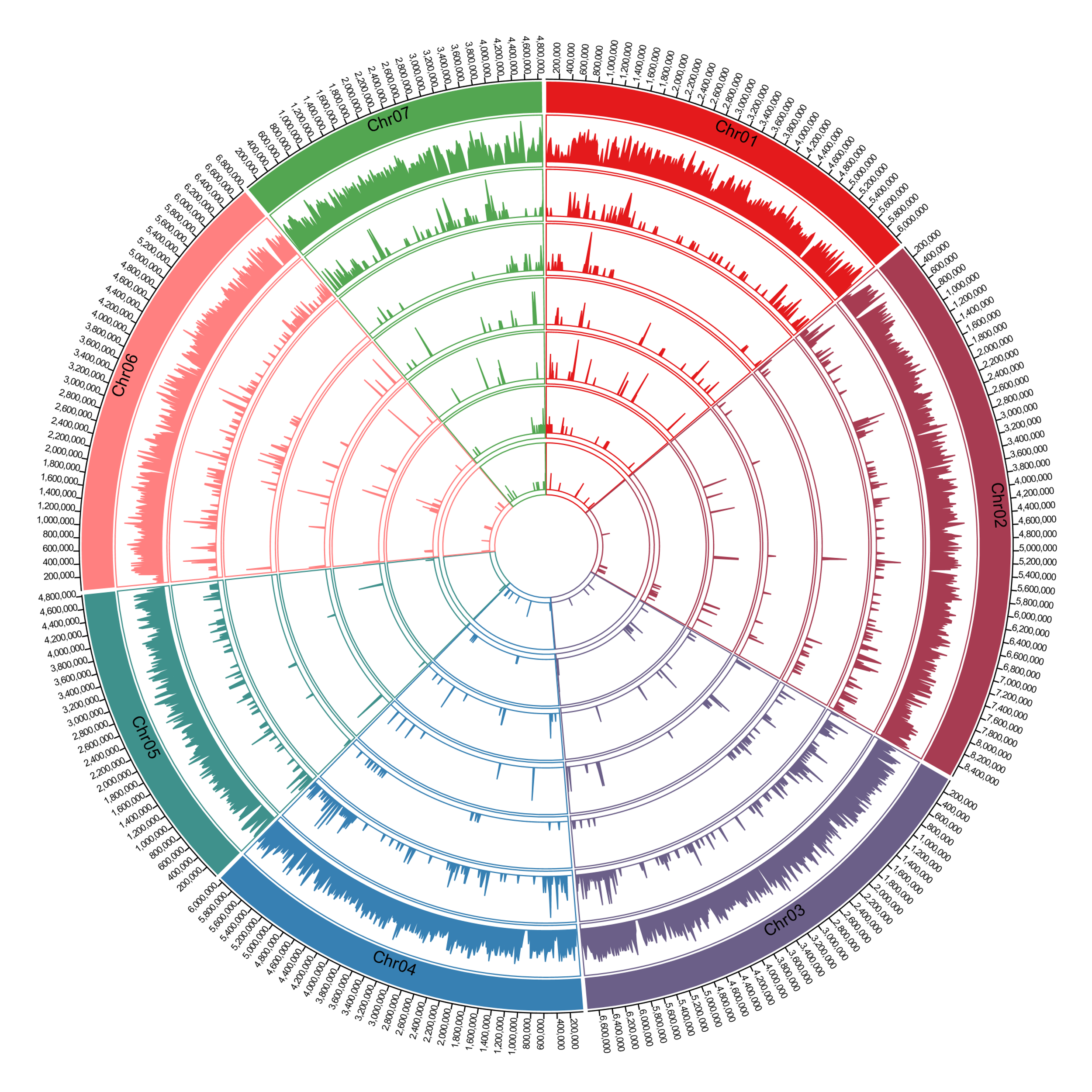

**Figure S11.** Circos diagram showing, from outside to inside, the seven M. oryzae chromosomes (color-coded), and the distribution (number of genes or TEs per 20 kb) across the chromosomes of genes (min = 0, max = 12), effector genes (min = 0, max = 5), and the transposable elements Pot2, Grasshopper, Family 13, Family 1, Inago2 and Pyret (min = 0, max = 5). The distribution was modeled using RepeatMasker v4.1 with default parameters, and includes any repeat fragment larger than 20 bp.

A.

LTR of *Family 1*

TGTTAATTATATGTATTATTTTGCATGTAATAAGCCCACTTTTATTTGTATATAACCCATTTTATATAGTGCCGAAGTGGGGTGGTGGAATTCCGTTAATTTTGTTAATGCAGGAATGCAATTTAAAGGTCAGCGAGTGGGGTTATTTACCCTGTACCGGGTTTAATACCCACCCTGTACCTACGAGTCAGCTACCCTGTACCGGACCGAAAATTTCACCAAGATTTTACGTGCGAACCCAGAGATGGTTTGTTAGTAAATGAGGGTGTTTTTTCAAAACGGCATACAAAATAATTTTTCGGAAAACCATTTGCGGCACCGGAACCTTTTCTGGCGTGCGGACCCCCACTTCGATGTCGGAGAGTATGTAAGGGAAATAAAGCAAAATCTCCCCCACCAATTCAATTATTTATCAGTACCAAAATTATAGTGCAATCCCACCTATCATTTAACGCTTTTAGCACTGGTTATTTACCACGCCGCACCTAGGGTTTACCCCTATATCCCCTGTTAGCACCCGCATTGTTAACA

B.

LTR of *Family 13*

TGTTGGAATAGCCTCACTCGCGCGCGCGCAAGGAAGGGAGTCAATACAGGCCGGGGTTGCGGCTCCGGCGACCGGGTGACTAAGCAGGGAGCACGGACCGCACGAAGACTTTTCACCCGGCGGTAAACGCTGAATTGTAGTCGCACGACTTAGATAGAAGACAGACCGAATATATCCAATTGACAGAGGTATAACGTTCTCACCATACGAATAGATCTACCACGGATGCATCACGTAGATCCACGAATCTACGAACA

**Figure S12.** LTR sequences of LTR-retrotransposons with ID *Family 1* and *Family 13* identified in *MoE* reference strain E2. The 25 bp shown in red were used to search for presence of the elements in Illumina whole-genome shotgun reads generated for blast isolates.

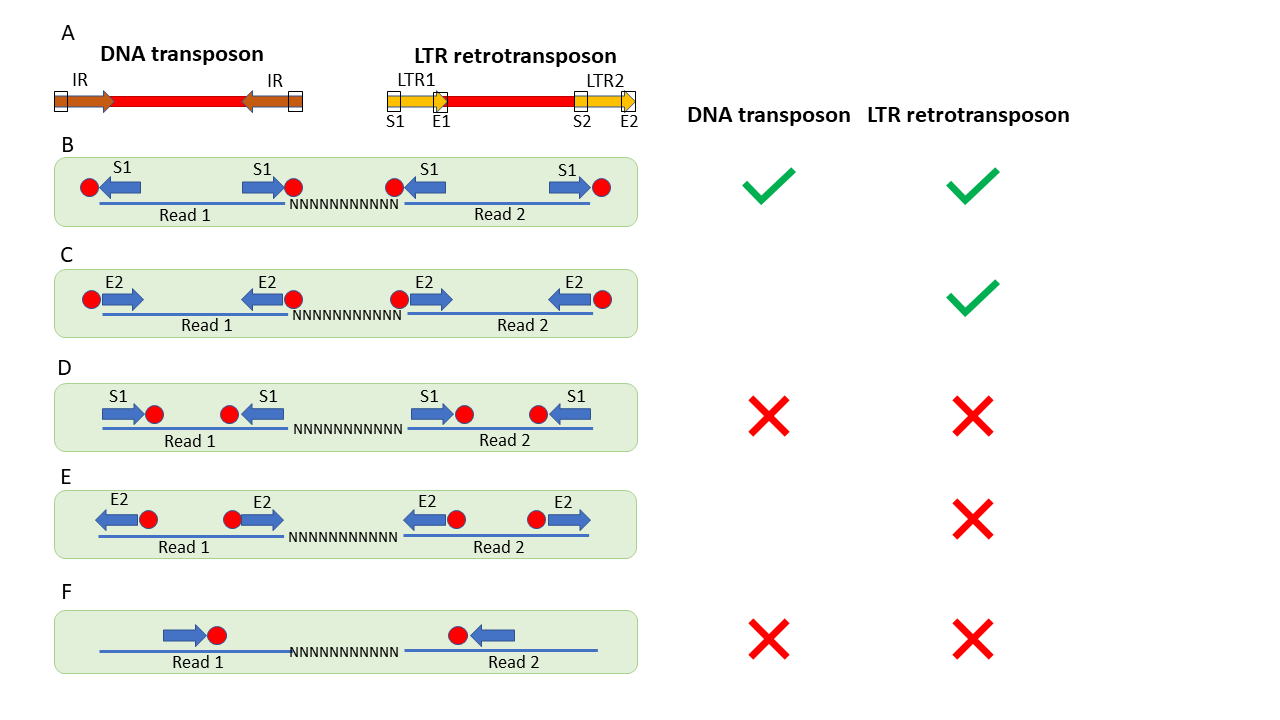
**Figure S13.** General structure of DNA and LTR-retrotransposons (A), and schematics of Illumina reads with homology to the terminal 25 bp of the TEs (B-F). A. Inverted repeats (IRs) in the DNA transposon are shown in brown; Long terminal repeats (LTRs) in the LTR-retrotransposon are shown in yellow. The black boxes on the IRs and LTRs indicate the regions used in the BLASTN searches. B-F. Blue lines separated by Ns represent paired-end Illumina reads. Blue arrows correspond to boxed regions in A, and indicate regions and direction of homology. The red dot represents the position of the internal region of a TE. Illumina reads in which the terminal 20 – 25 bp of the IR or LTR mapped to the end of a read were retained as indicated with a green tick symbol (B, C). Illumina reads with BLASTN hits were discarded if more than 25 bp of the repeat was present on the read (D-F) indicated by the terminal 25 bp of the TE mapping to the end of the read but the TE extending further into the read (D, E), or the terminal 25 bp of the TE not mapping to the end of the read (F). Because we were unable to differentiate E1 from E2, and S1 from S2 (see Figure A), some Illumina reads that consisted completely of TE sequence were also retained at this stage of the analysis.

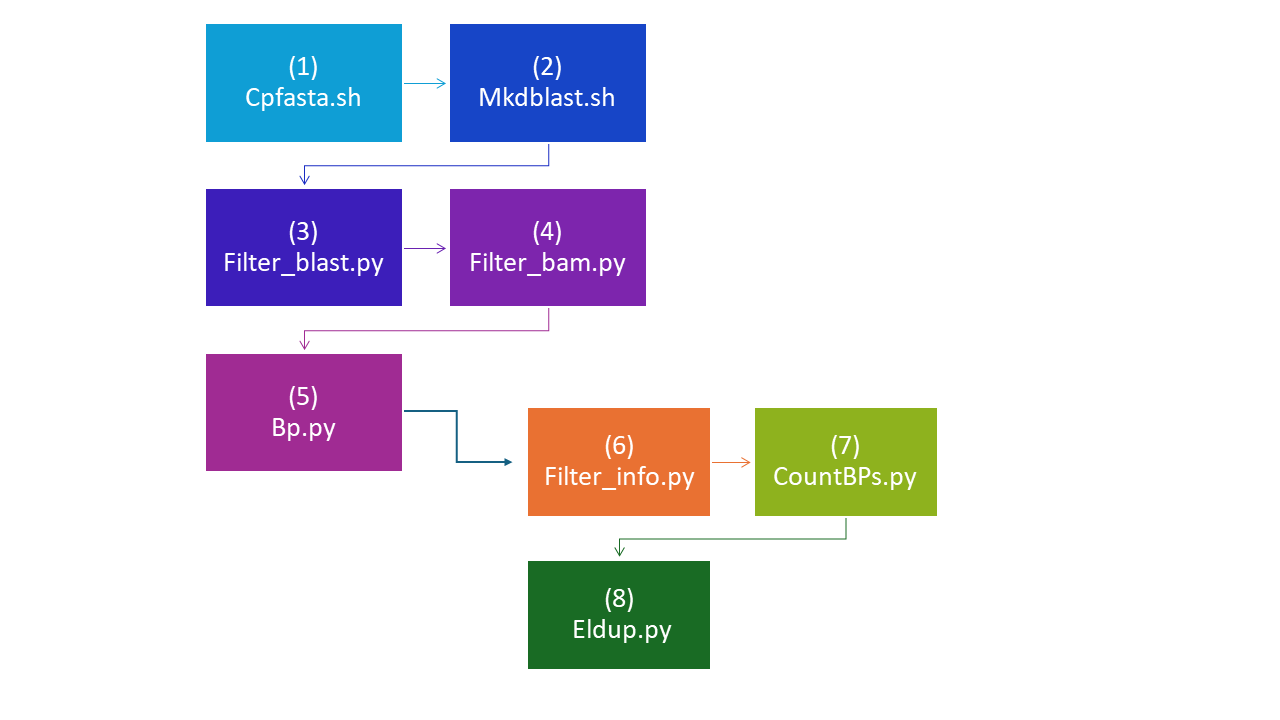

**Figure S14.** Flow chart showing scripts used in identification of deletions associated with a transposable element. Scripts are available from Github (https://github.com/ysahin1/DevosLab-Pan_genome_fingermillet_infecting_MagnaportheOryzae).

**Supplementary Tables**

**Table S1**. Summary of country, tissue origin and year of collection of sequenced finger millet-infecting *M. oryzae* strains

|  | **Leaf^1^** | | **Peduncle^1^** | | **Panicle^1^** | |
| --- | --- | --- | --- | --- | --- | --- |
| **Country** | **2000-2002** | **2015-2017** | **2000-2002** | **2015-2017** | **2000-2002** | **2015-2017** |
| Ethiopia | 0 | 0 | 0 | 33 | 0 | 23 |
| Kenya | 0 | 0 | 7^2^ | 37 | 3 | 4 |
| Tanzania | 0 | 0 | 0 | 47 | 0 | 7 |
| Uganda | 3 | 0 | 1 | 53 | 5 | 3 |

^1^ Organs from which the strains were isolated; Note, however, that strains can infect all organs

^2^ For two of the isolates, the source material is either peduncle or panicle

**Table S2.** PacBio library statistics for the libraries included in the *M. oryzae* strain E2 genome assembly and their respective assembled sequence coverage levels

| **Cutoff** | **Number of Reads** | **Basepairs** | **Average Read Length** | **Coverage** |
| --- | --- | --- | --- | --- |
| 0 | 1,020,525 | 10,596,160,418 | 7,855 | 235.47x |
| 1,000 | 944,437 | 10,559,911,318 | 8,669 | 234.66x |
| 2,000 | 871,207 | 10,449,642,497 | 9,498 | 232.21x |
| 3,000 | 797,851 | 10,266,524,021 | 10,392 | 228.14x |
| 4,000 | 728,913 | 10,025,659,849 | 11,293 | 222.79x |
| 5,000 | 665,522 | 9,740,893,319 | 12,203 | 216.46x |
| 6,000 | 606,834 | 9,418,653,570 | 13,110 | 209.30x |
| 7,000 | 552,864 | 9,068,232,061 | 14,015 | 201.52x |
| 8,000 | 503,282 | 8,696,716,072 | 14,917 | 193.26x |
| 9,000 | 457,217 | 8,305,471,885 | 15,836 | 184.57x |
| 10,000 | 414,792 | 7,902,790,040 | 16,752 | 175.62x |
| 11,000 | 375,480 | 7,490,238,158 | 17,684 | 166.45x |
| 12,000 | 339,574 | 7,077,665,886 | 18,631 | 157.28x |
| 13,000 | 306,841 | 6,668,830,813 | 19,575 | 148.20x |
| 14,000 | 276,905 | 6,264,881,035 | 20,535 | 139.22x |
| 15,000 | 249,496 | 5,867,698,949 | 21,491 | 130.39x |
| 16,000 | 224,621 | 5,482,335,798 | 22,453 | 121.83x |
| 17,000 | 202,029 | 5,109,785,991 | 23,398 | 113.55x |
| 18,000 | 181,493 | 4,750,604,043 | 24,321 | 105.57x |
| 19,000 | 163,311 | 4,414,390,437 | 25,190 | 98.10x |

**Table S3.** Summary statistics of the initial output of the QUIVER polished MECAT assembly

| **Minimum Scaffold Length^1^** | **Number of Scaffolds** | **Number of Contigs** | **Scaffold Size** | **Basepairs** | **% Non-gap Basepairs** |
| --- | --- | --- | --- | --- | --- |
| 5 Mb | 3 | 3 | 28,497,616 | 28,497,616 | 100.00% |
| 2.5 Mb | 6 | 6 | 37,528,315 | 37,528,315 | 100.00% |
| 1 Mb | 8 | 8 | 40,699,759 | 40,699,759 | 100.00% |
| 500 Kb | 12 | 12 | 43,322,518 | 43,322,518 | 100.00% |
| 250 Kb | 13 | 13 | 43,744,214 | 43,744,214 | 100.00% |
| 100 Kb | 14 | 14 | 43,924,383 | 43,924,383 | 100.00% |
| 50 Kb | 16 | 16 | 44,053,457 | 44,053,457 | 100.00% |
| 25 Kb | 21 | 21 | 44,253,274 | 44,253,274 | 100.00% |
| 10 Kb | 27 | 27 | 44,329,515 | 44,329,515 | 100.00% |
| 5 Kb | 29 | 29 | 44,347,618 | 44,347,618 | 100.00% |
| 2.5 Kb | 29 | 29 | 44,347,618 | 44,347,618 | 100.00% |
| 1 Kb | 29 | 29 | 44,347,618 | 44,347,618 | 100.00% |
| 0 bp | 29 | 29 | 44,347,618 | 44,347,618 | 100.00% |

^1^ For each set of scaffolds greater than the size listed, the total contigs and total assembled basepairs are shown

|  | E2 | | | Mz5-1-6 | | | B71 | | | P131 | | |
| --- | --- | --- | --- | --- | --- | --- | --- | --- | --- | --- | --- | --- |
|  | Number of elements | Length occupied (bp) | Percentage of sequence (%) | Number of elements | Length occupied (bp) | Percentage of sequence (%) | Number of elements | Length occupied (bp) | Percentage of sequence (%) | Number of elements | Length occupied (bp) | Percentage of sequence (%) |
| **Retroelements** | **3427** | **3854360** | **8.67** | **2978** | **3487919** | **8.17** | **3681** | **3677200** | **8.20** | **4132** | **4784421** | **11.06** |
| LINEs | 282 | 321293 | 0.72 | 173 | 144826 | 0.34 | 479 | 642855 | 1.43 | 444 | 823152 | 1.90 |
| CRE/SLACS | 93 | 44885 | 0.10 | 53 | 20124 | 0.05 | 197 | 296324 | 0.66 | 10 | 1800 | 0.00 |
| LTR elements | 3145 | 3533067 | 7.95 | 2805 | 3343093 | 7.83 | 3202 | 3034345 | 6.77 | 3688 | 3961269 | 9.15 |
| Ty1/Copia | 491 | 542604 | 1.22 | 485 | 513544 | 1.20 | 542 | 431074 | 0.96 | 467 | 598761 | 1.38 |
| Gypsy/DIRS1 | 1031 | 948017 | 2.13 | 819 | 911397 | 2.13 | 978 | 777790 | 1.73 | 1384 | 917564 | 2.12 |
| **DNA transposons** | **1205** | **560751** | **1.26** | **1041** | **561563** | **1.32** | **1238** | **494134** | **1.10** | **1119** | **836475** | **1.93** |
| Tc1-IS630-Pogo | **5** | **5440** | **0.01** | **3** | **3090** | **0.01** |  |  |  | **4** | **5524** | **0.01** |
| **Unclassified** | **185** | **29007** | **0.07** | **175** | **29641** | **0.07** | **194** | **30479** | **0.07** | **141** | **23331** | **0.05** |
| **Total interspersed repeats** |  | **4444118** | **10.00** |  | **4079123** | **9.55** |  | **4201813** | **9.37** |  | **5644227** | **13.04** |
| **Simple repeats** | **13665** | **506625** | **1.14** | **13529** | **512240** | **1.20** | **13297** | **489465** | **1.09** | **12074** | **447142** | **1.03** |
| **Low complexity** | **1759** | **78020** | **0.18** | **1714** | **76800** | **0.18** | **1709** | **76742** | **0.17** | **1529** | **67395** | **0.16** |
| **Total repeats** |  | **5028763** | **11.31** |  | **4668163** | **10.93** |  | **4768020** | **10.63** |  | **6158764** | **14.23** |

**Table S4.** Summary of repeat annotation of blast strains E2, Mz5-1-6, B71 and P131

**Table S5.** Haplotypes comprising ≥ 2 strains with a similarity higher than 99.9%

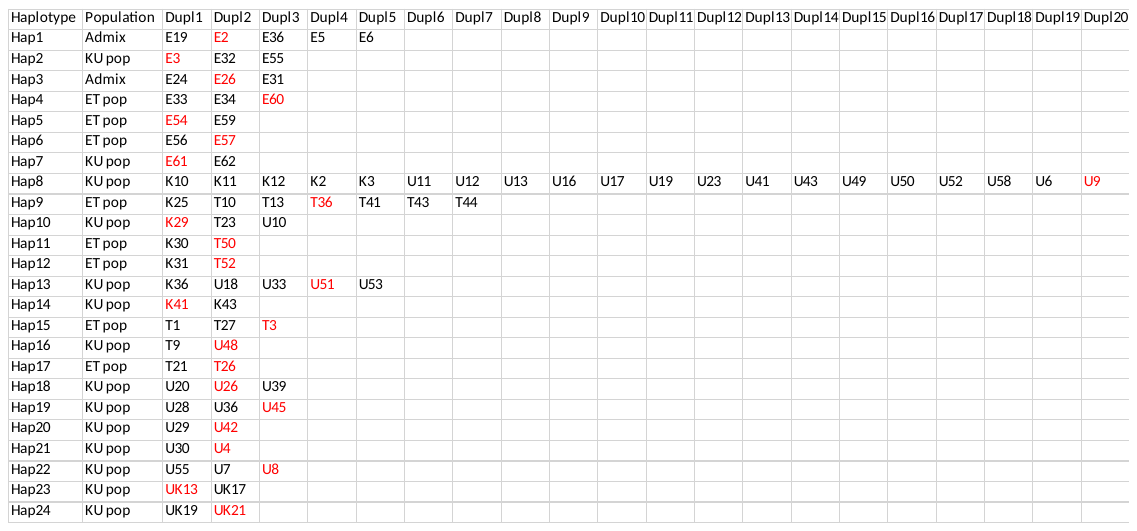

Representative strains for each haplotype are indicated in red

**Table S6. Similarity between strains isolated from peduncle and panicle from the same plant**

| mci ID | Population^1^ | Organ from which mci was isolated | % similarity |
| --- | --- | --- | --- |
| E4 | 0.877 | Panicle | 87.9 |
| E5 | 0.874 | Peduncle |  |
| E61 | KU | Panicle | 100 |
| E62 | KU | Peduncle |  |
| E38 | ET | Panicle | 90.2 |
| E39 | ET | Peduncle |  |
| E47 | ET | Panicle | 90.5 |
| E48 | ET | Peduncle |  |
| E49 | ET | Panicle | 89.4 |
| E50 | ET | Peduncle |  |
| E41 | ET | Panicle | 89.3 |
| E42 | ET | Peduncle |  |
| E43 | KU | Panicle | 45.2 |
| E44 | ET | Peduncle |  |
| E51 | ET | Panicle | 93.1 |
| E52 | ET | Peduncle |  |
| E53 | ET | Panicle | 88.5 |
| E54 | ET | Peduncle |  |
| E13 | ET | Panicle | 92.9 |
| E14 | ET | Peduncle |  |
| E56 | ET | Panicle | 100 |
| E57 | ET | Peduncle |  |

^1^ ET: ≥90% membership to Ethiopia/Tanzania subpopulation; KU: ≥90% membership Kenya/Uganda subpopulation; Values >50% have majority membership to ET population

**Table S7**. *M. oryzae* strains with different host specificities used in phylogenetic analyses

| **Isolate ID** | **Host** | **Source Country** | **SRA Accession No** |
| --- | --- | --- | --- |
| BtP29^18^ | *Bromus tectorum* | Paraguay | SAMN05898532 |
| DsU167^19^ | *Digitaria sanguinalis* | Uruguay | SRR14705972 |
| EcU170^19^ | *Echinochloa sp.* | Uruguay | SRR14705961 |
| EiU169-v1^20^ | *Eleusine* | Uruguay | SRR14705979 |
| EcJP29^21^ | *Eleusine coracana* | Japan | SRR14705950 |
| EcMG03^22^ | *Eleusine coracana* | India: Bangalore | SRR3056584 |
| EcMZ5-1-6^23^ | *Eleusine coracana* | Japan | SRR8258940 |
| EcMG04^22^ | *Eleusine coracana* | India: Bangalore | SRR3056585 |
| EcMG12^22^ | *Eleusine coracana* | India: Bangalore | SRR3056614 |
| EiU229^20^ | *Eleusine indica* | Uruguay | SRR14705991 |
| EiJA56^19^ | *Eleusine indica* | Brazil | SRR14705992 |
| Ei9604^24^ | *Eleusine indica* | China: Zhejiang | SRR11836442 |
| Ei9411^24^ | *Eleusine indica* | China: Fujian | SRR11836441 |
| Ei8927^25^ | *Eleusine indica* | Philippines | SRR14705983 |
| Ei88365^25^ | *Eleusine indica* | Philippines | SRR14705984 |
| Ei8303^25^ | *Eleusine indica* | Philippines | SRR14705985 |
| EiU231^20^ | *Eleusine indica* | Uruguay | SRR14705980 |
| EiJA178^19^ | *Eleusine indica* | Brazil | SRR14705981 |
| EcuAR4^26^ | *Eragrostis curvula* | Japan | SRR14705978 |
| FaPg1213^18^ | *Festuca arundinaceum* | Georgia, USA | SAMN08009569 |
| Lh8844^25^ | *Leersia hexandra* | Philippines | SRR14705976 |
| LpHO^19^ | *Lolium perenne* | Richmond, PA, USA | SAMN08009558 |
| LpCHW^18^ | *Lolium perenne* | Annapolis, MD, USA | SAMN08009549 |
| OsU198^19^ | *Oryza sativa* | Uruguay | SRR14705967 |
| OsU107^19^ | *Oryza sativa* | Uruguay | SRR14705968 |
| OsSSID116^20^ | *Oryza sativa* | USA | SRR14705969 |
| Pr88165^20^ | *Panicum repens* | Philippines | SRR14705966 |
| PnML36^27^ | *Pennisetum sp.* | Mali | SRR14705965 |
| PtKY18-1^19^ | *Poa trivialis* | USA | SRR14705964 |
| SiU232^20^ | *Setaria italica* | Uruguay | SRR14705963 |
| SiMG03^22^ | *Setaria italica* | India: Bangalore | SRR3056611 |
| Sv9623^24^ | *Setaria viridis* | China: Zhejiang | SRR11836445 |
| Sv9610^24^ | *Setaria viridis* | China: Zhejiang | SRR11836446 |
| Ta37^28^ | *Triticum aestivum* | Brazil | SRR3624699 |
| Ta117^28^ | *Triticum aestivum* | Brazil | SRR3624701 |
| Ta127^28^ | *Triticum aestivum* | Brazil | SRR3624702 |
| Ta169^28^ | *Triticum aestivum* | Brazil | SRR3624703 |
| Ta204^28^ | *Triticum aestivum* | Brazil | SRR3624705 |
| Ta205^28^ | *Triticum aestivum* | Brazil | SRR3624706 |
| Ta032i^28^ | *Triticum aestivum* | Brazil | SRR3624698 |
| Ta053i^28^ | *Triticum aestivum* | Brazil | SRR3624700 |
| TaWB127^29^ | *Triticum aestivum* | Brazil | SRR14705989 |
| TaT46-2^19^ | *Triticum aestivum* | Brazil | SRR14705951 |
| E2 | *Eleusine coracana* | Ethiopia |  |
| E28 | *Eleusine coracana* | Ethiopia |  |
| E48 | *Eleusine coracana* | Ethiopia |  |
| E61 | *Eleusine coracana* | Ethiopia |  |
| K27 | *Eleusine coracana* | Kenya |  |
| T35 | *Eleusine coracana* | Tanzania |  |
| T38 | *Eleusine coracana* | Tanzania |  |
| T45 | *Eleusine coracana* | Tanzania |  |
| U2 | *Eleusine coracana* | Uganda |  |

**Table S8.** Number of full length and solo LTR-retrotransposons in finger millet, rice and wheat blast genome assemblies

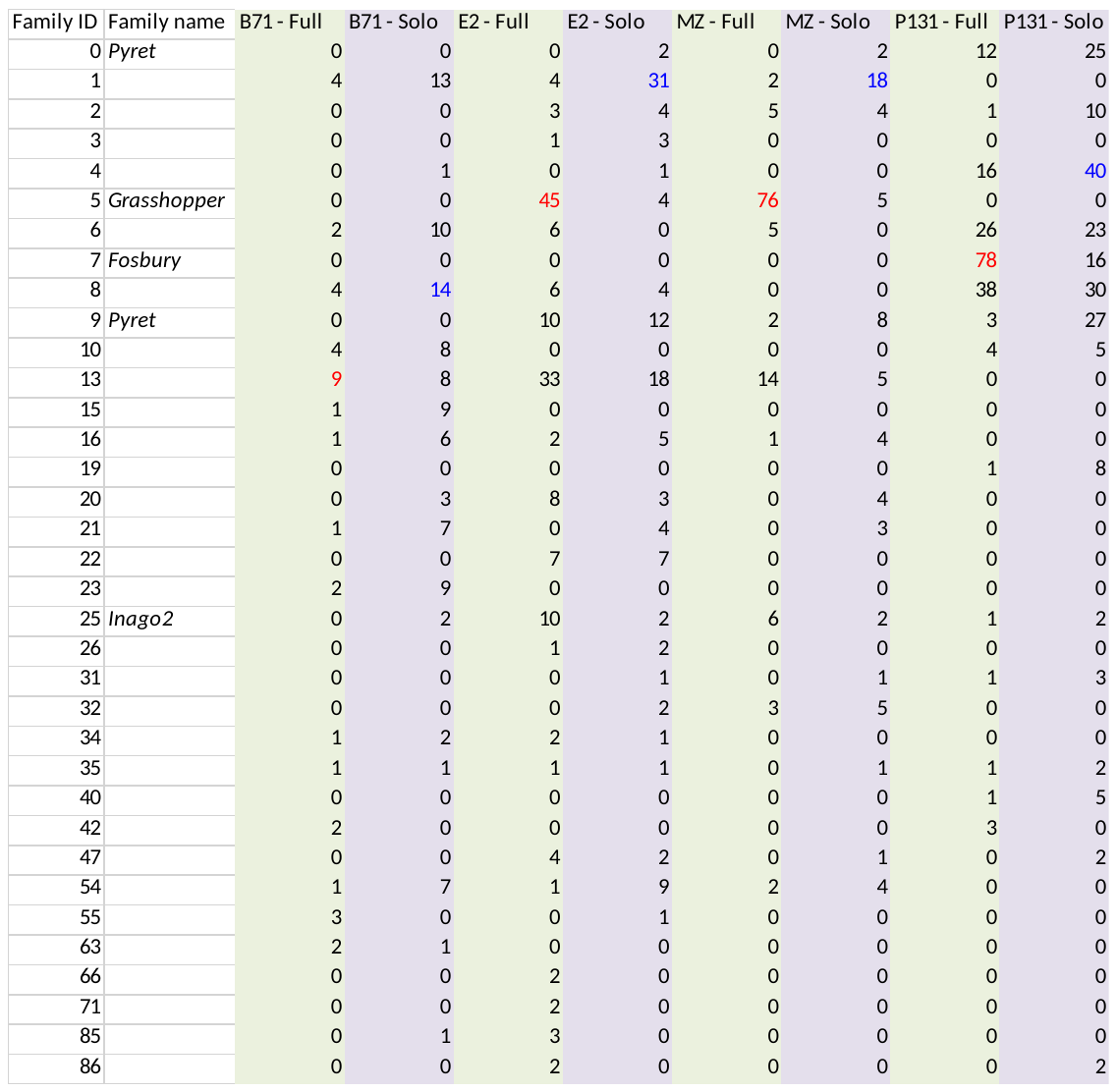

For consistency, repeats were *de novo* annotated in the genome assemblies of B71 (wheat-infecting isolate)^30^, E2 (finger millet-infecting isolate from Ethiopia), MZ (finger millet-infecting isolate from Japan (MZ5-1-6))^31^ and P131 (rice-infecting isolate)^32^

**Table S9.** Estimated average copy numbers in KU and ET populations for seven transposable elements

| TE family | KU strains | ET strains | Admixed strains |
| --- | --- | --- | --- |
| *Family 1* | 68.1 | 36.8 | 42.1 |
| *Family 13* | 35.9 | 31.3 | 30.8 |
| *Fosbury* | 0.5 | 13.8 | 0.2 |
| *Grasshopper* | 4.5 | 31.7 | 37.2 |
| *Inago2* | 11.2 | 17.2 | 17.4 |
| *Pyret* | 17.7 | 39.7 | 31.6 |
| *Pot2* | 10.6 | 121.2 | 97.5 |

**Table S10.** Pairwise median Ks values and divergence ages for blast isolates with different host-specificity

|  | E2  (Finger millet) | MZ5-1-6  (Finger millet) | B71  (Wheat) | P131  (Rice) |
| --- | --- | --- | --- | --- |
| E2  (Finger millet) | - | 0 | 0.00155 | 0.00665 |
| MZ5-1-6  (Finger millet) | 0 | - | 0.00230 | 0.00630 |
| B71  (Wheat) | 37,511 | 55,661 | - | 0.00580 |
| P131  (Rice) | 160,932 | 152,462 | 140,362 | - |

Median Ks values, calculated across 1303 orthologous single copy genes, are given above the diagonal; estimated divergence ages (in years) using a substitution rate of 2.07e^-8^ substitutions /site/year (average of 2.16e^-8^ ^ref33^ and 1.98e^-8^ ^ref34^) are given below the diagonal.

**Table S11: Tukey multiple comparison test tables of frequency of effector and non-effector gene presence, as predicted by blast origin (grouped by country and strain), with gene as random effect**

|  |  |  | Tanzania | Kenya | Uganda | Ethiopia |
| --- | --- | --- | --- | --- | --- | --- |
|  | Country | Population | KU | KU | KU | ET |
| Effector Genes (96) | Tanzania | KU |  |  |  |  |
|  | Kenya | KU | 0.8674 |  |  |  |
|  | Uganda | KU | 0.8168 | 1 |  |  |
|  | Ethiopia | ET | **0.0146** | **0.0004** | **0.0003** |  |
|  | Tanzania | ET | 0.383 | **0.0476** | **0.0353** | 0.6443 |
| Non-Effector Genes (829) | Tanzania | KU |  |  |  |  |
|  | Kenya | KU | 0.6287 |  |  |  |
|  | Uganda | KU | 0.9 | 0.9857 |  |  |
|  | Ethiopia | ET | 0.4498 | **0.0182** | 0.0789 |  |
|  | Tanzania | ET | 0.0565 | **0.0004** | **0.0034** | 0.8511 |

Highlighted cells indicate significant differences. Parenthetical numbers indicate number of genes

**Table S12: Number of expressed genes in *MoE* isolate CKF4046 under different conditions**

| **Condition** | Number of expressed genes^1^ | Number of uniquely expressed genes^1^ |
| --- | --- | --- |
| ≥ 1 of five conditions^2^ | 7870 | N/A^3^ |
| All five conditions^2^ | 5395 | N/A |
| ≥ 1 infection conditions | 7621 | N/A |
| All four infection conditions | 5806 | 411 |
| Axenic 4d | 6445 | 249 |
| TZA1637 (incompatible) – 30 hpi | 6928 | 246 |
| TZA1637 (incompatible) – 48 hpi | 6884 | 45 |
| AAUFM-44 (compatible) – 30 hpi | 6429 | 50 |
| AAUFM-44 (compatible) – 48 hpi | 6779 | 77 |

^1^ Annotated genes with cDNA length ≥200 bp; A gene was considered expressed if the average normalized read count across three replicates was ≥ 5

^2^ Five conditions are (1) axenic growth for 4 days; (2) 30 hpi on finger millet accession AAUFM-44; (3) 48 hpi on AAUFM-44; (4) 30 hpi on TZA1637; (5) 48 hpi on TZA1637

^3^ N/A = Non-applicable

**Table S13.** Number of genes in each of 13 co-expression clusters

| **Cluster** | **Total gene models in cluster** | **Number (percentage) of annotated genes (≥200 bp) in cluster** | **Number (percentage) of predicted effectors (≥200 bp) in cluster** | **Percentage of genes in cluster corresponding to predicted effectors** | **Number (percentage) of upregulated^1^ effectors in cluster @ 30 hpi** | **Number (percentage) of upregulated^1^ effectors in cluster @ 48 hpi** | **Chi-square statistic with Yates correction** | **Chi-square P-value^2^** |
| --- | --- | --- | --- | --- | --- | --- | --- | --- |
| 1 | 257 | 193 (1.9%) | 30 (4.3%) | 15.5% | 0 (0.0%) | 9 (30.0%) | 17.027 | <0.0001 |
| 2 | 1367 | 1004 (9.7%) | 34 (4.8%) | 3.4% | 3 (8.8%) | 1 (2.9%) | 15.151 | (<0.0001) |
| 3 | 317 | 243 (2.3%) | 151 (21.5%) | 62.1% | 28 (18.5%) | 148 (98.0%) | 559.595 | <0.0001 |
| 4 | 940 | 714 (6.9%) | 81 (11.5%) | 11.3% | 9 (11.1%) | 21 (25.9%) | 17.106 | <0.0001 |
| 5 | 6593 | 5992 (57.9%) | 130 (18.5%) | 2.2% | 25 (19.2%) | 9 (6.9%) | 151.739 | (<0.0001) |
| 6 | 321 | 263 (2.5%) | 9 (1.3%) | 3.4% | 3 (33.3%) | 2 (22.2%) | 3.670 | 0.0554 |
| 7 | 1126 | 866 (8.4%) | 46 (6.6%) | 5.3% | 0 (0.0%) | 15 (32.6%) | 2.231 | 0.1352 |
| 8 | 181 | 173 (1.7%) | 32 (4.6%) | 18.5% | 4 (12.5%) | 4 (12.5%) | 26.832 | <0.0001 |
| 9 | 206 | 143 (1.4%) | 19 (2.7%) | 13.3% | 0 (0.0%) | 6 (31.6%) | 6.815 | 0.0090 |
| 10 | 277 | 197 (1.9%) | 23 (3.3%) | 11.7% | 1 (4.3%) | 8 (34.8%) | 5.381 | 0.0204 |
| 11 | 111 | 97 (0.9%) | 26 (3.7%) | 26.8% | 22 (84.6%) | 20 (76.9%) | 41.311 | <0.0001 |
| 12 | 75 | 72 (0.7%) | 6 (0.9%) | 8.3% | 0 (0.0%) | 3 (50.0%) | 0.063 | 0.8013 |
| 13 | 476 | 398 (3.8%) | 115 (16.4%) | 28.9% | 58 (50.4%) | 77 (67.0%) | 190.433 | <0.0001 |
| Across all clusters | 12,247 | 10,355 | 702 | 6.8% | 153 (21.8%) | 323 (46.0%) |  |  |

^1^ Upregulated in the incompatible (infection on TZA1637) compared to the compatible (infection on AAUFM-44) interaction

^2^ Under-representation of effectors in cluster is indicated by P-value in parenthesis

**Table S14. Summary of biochemical gene clusters identified in E2**

| Chromosome | Cluster number | Annotated genes in BGC (start gene -end gene) | BGC size (# genes) | BGC type | # genes more than 4-fold up- or downregulated at 30 hpi | # genes more than 4-fold up- or downregulated at 48 hpi | Four consecutive genes in cluster regulated in same direction |
| --- | --- | --- | --- | --- | --- | --- | --- |
| 1 | 1.01 | EcM01g00000610 - EcM01g00000680 | 8 | Terpene | 0 | 0 | N/A |
| 1 | 1.02 | EcM01g00002920 - EcM01g00003140 | 23 | PKS | 2 | 0 | N/A |
| 1 | 1.03 | EcM01g00004430 - EcM01g00004560 | 14 | NRPS | 1 | 2 | N/A |
| 1 | 1.04 | EcM01g00011020 - EcM01g00011080 | 7 | PKS | 0 | 0 | N/A |
| 1 | 1.05 | EcM01g00012780 - EcM01g00012860 | 10 | Terpene | 0 | 0 | N/A |
| 1 | 1.06 | EcM01g00012910 - EcM01g00013130 | 23 | RiPP | 1 | 0 | N/A |
| 1 | 1.07 | EcM01g00015110 - EcM01g00015190 | 9 | Terpene | 0 | 0 | N/A |
| 1 | 1.08 | EcM01g00015280 - EcM01g00015340 | 7 | Terpene | 1 | 0 | N/A |
| 1 | 1.09 | EcM01g00017740 - EcM01g00017860 | 12 | NRPS | 1 | 4 | No |
| 2 | 2.01 | EcM02g00022140 - EcM02g00022230 | 10 | PKS | 0 | 0 | N/A |
| 2 | 2.02 | EcM02g00024360 - EcM02g00024580 | 23 | RiPP | 0 | 0 | N/A |
| 2 | 2.03 | EcM02g00025990 - EcM02g00026130 | 15 | NRPS | 0 | 0 | N/A |
| 2 | 2.04 | EcM02g00032280 - EcM02g00032350 | 8 | Terpene | 0 | 0 | N/A |
| 2 | 2.05 | EcM02g00033130 - EcM02g00033180 | 6 | Terpene | 0 | 0 | N/A |
| 2 | 2.06 | EcM02g00040730 - EcM02g00040820 | 10 | PKS | 0 | 0 | N/A |
| 2 | 2.07 | EcM02g00043100 - EcM02g00043200 | 10 | Hybrid | 1 | 1 | N/A |
| 2 | 2.08 | EcM02g00043860 - EcM02g00043910 | 6 | NRPS | 0 | 3 | N/A |
| 2 | 2.09 | EcM02g00044220 - EcM02g00044350 | 14 | Hybrid | 3 | 2 | N/A |
| 2 | 2.10 | EcM02g00044370 - EcM02g00044540 | 18 | PKS | 0 | 1 | N/A |
| 2 | 2.11 | EcM02g00045350 - EcMO2g00045510 | 17 | Hybrid | 1 | 0 | N/A |
| 3 | 3.01 | EcMO3g00048800 - EcMO3g00049010 | 22 | RiPP | 2 | 0 | N/A |
| 3 | 3.02 | EcMO3g00050800 - EcMO3g00050950 | 16 | PKS | 2 | 3 | N/A |
| 3 | 3.03 | EcMO3g00065380 - EcMO3g00065560 | 19 | NRPS | 0 | 0 | N/A |
| 3 | 3.04 | EcMO3g00066070 - EcMO3g00066220 | 16 | NRPS | 1 | 4 | No |
| 3 | 3.05 | EcMO3g00066740 - EcMO3g00066900 | 16 | NRPS | 3 | 6 | No |
| 3 | 3.06 | EcMO3g00067640 - EcMO3g00067820 | 19 | NRPS | 2 | 0 | N/A |
| 4 | 4.01 | EcMO4g00070520 - EcMO4g00070620 | 11 | Other | 0 | 0 | N/A |
| 4 | 4.02 | EcMO4g00075140 - EcMO4g00075480 | 35 | Hybrid | 2 | 0 | N/A |
| 4 | 4.03 | EcMO4g00075560 - EcMO4g00075710 | 16 | NRPS | 0 | 1 | N/A |
| 4 | 4.04 | EcMO4g00077510 - EcMO4g00077640 | 14 | NRPS | 1 | 0 | N/A |
| 4 | 4.05 | EcMO4g00081010 - EcMO4g00081050 | 5 | Other | 1 | 0 | N/A |
| 4 | 4.06 | EcMO4g00084520 - EcMO4g00084670 | 16 | PKS | 1 | 2 | N/A |
| 4 | 4.07 | EcMO4g00084930 - EcMO4g00085110 | 19 | PKS | 0 | 4 | No |
| 4 | 4.08 | EcMO4g00085560 - EcMO4g00085720 | 17 | PKS | 2 | 1 | N/A |
| 4 | 4.09 | EcMO4g00086360 - EcMO4g00086490 | 13 | PKS | 1 | 1 | N/A |
| 5 | 5.01 | EcMO5g00088700 - EcMO5g00088800 | 11 | PKS | 6 | 8 | Yes |
| 5 | 5.02 | EcMO5g00089060 - EcMO5g00089140 | 9 | Other | 0 | 0 | N/A |
| 5 | 5.03 | EcMO5g00093350 - EcMO5g00093470 | 13 | PKS | 0 | 0 | N/A |
| 5 | 5.04 | EcMO5g00093570 - EcMO5g00093650 | 9 | Terpene | 0 | 0 | N/A |
| 5 | 5.05 | EcMO5g00094080 - EcMO5g00094130 | 6 | Terpene | 0 | 0 | N/A |
| 5 | 5.06 | EcMO5g00095010 - EcMO5g00095200 | 20 | RiPP | 1 | 0 | N/A |
| 5 | 5.07 | EcMO5g00098900 - EcMO5g00099110 | 22 | PKS | 3 | 0 | N/A |
| 5 | 5.08 | EcMO5g00099410 - EcMO5g00099590 | 19 | NRPS | 1 | 1 | N/A |
| 5 | 5.09 | EcMO5g00100810 - EcMO5g00101030 | 23 | RiPP | 2 | 0 | N/A |
| 5 | 5.10 | EcMO5g00101190 - EcMO5g00101340 | 16 | PKS | 0 | 1 | N/A |
| 5 | 5.11 | EcMO5g00101690 - EcMO5g00101760 | 8 | PKS | 1 | 1 | N/A |
| 5 | 5.12 | EcMO5g00103940 - EcMO5g00104070 | 14 | NRPS | 6 | 0 | Yes |
| 6 | 6.01 | EcMO6g00106230 - EcMO6g00106390 | 17 | NRPS | 0 | 0 | N/A |
| 6 | 6.02 | EcMO6g00106590 - EcMO6g00106630 | 5 | Terpene | 0 | 0 | N/A |
| 6 | 6.03 | EcMO6g00107320 - EcMO6g00107610 | 30 | PKS | 0 | 4 | No |
| 6 | 6.04 | EcMO6g00110500 - EcMO6g00110680 | 19 | PKS | 1 | 1 | N/A |
| 6 | 6.05 | EcMO6g00113920 - EcMO6g00114040 | 13 | RiPP | 1 | 0 | N/A |
| 6 | 6.06 | EcMO6g00114600 - EcMO6g00114730 | 14 | NRPS | 1 | 0 | N/A |
| 6 | 6.07 | EcMO6g00115370 - EcMO6g00115470 | 11 | PKS | 1 | 0 | N/A |
| 6 | 6.08 | EcMO6g00116160 - EcMO6g00116340 | 18 | Hybrid | 1 | 0 | N/A |
| 6 | 6.09 | EcMO6g00116540 - EcMO6g00116640 | 11 | NRPS | 0 | 0 | N/A |
| 6 | 6.10 | EcMO6g00121240 - EcMO6g00121300 | 7 | Terpene | 0 | 0 | N/A |
| 6 | 6.11 | EcMO6g00121370 - EcMO6g00121600 | 24 | Hybrid | 0 | 2 | N/A |
| 6 | 6.12 | EcMO6g00121860 - EcMO6g00122040 | 19 | PKS | 0 | 2 | N/A |
| 6 | 6.13 | EcMO6g00122730 - EcMO6g00123090 | 37 | RiPP | 0 | 4 | No |
| 7 | 7.01 | EcMO7g00125730 - EcMO7g00126000 | 28 | RiPP | 1 | 1 | N/A |
| 7 | 7.02 | EcMO7g00126510 - EcMO7g00126710 | 18 | RiPP | 3 | 2 | N/A |
| 7 | 7.03 | EcMO7g00133470 - EcMO7g00133590 | 13 | PKS | 0 | 0 | N/A |
| 7 | 7.04 | EcMO7g00134540 - EcMO7g00134700 | 17 | NRPS | 0 | 0 | N/A |
| 7 | 7.05 | EcMO7g00137370 - EcMO7g00137640 | 27 | PKS | 2 | 5 | No |
| 7 | 7.06 | EcMO7g00137950 - EcMO7g00138010 | 7 | Hybrid | 0 | 0 | N/A |
| 7 | 7.07 | EcMO7g00138720 - EcMO7g00138790 | 8 | NRPS | 0 | 0 | N/A |

**Table S15: Presence/absence of genes belonging to BGCs 5.01 and 5.12 in *M. oryzae* species with different host specificities**

| Cluster 5.01 | Finger millet | Finger millet | Perennial ryegrass | Wheat | Wheat | Rice | Rice | *Leersia Oryzoides* |
| --- | --- | --- | --- | --- | --- | --- | --- | --- |
|  | E2 | MZ5-1-6 | LpKY97 | Br48 | B71 | ZJG1 | P131 | NI919 |
| EcMO5g00088710 | √ | X | √ | √ | √ | √ | √ | √ |
| EcMO5g00088720 | √ | X | √ | √ | √ | √ | √ | X |
| EcMO5g00088730 | √ | X | √ | √ | √ | √ | X | √ |
| EcMO5g00088740 | √ | X | √ (P)^1^ | √ | √ | √ | X | √ |
| EcMO5g00088750 | √ | √ (Chr7)^2^ | √ | √ | √ | √ | X | X |
| EcMO5g00088760 | √ | X | √ | √ | √ | √ | X | √ |
| EcMO5g00088770 | √ | X | √ | √ | √ | √ (P)^1^ | X | √ (P)^1^ |
| EcMO5g00088780 | √ | X | √ | √ | √ | √ | X | √ |
| EcMO5g00088790 | √ | X | √ | √ | √ | √ | X | √ |

^1^ P indicates a partial gene

^2^ Present on Chr7 rather than the expected Chr5

| Cluster 5.12 | Finger millet | Finger millet | Perennial ryegrass | Wheat | Wheat | Rice | Rice | *Leersia Oryzoides* |
| --- | --- | --- | --- | --- | --- | --- | --- | --- |
|  | E2 | MZ5-1-6 | LpKY97 | Br48 | B71 | ZJG1 | P131 | NI919 |
| EcMO5g00103950 | √ | √ | √ | √ | √ | √ | X | X |
| EcMO5g00103960 | √ | √ | √ | √ | √ | √ | X | X |
| EcMO5g00103970 | √ | √ | √ | √ | X | X | X | X |
| EcMO5g00103980 | √ | √ | √ | √ | √ | √ | X | X |
| EcMO5g00103990 | √ | √ | √ | √ | √ | √ | X | X |
| EcMO5g00104000 | √ | √ | √ | √ | √ | √ | X | X |
